# Clonality and inbreeding amplifies genetic isolation and mate limitation in a rare montane woody plant (*Persoonia hindii*; Proteaceae)

**DOI:** 10.1101/2020.05.25.114231

**Authors:** Collin Ahrens, David Tierney, Paul D. Rymer

## Abstract

Small populations have genetic attributes that make them prone to extinction, including low effective population size (*Ne*), increased levels of inbreeding, and negative impacts from genetic drift. Some small populations are also clonal with low levels of genetic diversity, restricted seed dispersal, and high levels of genetic structure. Together, these attributes make species with small, isolated, clonal populations unlikely to persist under environmental change. We investigated an endangered woody plant species (*Persoonia hindii*) in eastern Australia to answer key questions about genetic differentiation, migration rates, population sizes, size of clones, mating system and *Ne*. We identified 587 single nucleotide polymorphisms. Of the 88 individual stems collected from 15 sites across the entire distribution of *P. hindii*, we identified 30 multi-locus genotypes (MLG). Additional fine-scale genotyping of two sites (49 and 47 stems) detected a dominant genet within each site occupying a minimum area of 20 m^2^. Global population differentiation was high (*F*_*ST*_ 0.22) with very low migration rates (0.048 - 0.064). We identified some population structure with variable site pairwise differentiation (0.015 - 0.32) with no detectable spatial autocorrelation. Species wide inbreeding coefficient was 0.42 (*F*_*IT*_), supporting the direct estimate of 82% selfing. Estimated *Ne* was extremely small (15), indicating that genetic drift may be reducing genetic diversity and increasing genetic load through fixation of deleterious alleles. Clonality and inbreeding combined with negligible gene flow suggests limited adaptive capacity to respond to climate challenges. Genetic rescue, through assisted gene migration and experimental translocations, would enhance the persistence of natural populations.

## Introduction

Population size and geographic isolation strongly impact the genetic diversity of sexually reproducing plant populations (Ellstrand and Elam, 1993; Young *et al.*, 1996). Small population sizes are known to decrease genetic diversity, increase inbreeding events through mate limitation, and increase the power of genetic drift (Keller and Waller, 2002; Willi *et al*. 2006). Additionally, geographic isolation among populations inhibits gene flow, limiting migration of new genetic variants into other populations, and decreases effective population size (Young *et al.*, 1996; Lowe *et al.*, 2005; Aguilar *et al.*, 2008). Population size and genetic diversity interact and effect fitness, particularly through processes such as inbreeding, the accumulation of deleterious mutations through genetic drift (genetic load), and genetic incompatibility due to mating systems (Frankham *et al*., 2002)

Effective population size (*Ne*) determines the strength of random genetic drift within populations (Charlesworth, 2009). Theoretically, an ideal population without demographic and evolutionary processes acting would maintain the same variance of gene frequency in the population through time. Fluctuations of population size, generation times, mating systems, unequal sex ratios, and variances in family size all contribute to reducing *Ne* (Nunney, 1993; Frankham, 1995), as such *Ne* is generally much lower than the number of breeding individuals (Nunney, 1993). Small populations may produce a negative feedback loop where mate limitation results in reproductive failure (allee effect; Gascoigne *et al*. 2009), and increased inbreeding through self-pollination or mating among close relatives (biparental inbreeding) further reducing *Ne*. As *Ne* reduces the strength of random genetic drift increases contributing to the distribution and abundance of diversity within and among populations. For instance, small effective population sizes are more likely to have alleles reach fixation through random sorting processes and as genetic drift accrues randomly fixed alleles in different populations, population differentiation is enhanced. In very small populations the strength of genetic drift makes selection ineffective, such that deleterious alleles become fixed and accumulate within small populations, increasing genetic load (Kirkpatrick and Jarne 2000). Ultimately, allele fixation can have dire consequences coalesced as reduced genetic diversity, decreasing adaptive capacity to perturbations, including climate change (Loeschcke *et al.*, 2013).

Lack of genetic diversity associated with small, isolated populations is exacerbated in clonal plants, even those that exhibit patterns of facultative outcrossing (Fischer *et al.*, 2000; de Vere *et al.*, 2009). Clonal plants with many ramets generally consist of populations with few multilocus genotypes (MLG), further reducing mate availability and increasing the likelihood of inbreeding (Barrett, 2015). Outcrossing events are likely infrequent between clonal populations (Karron *et al.*, 1995), due to pre-zygotic natural barriers between populations such as non-overlapping flowering times and low pollen dispersal. Clonal species may lack evolutionary mechanisms to adapt quickly and accumulate more mutations than outcrossers. This scenario will have two contrasting effects on fitness, increasing fitness due to ramet survivorship or genet persistence (Eriksson and Jerling, 1990); and decreasing fitness due to lower fecundity and decreasing the likelihood of colonisation events (Pan and Price 2002). In addition, clonal plants are potentially unable to repair damaged DNA during cell reproduction, decreasing survivorship (i.e. mutational meltdown and Muller’s ratchet; Muller, 1964; Gabriel *et al.*, 1993; Lynch *et al.*, 1993; Douhovnikoff and Dodd, 2015). Yet there remains questions about the maintenance of genetic diversity within clonal species, as studies on clonal plants report both high and low genetic diversity (Pluess and Stocklin, 2004; Bartlewicz *et al.*, 2015; Wan *et al.*, 2016).

Important developments in genomic analyses are now enabling the characterisation of fine-scale population structure, clonal spread, and demographic patterns through pollen and seed dispersal. It also offers unique insights into the processes maintaining or eroding the genetic diversity in clonal species. In addition to these fundamental insights in evolutionary biology, outputs can improve conservation management by providing a scientific basis for adaptive management strategies. For instance, we are able to target populations/species for *ex situ* conservation (e.g. seed and tissue banking), assisted migration to facilitate genetic rescue (Whiteley *et al.*, 2015), and/or the creation of populations with novel diversity through experimental translocations (Rigg *et al.*, 2017). These outcomes are particularly important for clonal species with small populations, which are prone to genetic impoverishment, and may require interventions for long-term persistence, particularly because of new environmental challenges such as climate change. However, before new management strategies can be applied, knowledge of population structure, genetic diversity, gene flow, effective population size, and mating system, along with their potential interactions, need to be obtained.

*Persoonia hindii* P.H.Weston & L.A.S.Johnson (Proteaceae family) was described in 1997 (Weston and Johnson, 1997) following the first collection in 1989 just 100 km west of Sydney (NSW, Australia) from the upper Blue Mountains. It is a long lived shrubby species that reproduces via rhizomes, but is also known to produce few seeds from sporadic flowering events from January to March (Weston and Johnson, 1997). The species occurs over a 52 km^2^ area in an open, montane, woodland that is dominated by a variety of eucalypt species. *Persoonia hindii* is of significant conservation concern, because it only occurs in 30 sites and is under immediate threat of disturbance from sand mines, forestry, and fire (Weston and Johnson, 1997), and potentially climate change (Beaumont *et al.*, 2019). It is formally listed as an endangered species by the NSW government. Critically, the extent of clonal spread is unknown, the amount of genetic diversity contained within the species is undetermined, nor is the plant mating system or the level of dispersal via pollen and seed known. Predictions for the long-term viability of the species are difficult to ascertain because no studies have been conducted to estimate the *Ne* and species-wide genetic diversity of mature plants and seeds. In this way, *P. hindii* is an ideal species to study the interplay between clonality and sexual reproduction with concurrent studies indicating the species has limited seed production, potentially due to mate limitation with preferential cross pollination between sites potentially increasing seed set (Tierney *et al.*, 2020). Therefore, we focus on filling in gaps in knowledge by quantifying dispersal and population dynamics while estimating species-wide genetic diversity.

We predicted that vegetative reproduction via clonal spread is the dominant mode of reproduction for *Persoonia hindii*. We hypothesise that genetic diversity and *Ne* are low, and gene flow via pollen and seed is limited among populations resulting in high genetic structure. However, for the few flowering events within the species, we predicted to find a predominantly outcrossing mating system (Tierney *et al*., 2020). If these hypotheses are confirmed, then adaptive management strategies may need to be implemented to bolster the genetic diversity of populations to mitigate the impacts from disturbance and climate change.

## Methods

### Sample Collection and Preparation

*Persoonia hindii* is a small, woody species that grows in clonal patches through rhizomatous growth. It has a very narrow distribution that is located west of Sydney in the Newnes Plateau, part of the Blue Mountains World Heritage Area (NSW, Australia). All known populations occur within 10 km of each other at the same elevation (~1000 m above sea level) along consistently flat terrain on sandstone soils supporting dry sclerophyll woodlands (Figure 1a). Detailed ecological surveys have been conducted to document the species distribution and abundance (Ecological Assessments 2014–16). There are at least 30 discrete ‘sites’ (locations with plants separated by at least 20 m from other observed plants) with an unknown number of genets per site. Sampling for genomic analyses was conducted at three different levels to test the core predictions: (1) population structure and seed dispersal; (2) clonality and genet size; and (3) pollen dispersal, mating system, and effective population size.

**Figure 1.**
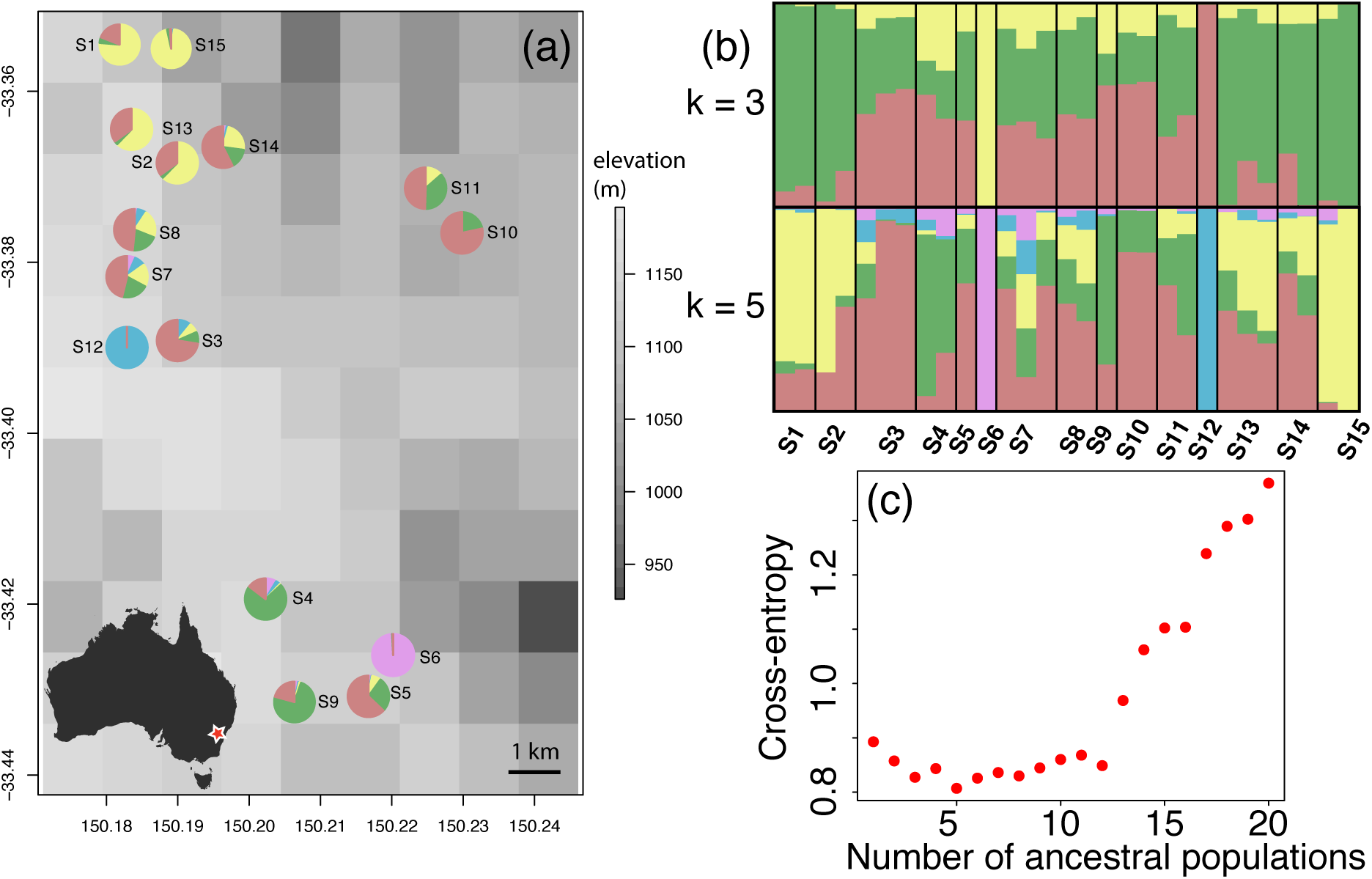
The geographic locations of the 15 sampled sites with elevation as the background and pie charts indicating the assigned proportion of its ancestral genetic clusters at a k-value of 5, Australian map (inset) shows the approximate location of the study area (a). Bar plot from sNMF showing k-values of 3 and 5 (b). Cross-entropy scores for each of the k-values between one and 20 with three and five being the most optimal (i.e. lowest score) (c).

To estimate population structure, seed dispersal, and clonality, we sampled 15 sites covering the entire geographic and climate distribution of the species (Table 1; Figure 1). We collected a leaf sample from six individual stems at each site. Sampling 1-2 plants separated by >4 m in each quarter around a center GPS point marked with a permanent metal stake at each site. This type of sample collection (six individuals per site) has been shown to be adequate for estimating genetic diversity using hundreds of single nucleotide polymorphisms (SNP) (Willing *et al.*, 2012).

**Table 1.**
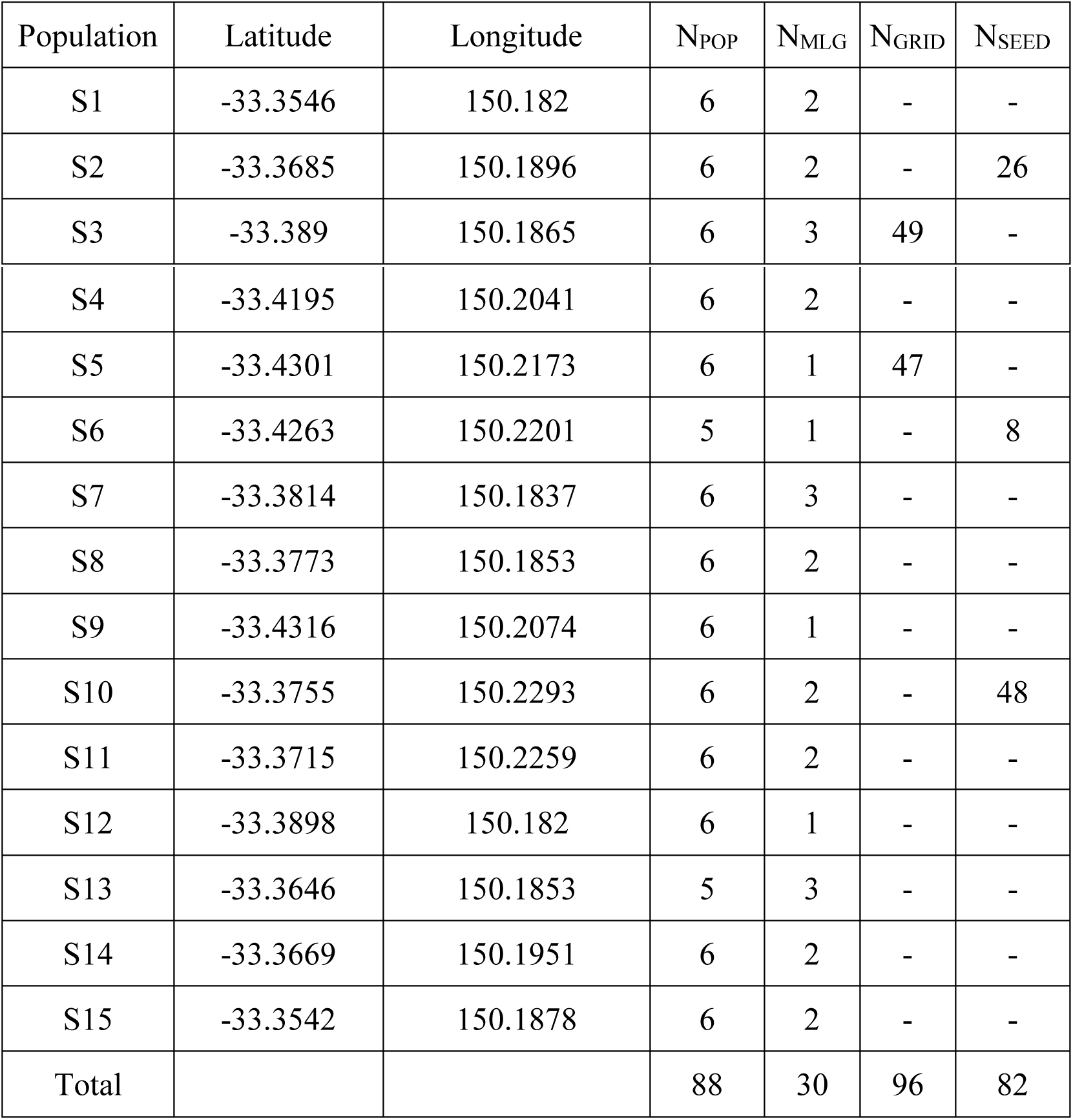
Sampling of *P. hindii* for genomic analyses on the Newnes Plateau (NSW, Australia)… Total sample collection within each population (N_POP_) and the number of unique individuals (N_MLG_) in each population. The number of samples genotyped for the grid analysis (N_GRID_), number of seeds collected (N_SEED_).

To quantify genet size, we exhaustively sampled a 25 m^2^ (5 m by 5 m) plot at two sites (site S3 and S5; Table 1; Figure 1). Both sites had a large number of ramets, however they differed in their fire history (recent/long time since fire, respectively) and seed production (high/low fruit numbers per plant, respectively). Leaf samples were collected from a single stem (where present) from every 0.5 m grid cell within the plot. Of the potential 100 grid cells we sampled stems from 49 and 47 cells for genomic analysis from S3 and S5, respectively. The scale of sampling was based on previous field excavations that traced rhizomes to 1 m in length (David Tierney personal observation).

To quantify the mating system, pollen dispersal, and effective population size in the species we sampled seeds and their parents from three different populations. Sites were limited to those in flower/seed and by the total number of flowers present. In total, three sites could be included (S2 [26 genotyped seeds; 7 mums], S6 [8; 3], and S10 [48; 14]). Seeds were collected by bagging whole ramets after pollination occurred to ensure collection of all fruit after maturation. When the bags were collected from the field after fruit drop, leaf material was collected directly from the mum and labeled with its unique plant ID. In total, 82 genotyped seeds from 24 corresponding mums could be analysed. Knowing the genotype of each seed and its mum, we are able to identify the species’ mating system, infer the probability of a sire (pollen donor), and estimate *Ne*.

Leaf material was collected directly into small seed envelopes and placed into silica gel. The next day, the leaf material was freeze dried using a benchtop freeze dryer (Alpha 1-4 LDplus Laboratory Freeze Dryer, Martin Christ, Harz, Germany). The leaf was then stored in silica gel at room temperature. DNA extraction was performed on 20 mg of dried leaf material for each individual, each sample was placed directly into one well of a 96-well plate designed for DNA extraction using clean tweezers (washed with 95% ethanol solution between the handling of each sample). Collected seeds were placed into a bag with a small amount of silica gel to ensure that the seed would not have fungal contamination. The day before DNA extraction took place, the seed was extracted from the woody endocarp using a vice clamped to a lab bench and the embryo carefully removed using clean tweezers (washed with 95% ethanol solution between the handling of each sample) and placed directly into one well of a 96-well plate designed for DNA extraction. DNA extraction was performed using a modified CTAB procedure at Diversity Array Technologies (University of Canberra, Canberra, Australia).

### Genotyping and SNP calling

DNA samples (400 ng) were sequenced using DArTseq™ protocols (Diversity Arrays Technology Pty Ltd, Canberra Australia) that represent a combination of a double digest complexity reduction method and next generation sequencing platforms (Kilian *et al.*, 2012). A detailed description of the DArTseq™ methodology can be found in Kilian et al. (2012) and Grewe et al. (2015) and has been successfully used for plant population genetic analyses (e.g. Ahrens *et al.*, 2019). Briefly, reduction of the genome was performed using a combination of PstI and HpaII enzymes in digestion/ligation reactions principally as per Kilian et al. (2012) with adapters that include individual varying length barcodes, flowcell attachment sequence, and sequencing primer, similar to Elshire et al. (2011). Raw fastq files were demultiplexed and aligned using Diversity Array Technology’s proprietary bioinformatics pipeline. Poor quality sequences filtered out of the Fastq files with a Phred pass score of 30 (probability of incorrect base is 1 in 1000). Minimum read depth for each individual was set to 6 and average read depth was 82.8 across all SNPs, ensuring call quality for all SNPs. Read lengths consisted of 75 bp and only nucleotide substitutions were considered a SNP for the SNP calling algorithm (proprietary DArTsoft14).

Further SNP filtering was performed in R (R Core Development Team 2018) using custom scripts. For the population genetic and clonal grid analysis, the missing data threshold was set at 35%. However, for the parentage analysis (details below) missing data was set to 10% across all individuals to more accurately assign the probability of sires (pollen donors). Minor allele frequency (MAF) was set to 0.05 to avoid rare alleles that might make population structure analyses unstable. Only one random SNP per 75 bp read was retained to ensure independence and avoid inherent linkage disequilibrium bias. Linkage disequilibrium pruning was done at a threshold of 0.75 using the snpgdsLDpruing function in the SNPRelate package in R (Zheng *et al.*, 2012). We used this threshold because of the clonal nature of the species and its small population size. We called SNPs independently for the population genetic analysis, each of the grids at sites S3 and S5, and the mating system dataset.

### Analysis

#### Identifying multi-locus genotypes

In order to identify the total number of genotypes collected within and among sites to ensure individual genets are not overrepresented in the dataset, we estimated clonality using a ‘clonal threshold’, which was estimated using a series of correlations between known relationships (see below for details). This threshold accounts for inconsistencies in heterozygous calls and sequencing error. The clonal threshold was determined after comparing known clonal material (technical replicates from the same ramet (10 different individuals)), seedlings and mums, and seedlings and other seedlings from the same mum using a correlation analysis (function *cor*) in R and standardising the correlation coefficient between 0 and 1 from a possible maximum and minimum correlation of 1 and −1, respectively. The correlation between clones provided us with a mean and standard error for our expectations of vegetative clones. By using the clonal threshold estimate, the individual data set was reduced to a number of multi-locus genotypes (MLGs). This ensured that each genet was only represented once in each site for the final data set used in the genetic diversity estimates and mating system analysis, but MLGs could occur in different sites, which would account for possible vegetative dispersal events.

#### Population structure and migration estimates

Global population differentiation (F_ST_) and inbreeding (F_IS_) were estimated in the Hierfstat package in R (R Development Team 2018). Global genetic diversity indices were calculated because too few MLGs were present for accurate population level estimates. Shannon’s Diversity Index (H_SHAN_) and expected heterozygosity (H_EXP_) were calculated using poppr in R (Kamvar *et al.*, 2014). In addition, we calculated total heterozygosity (Ht) using Genodive v3.03 (Meirmans and Van Tienderen, 2004).

We used sparse non-negative matrix factorization (sNMF) (Frichot *et al.*, 2014) in the LEA package in R (Frichot and François, 2015) to investigate and visualise individual ancestral cluster assignment. sNMF estimates ancestry coefficients based on sparse non-negative matrix factorisation and least-squares optimisation. The sparse non-negative matrix factorisation is robust to departures from traditional population genetic model assumptions, making this algorithm ideal to use with rare species such as *P. hindii*. We performed sNMF with a k = 1-20, 10 replications per k-value, and 10,000 iterations. Entropy scores for each k-value were compared to choose the optimal number of clusters using the recommendations in the sNMF instruction manual. A consensus for the optimal k-value was created by averaging the results over the 10 replicate runs using CLUMPP v1.1.2 (Jakobsson and Rosenberg, 2007) and drawn using DISTRUCT v1.1 (Rosenberg, 2003; Jakobsson and Rosenberg, 2007).

To see if spatial autocorrelation was a driving force for the genetic diversity patterns that we uncovered, we estimated the coefficient (r) using the *spautocor* function in the package popgenreport (Adamack and Gruber, 2014). The autocorrelation coefficient (r) is calculated for each pairwise genetic distance pairs for all specified distance classes and bootstrapped to estimate 95% null hypothesis confidence intervals for significant spatial autocorrelation (Smouse and Peakall 1999; Smouse et al. 2008). Spatial autocorrelation is present if the r coefficient is outside of the null confidence intervals. We also determined the effective number of migrants using the private alleles method (Barton and Slatkin 1986) in Genepop version 4.2. We predicted migration rates through geographic space using migration surface plots in the MAPS software (Al-Asadi *et al.*, 2019) developed as an extension of EEMS software (Petkova *et al.*, 2016). MAPS uses a coalescent model parameterised with migration rates and population sizes to estimate the migration surface plots. We use the equations provided by Al-Asadi et al (2019) to transform the migration rates to effective spatial diffusion (a dispersal parameter) and population sizes to effective population density. Our data differs from Al-Asadi et al.’s (2019) in that we use a SNP based data set to directly estimate identity by descent, whereas Al-Asadi et al. (2019) uses the number of long segments of haplotype sharing. For both cases, genetic similarities are used to estimate the relatedness. We used all of the scripts provided by the authors of MAPS except for one difference where we used the snpgdsPairIBD function in the SNPRelate package of R to create the identity-by-descent matrix. We also included edges between all sites because the total geographical area encompassed by the study was small and we assumed there was equal probability for dispersal to all sites (also spatial autocorrelation results reinforce this assumption). For visualisation, we bounded the migration rates by the minimum and maximum estimations so we could better identify if populations have heterogeneous migration rates and which populations are a product of greater rates of migration.

#### Genet size (within-site clonality)

Fine-scale within-population clonal structure was determined in two sites with a 5m × 5m grid. For all individuals within the grid, we calculated all pairwise R^2^ values. We used the clonal threshold to determine the number of genets, but also visualised the output as a heatmap for each grid, then mapped the genets on a grid. This provided an estimate for the minimum area occupied by one genet.

#### Pollen dispersal, mating system, and effective population size

Using the third data set with known mothers and seeds, along with potential fathers (37 MLGs from all 15 sites, plus all mothers to allow for selfing), we were interested in identifying the paternal parent across all populations to estimate pollen dispersal distances. This output provides an estimate of possible pollen dispersal within and among populations. We ran COLONY v2.0.6.5 (Jones and Wang, 2010) with known maternal parents and unknown paternal parents. Parameters for COLONY were set to polygamous, dioecious, inbreeding present (includes mothers as possible pollen donors), diploid, and with updating allele frequency. We ran COLONY with 10 runs, length of runs was set to 2 (medium; because we have large amounts of genetic data, and our runs converged during this length), allelic dropout rate was set to 0.1, false allele rate was set to 0.2, probability of dad and mum included in the candidates was set to 0.5 and 1 (respectively), there were no known paternity exclusions, and paternity exclusion threshold was set to 300 (approximately 30% of the data set). We estimated effective population size using the sibship assignment method, detailed by Wang (2009). Briefly, the sibship assignment method theorises that a small population will result in a high proportion of sibs, meaning that there is a high probability that two individuals drawn at random from the same cohort share the same parent(s), and has been shown to be more accurate than other methods (Wang, 2009). We used known MLGs (seeds) to estimate the probability of sibship. This method is well suited to our data set because the high number of markers decreases its bias. We provide results for both random (assuming a negligible deviation from Hardy-Weinberg equilibrium) and non-random mating (deviation from Hardy-Weinberg equilibrium is estimated from the genotype data). While we assume that the non-random mating output is the better estimate here, we show both outputs for transparency. We also estimated effective population size to compare the relative force of genetic drift.

## Results

Diversity Arrays Technology returned 20,385 polymorphic SNPs. Of these, 597 independent polymorphic SNPs were kept after removing uninformative SNPs for the population level analyses. For grids S3 and S5 770 SNPs and 580 SNPs were retained, respectively. For the paternity analyses, 1140 SNPs were maintained in the dataset. The higher number of SNPs for this analysis represents the diversity inherent among parents and seeds.

### Identifying multi-locus genotypes

We used 597 polymorphic SNPs to identify unique multi-locus genotypes (MLG). Correlation between known technical reps was 0.95 +/- 0.03 SD (Table S1), providing an estimate of the genotyping error rate given the expectation of a perfect match. Correlation between mums and their seeds was 0.64 +/- 0.11 SD (min 0.44, max 0.78; Table S2), which relates to the expectation of 50% and 75% similarity for outcrossed and selfed seed, respectively (de Jong and Klinkhamer, 2005). The mean correlation for seeds among all three populations was 0.42 +/- 0.17 SD (full correlation matrix in Table S3). We set the clonal threshold at an R^2^ cut off of 0.90 for calling MLGs (unique genets). This would differentiate clones from siblings in our data set, and aligns well with the theoretical expectation. Using this clonal threshold, we found 30 unique individuals (N_mlg_) from 88 individual stems collected from across the species distribution (i.e. ~6 individuals from 15 populations) (Figure S1a (88 stems) and S1b (30 MLGs)). Three MLGs from different sites (S10 and S11; S13 and S14; S1 and S2) were retained with an R^2^ value above 0.90, representing the only occurrences of MLGs shared among sites, providing evidence for vegetative dispersal between sites (Figure S1 b). The number of unique MLGs per population ranged between one and three individuals with an average of 2 (+/- 0.23 SE) genets. Four populations were reduced to one genotype, eight populations had two genotypes, and three populations had three genotypes (Table 1).

### Genetic diversity, population structure, migration and seed dispersal

Genetic diversity indices among the 30 MLGs across the 15 sites for H_EXP_ was 0.108, Ht was 0.110, and H_SHAN_ was 3.66. Within population heterozygosity (expected heterozygosity) was variable, ranging between 0.03 and 0.08 for populations with more than one MLG present.

Genomic differentiation among sites was F_ST_ = 0.22 using unique MLGs. There was a deficit of heterozygotes with an inbreeding coefficient (F_IS_) of 0.26 and species wide inbreeding coefficient (F_IT_) was 0.42. It is notable that the F statistics with 30 MLGs is in contrast to the 88 individual stems genotyped (F_ST_ 0.40; F_IS_ −0.18), indicating that clonality increases population differentiation and over represents heterozygosity.

Based on the 30 MLGs, pairwise F_ST_ estimates of population differentiation were between 0.01 and 0.35 with 33 of 105 pairwise estimates being > 0.20, highlighting the high levels of genetic differentiation among populations (Table S4). However, there were a few population pairs that had low pairwise F_ST_ estimates (e.g. S9 and S10 = 0.01; S6 and S10 = 0.01).

Population structure using clustering methods describe a biological system with differences between sites (Figure 1) with some detectable variation within sites. The sNMF analysis with a k-value of 3 shows that sites S6 and S12 are the most genetically unique (Figure 1b). However, we show the optimal k-value of 5 based on the cross-entropy plot (Figure 1c). Here, we show that some genomic clusters are shared across sites, providing some evidence of past dispersal between sites. Specifically, high proportions of shared genomic clusters are found between S1, S2, S13, and S15 (yellow; Figure 1b) and located in the northern portion of the distribution (Figure 1a); and between S4 and S9 (green; Figure 1b), located in the southern portion of the distribution. There is also a common genomic cluster found in 10 (>50% assignment; brown; Figure 1b) of the individuals, which might be an ancestral variant.

Spatial autocorrelation was tested with 12 distance bins with a maximum distance of 8km between populations (Figure 2). This analysis revealed that the genetic variation identified at smaller geographic distances is randomly distributed, accept for the 1.98 km bin which was slightly more autocorrelated than expected. As the sites became more distant, the spatial autocorrelation value became lower at the 3.96 km bin (Smouse and Peakall 1999), indicating that these sites at 3.96 km were slightly more differentiated than randomly distributed sites would suggest. All other bins showed random distribution of genetic variation among sites.

**Figure 2.**
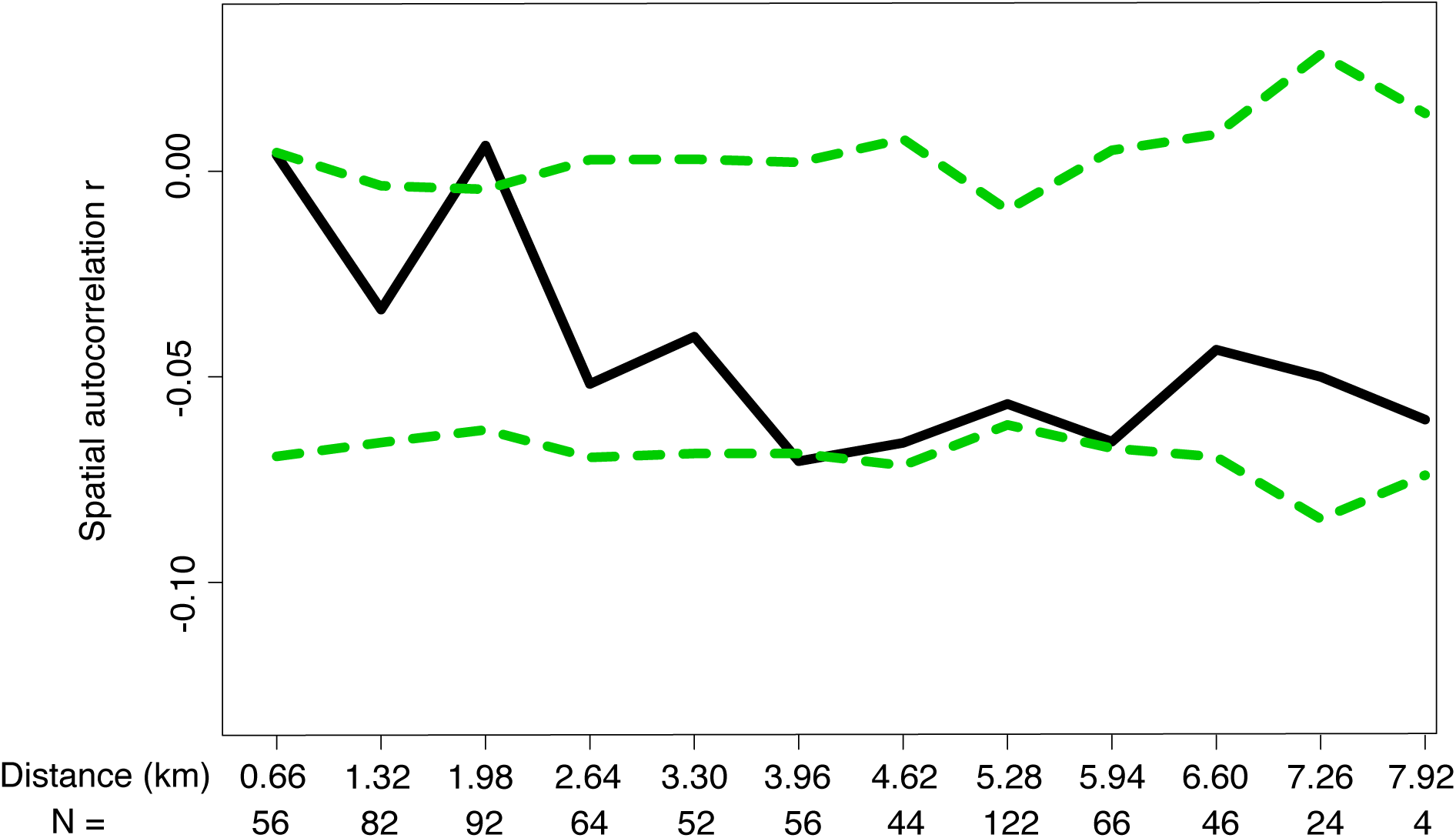
Spatial autocorrelation (r) between all loci across the species distribution shown by the correlogram. The solid line represents the mean estimate of autocorrelation (r) and the 95% null hypothesis confidence regions is shown with the dotted lines. Bins are defined by the geographic distance between individuals (km) and N shows the number of pairwise comparisons within each bin.

Evidence of seed dispersal was found using the frequency of private alleles (0.452). The number of migrants (Nm) was very low (0.058 per mean of 10 individuals) indicating low, but detectable, levels of seed dispersal. This agrees with our correlation analysis, which suggests some dispersal, as we did find shared genets among sites (Figure S1b). For example, one individual from S10 and one from S11 are highly correlated (R^2^ = 0.98); along with one genet from S13 being very similar to one from S14 (R^2^ = 0.94). This information suggests that some seed and/or vegetative dispersal likely occurs. The MAPS method identified migration rates (*m*) between 0.048 and 0.07, showing that migration is very low across the whole distribution with some heterogeneity of migration (Figure 3a). When transforming *m* to dispersal distances, we find the average distance traveled by an individual after one generation is close to 20 meters (typically within sites). Likewise, the posterior mean of the population size was estimated as being between 0.96 and 1.12 across the distribution (Figure 3b), and transformed to population density we find that on average there are 3.47 genets per km^2^.

**Figure 3.**
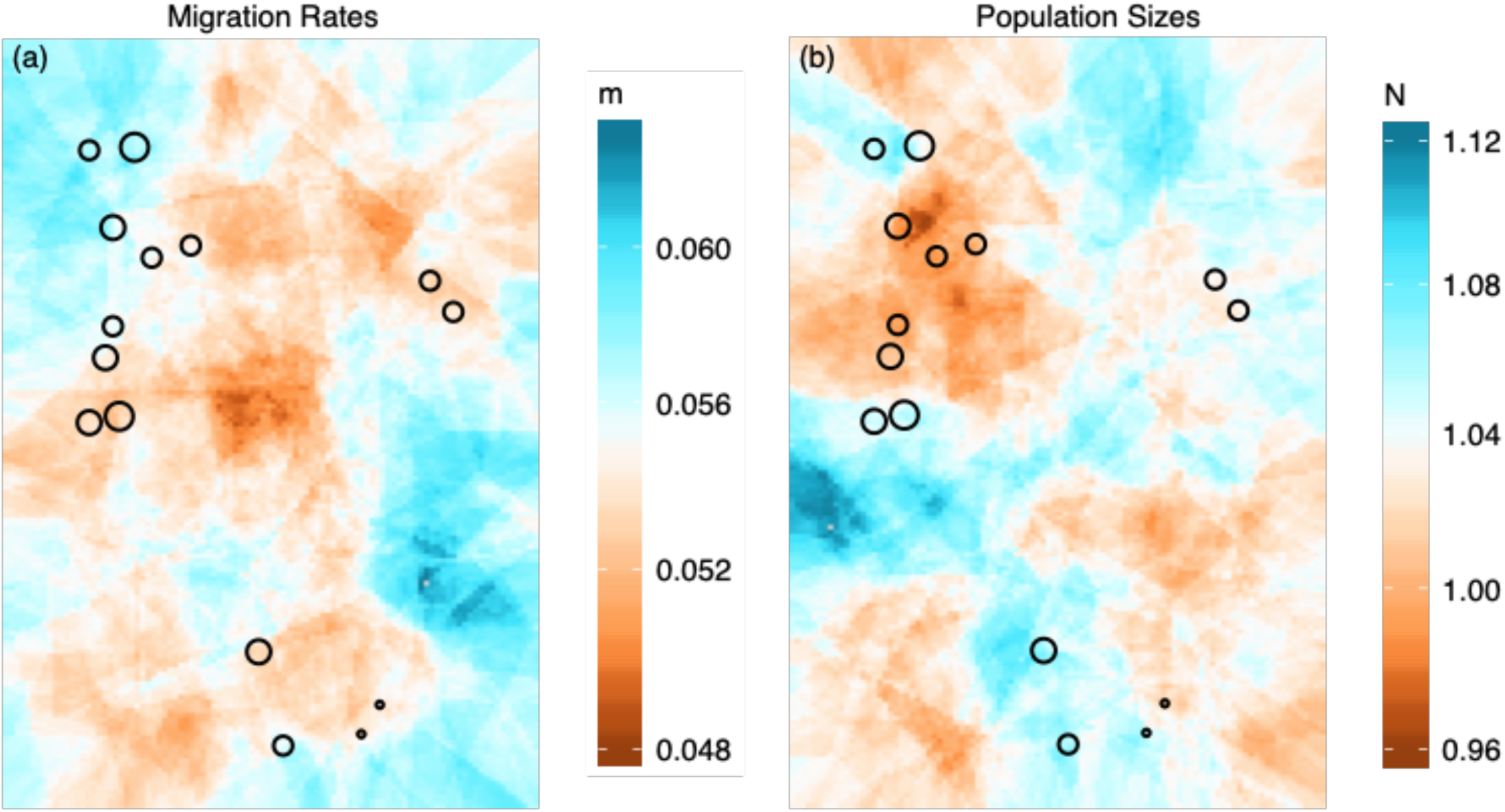
Estimated migration rates and population sizes in geographic space across the Newnes Plateau (NSW, Australia). Color represents (a) posterior mean of migration rates and (b) posterior mean of population size and circle size represents the number of MLGs (from 1 to 4 MLGs).

### Genet size (within site clonality)

We estimated clonality and genet size from 49 stems at S3 with 544 SNPs and 47 stems at S5 with 771 SNPs. The fine-scale spatial structure revealed that one major genet occurs within each site (Figure 4). In addition, both sites had two other genets consisting of multiple ramets with four and two ramets and surrounding genets (12 singletons at site S3 and 5 singletons at site S5) that might be closely related family members (i.e. half and full sibs; Table S5). In Figure 4c & d, we represent these multiple genets as one colour because we are interested in the area occupied by the dominant genet (grey). The lowest R^2^ value for the S3 grid was 0.36, representing the differences between the dominant genet and the other genets and the area occupied by the dominant genet was 5m × 3m (15m^2^). Likewise, the lowest R^2^ value for the S5 grid was 0.39 and the area occupied was 5 m × ~4m (just less than <20m^2^).

**Figure 4.**
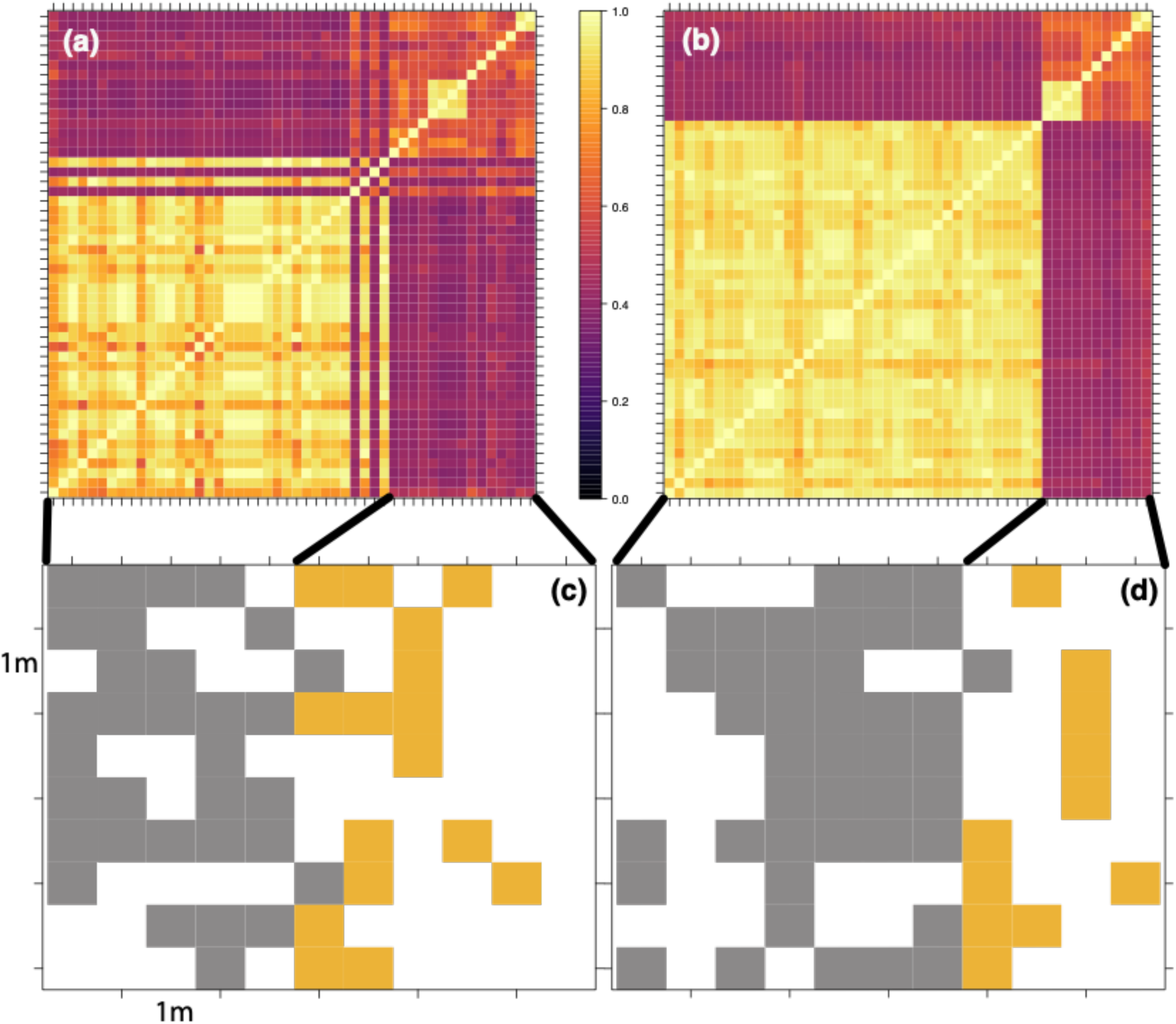
Fine scale population structure within a site. Pairwise correlation (R^2^) values between all stems for each site S3 (a) and S5 (b). While (c) and (d) show the spatial distribution of the stems, highlighting the dominant genet size of each site respectively (grey) with other all other genets as yellow (14 smaller genets in S3; seven smaller genets in S5). Each square is 0.5 m by 0.5 meter making the S3 grid (c) 5 m by 5 m and the S5 grid 5 m by 5.5 m.

### Pollen dispersal, mating system, and effective population size

Of the 82 seeds collected, 13 seeds were dropped due to patchy variant calling, as such 69 were included in these analyses after genomic filtering. Sires were determined for 68 of the 69 offspring with greater than 0.70 probability. Of these 68 offspring with identified sires, 19 were from S2, eight were from S6, and 41 from S10; the one seed without an inferred sire was from S10. Selfing was apparent in all three sites. All of the seeds from S6 and S10 were assigned their mother plant (i.e.100% selfed). Site S2 had the most outcrossing events with 12 seeds being sired from other parental plants (63% outcrossed). Three of the seeds from S2 were assigned to sires from S1, but the correlation between the parents from S1 and S2 had correlation values that were greater than 90%. Across all three sites we found that 18% of the seeds resulted from outcrossing events, while 82% of the seeds were from selfing events. In total, 65 sires were identified and all of the identified sires were represented by only four MLGs based on the clonal threshold (Table S2).

Using the seed to mother relationships, we estimated the species level *Ne* to be 14 (95% CI 8 - 30) assuming random mating and 15 (95% CI 8 - 30) assuming nonrandom mating. However, pollen dispersal analyses suggests that mating within this species is not random and mostly confined to within sites and even within genets.

## Discussion

*Persoonia hindii* is a geographically restricted species under immediate threat from ongoing disturbances that likely interact with its biological characteristics to hinder its persistence. These characteristics include small effective population sizes, low seed dispersal rates, high levels of selfing, and large genets. By elucidating details about its life history, we are able to provide critical information to conserve standing genetic diversity through scientifically informed adaptive management strategies. Here we discuss the fundamental insights into the biology of the system, including clonality, gene flow via pollen and seed, genetic drift, and the capacity to purge deleterious variants. Then we apply our understanding to conservation management.

Rare species typically have low levels of genetic diversity within populations, and genetic diversity for *P. hindii* was generally in-line with other studies of rare species. Genetic differentiation between populations of *P. hindii* was similar to four other outcrossing *Persoonia* species (two common and two rare; Rymer and Ayre 2006), but there were differences between H_EXP_, with *P. hindii* having much lower H_EXP_ than the outcrossing species but Shannon’s *I* was similar. The lower H_EXP_ can be explained through the high clonality found within *P. hindii*, reducing the allele frequencies of the minor alleles compared to the four *Persoonia* obligate outcrossers (Rymer and Ayre 2006). Considering non-related species, H_EXP_ was lower but similar for *P. hindii* (0.10) compared to an endemic species to China *Loropetalum subcordatum* (0.16) (Li *et al.*, 2018), and the difference between the species can be attributed to higher selfing rates in *P. hindii*. In addition, *P. hindii* has similar differentiation to other clonal species such as *Linaria vulgaris* (F_ST_ = 0.33) (Bartlewicz *et al.*, 2016) and *Senecio macrocarpus* (0.33) which displays patterns similar to apomixis (Ahrens and James, 2015). We have also found very different patterns of diversity compared to other rare and clonal species, for example Hewitt et al. (2019) found much higher heterozygosity (H_EXP_ = 0.59) and much lower inbreeding (F_IS_ = 0.024) in a clonal shrub. These differences portray rare, clonal species as having variable genomic diversity with *P. hindii* being on the lower end of the diversity spectrum.

Vegetative clonality within populations seems to be the dominant mode of reproduction, and we show that vegetative clones can grow up to at least 20 m^2^ and likely larger as the clones exceeded the boundaries of our two collection grids. Vegetative reproduction via rhizomes or vegetative propagules is not unique and found in invasive (*Phragmites australis*; Kettenring *et al.*, 2016), common (*Agrostis stolonifera*; Ahrens and Auer, 2012) and rare species (*Comesperma polygaloides*; Ahrens and James, 2016). Therefore, this vegetative mode of reproduction does not preclude rarity, but rather it is likely an important component of a number of other natural processes that make it difficult to persist in and expand to different environments. For instance, the amount of outcrossing is much lower for *P. hindii* compared to these other species, including low migration rates, outcrossing rates, and seed production, creating a system typified by limited genetic diversity with few chances of increasing within population diversity.

The genetic diversity of *Persoonia hindii* is likely strongly linked to mating system, as found in the clonal orchid species *Rutidosis leiolepis* with different levels of mating systems along with different levels of genetic diversity between four populations (Young *et al.*, 2002). Indeed, the high levels of selfing in *P. hindii* with low levels of bi-parental inbreeding due to mate limitation results in decreased genetic diversity. We expected to find higher outcrossing rates in *P. hindii* because concurrent pollination experiments have indicated cross pollination among sites (each genetically distinct) increases seed initiation, relative to within site pollination (Tierney *et al*., 2020). However, our data suggests that outcrossing between different MLGs and sites occurs infrequently and any gene flow that does occur is likely swamped by selfing. In addition, anthropogenic disturbance is known to reduce outcrossing events (Eckert et al. 2010), evolutionary consequences of mate limitation results in increased genetic load leading to decreased progeny vigor and changing demography (Cheptou 2004). These types of consequences are likely impacting *P. hindii*, which already has limited mates, with the threat of continued anthropogenic disturbance there will be fewer outcrossing events. This is notable because outcrossing events provide opportunities of novel combinations of genetic diversity, which are likely critical for long-term persistence of the species.

Low Ne is documented for many species that reproduce mostly via selfing and cloning (Glémin *et al.*, 2019), resulting in high levels of genetic drift. From a theoretical perspective, it is known that genetic drift could have major ramifications on the persistence of a species (Lanfear *et al*., 2014), particularly when 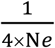 (for *P. hindii*: 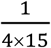 = 0.017) is much greater than the mutation rate (Ellstrand and Elam, 1993). When comparing this (0.017) to mutation rates of other well described plant species (e.g. 3.23 × 10^−10^ in *Chlamydomonas reinhardtii* over 350 generations; Ness *et al.*, 2012). We demonstrate that genetic drift is a stronger agent of genetic change compared to selection in *P. hindii*, further evidence that the species is at risk of extinction. When in context of Mullers ratchet, the increase of genetic load due to genetic drift is a direct threat to the species persistence, because the accumulation of deleterious alleles will impede the ability to adapt to new conditions (Maynard Smith, 1978; Bell, 1982), but a “little bit” of outcrossing may protect this species from genomic decay (Hojsgaard and Hörandl 2015).

Outcrossing rates in *P. hindii* is fundamentally impeded by low migration rates and low population density. Plant migration is important for two main reasons: gene flow mixes genomic variants among populations; and dispersal is required to track changing environments. Gene flow plays a critical role in plant evolution (Ellstrand 2014), it spreads beneficial alleles across populations while decreasing the likelihood of fixation of deleterious alleles (i.e. counteracting genetic drift) and it maintains connectivity between populations and homogenises genetic diversity (Ellstrand 2014). If migration rates and dispersal distances are low, these positive outcomes of gene flow are neutralised. High migration rates would allow for the tracking with climate change (Corlett and Westcott 2013). If a habitat becomes uninhabitable through higher temperatures or reduced rainfall, higher migration rates and dispersal distances could conceivably allow for the species to move to more optimum habitats. For *P. hindii*, both of these outcomes are critical (sharing of variants among populations while counteracting drift and moving to more suitable habitat), however the migration rates and dispersal distances that we have estimated describe a system that has limited exchange of genes among populations or disperse to more suitable habitats. Leaving the species in dire circumstances, relying on isolated, clonal individuals to self for recombination events.

### Conservation and management

*Persoonia hindii* has a very limited distribution confined to a high elevation plateau with predominantly clonal reproduction, small effective population size, high levels of selfing and limited effective dispersal. These characteristics imply that adaptive potential is low, particularly when considering the potential effects of climate change (Willi *et al.*, 2006; Hoffmann *et al.*, 2017; Beaumont *et al*. 2019), indeed patterns of high genetic drift and low gene flow have been suggestive of limited adaptive potential in a fish species (Perrier et al. 2017). *Persoonia hindii* is unlikely to migrate through seed dispersal and occurs in the highest area of the region, indicating that even if it was able to migrate there is no suitable habitat available. As the climate continues to change and local disturbance jeopordises its habitat, there is concern that *P. hindii* will lose sites and diversity, further decreasing its likelihood of persistence. Our results suggest that *P. hindii* may benefit from adaptive management strategies (Holling, 2005).

There are a number of *in situ* and *ex situ* actions that could help protect the species. *In situ* actions include hand pollination events to mimic pollen dispersal, assisted gene migration to mimic seed dispersal, and translocation to new suitable habitats to mimic long(er)-distance migration. *Ex situ* actions include seed collection and banking, propagation and rearing of offspring between important breeding individuals, and experiments to test adaptive capacity. These actions could be trialed in some sites and then applied more broadly if they prove effective. For example, records for the species suggest that some loss has occurred via past clearing for forestry and road disturbance, likely resulting in the loss of genetic diversity (personal observations and unpublished modelling results from David Tierney). While there are some sites with unique diversity, such as S6 and S12, the Ne is so small that protecting all sites are a priority to maintain current diversity represented by the entire distribution. In other words, *any* diversity in a given site and its connection to other sites may be paramount for long-term species viability.

Assisted gene migration for genetic rescue would enrich site-level genetic diversity by moving vegetatively propagated individuals or hand-pollinated seed into established sites or translocated into new sites. The positive impacts of a strategy like this would decrease the rate of site-level genetic drift, increase site-level adaptive potential, and increase recombination rates, which could mitigate contemporary and future impacts. However, there are some possible negative consequences associated with these types of strategies that include increased genetic load and outbreeding depression. We have established that genetic drift is a strong force of shaping genetic diversity in *P. hindii* and assisted migration might have unintended random effects at the site level. For example, through random or selected movement of genotypes, genetic load could increase by chance (Bouzat, 2011), or the introgression of divergent populations could increase heterozygosity rates resulting in decreased fitness of populations, through epistatic interactions or underdominance (Schierup and Christiansen, 1996). However, previous work suggests that this may not be an issue because of the higher rates of seed initiation between divergent individuals (Tierney *et al.*, 2020), although further tests need to confirm this in mature plants. If the benefits outweigh the possible risks of genetic load and outbreeding depression (Frankham *et al*., 2011), then actions such as assisted gene migration, genetic rescue or translocation should be employed to limit the negative consequences associated with genetic drift and low genetic diversity. Specifically, these risky strategies would facilitate outcrossing events between genetically distinct individuals, increasing genetic recombination and heterozygosity, providing tangible benefits for the species (e.g. Ralls *et al.*, 2018). Our research provides empirical evidence to support improved relocation and translocation strategies to promote the long-term persistence of the species.

## Conclusion

Conserving small plant populations is integral to maintaining current biodiversity. Some plants provide more challenging obstacles for development of effective conservation management strategies than others. *Persoonia hindii* is a species with challenging impediments that need to be carefully considered to safeguard the species into the future. To its detriment, it has a very narrow distribution occurring at the top of a mountain, spanning 52 km^2^, within one plant community type. Therefore, the scope to identify suitable habitat in the future outside of its current distribution is limited based on ecology and climate (Beaumont et al. 2019). Further, we use genomics to describe a species that has very low breeding potential with high probability of accumulation of deleterious mutations (i.e. a high genetic load). The dominant mating system elucidated in *P. hindii* suggests that the species is at higher risk of extinction than previously thought with no foreseeable viable options through natural processes to ameliorate these negative attributes. Critically, our results show that *P. hindii* is prone to site-level processes (e.g. high inbreeding, low *Ne*, and low migration rates) that reduce genetic diversity through random events, ultimately leading to severe consequences within sites and the species more broadly. Therefore, we suggest leveraging the knowledge gained in this study to create effective management plans that prioritise actions such as the protection of all sites, the translocation of genotypes, the collection of seeds, and the pollination between independent genotypes. Creating effective monitoring and intervention strategies that may mitigate ongoing and predicted negative impacts from climate change and anthropogenic disturbances, greatly increasing the chances of long-term persistence.

## Acknowledgements

Funding for this research was provided by Conservation and Restoration Science, Science Division, Department of Planning, Industry and Environment, NSW

## Data accessibility

All data used in this study will be made available through DRYAD.

## Supplementary Information

Table S1 – Correlations between known technical replicates.

Table S2 – Correlations between mums and their seeds.

Table S3 – Correlations between all seeds.

Table S4 – Site pairwise FST matrix.

Table S5 – Correlations between all of the sampled ramets within each grid.

Figure S1 – Heat maps displaying the correlation (r^2^) between all (a) 88 individuals collected and (b) the 30 MLGs.

