## Supplementary material for "Clonality and inbreeding amplifies genetic isolation and mate limitation in a rare montane woody plant (*Persoonia hindii*; Proteaceae)": Figure S1

**Supplementary information**

| | Description | Page |
|---|---|---|
| Table S1 | Correlations between known technical replicates. | 2 |
| Table S2 | Correlations between mums and their seeds. | 3 |
| Table S3 | Correlations between all seeds. | 4 |
| Table S4 | Site pairwise FST matrix. | 5 |
| Table S5 | Correlations between all of the sampled ramets within each grid. | 6 |
| Figure S1 | Heat maps displaying the correlation ($r^2$) between all (a) 88 individuals collected and (b) the 30 MLGs. | 7 |

Table S1. Correlation coefficient between technical replicates, indicated by the outlined boxes.

| ID | P10_1_1 | P10_1_2 | P10_2_1 | P10_2_2 | P10_2_3 | P10_3_1 | P10_3_2 | P10_3_3 | P10_5_1 | P10_5_4 | P10_6_1 | P10_6_2 | P10_7_1 | P10_7_2 | P2_2_1 | P2_2_2 | P2_5_1 | P2_5_2 | P2_5_3 | P2_6_1 | P2_6_2 | P6_9_1 | P6_9_2 | P6_9_3 |
|---|---|---|---|---|---|---|---|---|---|---|---|---|---|---|---|---|---|---|---|---|---|---|---|---|
| P10_1_1 | 1.000 | 1.000 | 0.981 | 0.981 | 0.985 | 0.969 | 0.989 | 0.985 | 0.992 | 0.988 | 0.970 | 0.989 | 0.985 | 0.981 | 0.370 | 0.349 | 0.337 | 0.342 | 0.363 | 0.348 | 0.366 | 0.356 | 0.375 | 0.373 |
| P10_1_2 | 1.000 | 1.000 | 0.981 | 0.981 | 0.985 | 0.969 | 0.989 | 0.985 | 0.992 | 0.988 | 0.970 | 0.989 | 0.985 | 0.981 | 0.370 | 0.349 | 0.337 | 0.342 | 0.363 | 0.348 | 0.366 | 0.356 | 0.375 | 0.373 |
| P10_2_1 | 0.981 | 0.981 | 1.000 | 0.969 | 0.965 | 0.965 | 0.970 | 0.965 | 0.981 | 0.969 | 0.950 | 0.969 | 0.958 | 0.961 | 0.358 | 0.337 | 0.333 | 0.329 | 0.351 | 0.335 | 0.354 | 0.339 | 0.372 | 0.356 |
| P10_2_2 | 0.981 | 0.981 | 0.969 | 1.000 | 0.965 | 0.957 | 0.970 | 0.965 | 0.973 | 0.969 | 0.958 | 0.977 | 0.965 | 0.961 | 0.379 | 0.358 | 0.347 | 0.351 | 0.372 | 0.357 | 0.375 | 0.369 | 0.394 | 0.386 |
| P10_2_3 | 0.985 | 0.985 | 0.965 | 0.965 | 1.000 | 0.969 | 0.981 | 0.969 | 0.984 | 0.981 | 0.962 | 0.981 | 0.977 | 0.981 | 0.373 | 0.351 | 0.340 | 0.344 | 0.366 | 0.350 | 0.369 | 0.373 | 0.385 | 0.383 |
| P10_3_1 | 0.969 | 0.969 | 0.965 | 0.957 | 0.969 | 1.000 | 0.973 | 0.954 | 0.977 | 0.973 | 0.954 | 0.973 | 0.969 | 0.980 | 0.379 | 0.358 | 0.354 | 0.350 | 0.372 | 0.357 | 0.375 | 0.367 | 0.379 | 0.370 |
| P10_3_2 | 0.989 | 0.989 | 0.970 | 0.970 | 0.981 | 0.973 | 1.000 | 0.974 | 0.989 | 0.985 | 0.966 | 0.985 | 0.981 | 0.985 | 0.372 | 0.351 | 0.339 | 0.336 | 0.364 | 0.349 | 0.367 | 0.363 | 0.375 | 0.373 |
| P10_3_3 | 0.985 | 0.985 | 0.965 | 0.965 | 0.969 | 0.954 | 0.974 | 1.000 | 0.977 | 0.973 | 0.962 | 0.973 | 0.977 | 0.973 | 0.376 | 0.355 | 0.344 | 0.348 | 0.369 | 0.354 | 0.372 | 0.357 | 0.376 | 0.374 |
| P10_5_1 | 0.992 | 0.992 | 0.981 | 0.973 | 0.984 | 0.977 | 0.989 | 0.977 | 1.000 | 0.988 | 0.977 | 0.988 | 0.984 | 0.988 | 0.364 | 0.342 | 0.331 | 0.335 | 0.357 | 0.341 | 0.359 | 0.353 | 0.365 | 0.363 |
| P10_5_4 | 0.988 | 0.988 | 0.969 | 0.969 | 0.981 | 0.973 | 0.985 | 0.973 | 0.988 | 1.000 | 0.966 | 0.985 | 0.981 | 0.985 | 0.377 | 0.356 | 0.345 | 0.349 | 0.370 | 0.355 | 0.373 | 0.355 | 0.366 | 0.364 |
| P10_6_1 | 0.970 | 0.970 | 0.950 | 0.958 | 0.962 | 0.954 | 0.966 | 0.962 | 0.977 | 0.966 | 1.000 | 0.965 | 0.977 | 0.973 | 0.334 | 0.313 | 0.302 | 0.306 | 0.327 | 0.312 | 0.330 | 0.349 | 0.353 | 0.359 |
| P10_6_2 | 0.989 | 0.989 | 0.969 | 0.977 | 0.981 | 0.973 | 0.985 | 0.973 | 0.988 | 0.985 | 0.965 | 1.000 | 0.980 | 0.984 | 0.382 | 0.361 | 0.349 | 0.353 | 0.375 | 0.360 | 0.378 | 0.372 | 0.383 | 0.381 |
| P10_7_1 | 0.985 | 0.985 | 0.958 | 0.965 | 0.977 | 0.969 | 0.981 | 0.977 | 0.984 | 0.981 | 0.977 | 0.980 | 1.000 | 0.988 | 0.371 | 0.350 | 0.339 | 0.343 | 0.364 | 0.349 | 0.367 | 0.360 | 0.365 | 0.370 |
| P10_7_2 | 0.981 | 0.981 | 0.961 | 0.961 | 0.981 | 0.980 | 0.985 | 0.973 | 0.988 | 0.985 | 0.973 | 0.984 | 0.988 | 1.000 | 0.382 | 0.361 | 0.349 | 0.353 | 0.375 | 0.360 | 0.378 | 0.372 | 0.376 | 0.374 |
| P2_2_1 | 0.370 | 0.370 | 0.358 | 0.379 | 0.373 | 0.379 | 0.372 | 0.376 | 0.364 | 0.377 | 0.334 | 0.382 | 0.371 | 0.382 | 1.000 | 0.970 | 0.959 | 0.966 | 0.966 | 0.966 | 0.970 | 0.395 | 0.406 | 0.376 |
| P2_2_2 | 0.349 | 0.349 | 0.337 | 0.358 | 0.351 | 0.358 | 0.351 | 0.355 | 0.342 | 0.356 | 0.313 | 0.361 | 0.350 | 0.361 | 0.970 | 1.000 | 0.974 | 0.966 | 0.974 | 0.981 | 0.985 | 0.408 | 0.404 | 0.381 |
| P2_5_1 | 0.337 | 0.337 | 0.333 | 0.347 | 0.340 | 0.354 | 0.339 | 0.344 | 0.331 | 0.345 | 0.302 | 0.349 | 0.339 | 0.349 | 0.959 | 0.974 | 1.000 | 0.955 | 0.963 | 0.978 | 0.974 | 0.385 | 0.383 | 0.359 |
| P2_5_2 | 0.342 | 0.342 | 0.329 | 0.351 | 0.344 | 0.350 | 0.336 | 0.348 | 0.335 | 0.349 | 0.306 | 0.353 | 0.343 | 0.353 | 0.966 | 0.966 | 0.955 | 1.000 | 0.955 | 0.963 | 0.966 | 0.388 | 0.385 | 0.362 |
| P2_5_3 | 0.363 | 0.363 | 0.351 | 0.372 | 0.366 | 0.372 | 0.364 | 0.369 | 0.357 | 0.370 | 0.327 | 0.375 | 0.364 | 0.375 | 0.966 | 0.974 | 0.963 | 0.955 | 1.000 | 0.977 | 0.981 | 0.402 | 0.406 | 0.382 |
| P2_6_1 | 0.348 | 0.348 | 0.335 | 0.357 | 0.350 | 0.357 | 0.349 | 0.354 | 0.341 | 0.355 | 0.312 | 0.360 | 0.349 | 0.360 | 0.966 | 0.981 | 0.978 | 0.963 | 0.977 | 1.000 | 0.989 | 0.381 | 0.378 | 0.355 |
| P2_6_2 | 0.366 | 0.366 | 0.354 | 0.375 | 0.369 | 0.375 | 0.367 | 0.372 | 0.359 | 0.373 | 0.330 | 0.378 | 0.367 | 0.378 | 0.970 | 0.985 | 0.974 | 0.966 | 0.981 | 0.989 | 1.000 | 0.399 | 0.396 | 0.373 |
| P6_9_1 | 0.356 | 0.356 | 0.339 | 0.369 | 0.373 | 0.367 | 0.363 | 0.357 | 0.353 | 0.355 | 0.349 | 0.372 | 0.360 | 0.372 | 0.395 | 0.408 | 0.385 | 0.388 | 0.402 | 0.381 | 0.399 | 1.000 | 0.945 | 0.961 |
| P6_9_2 | 0.375 | 0.375 | 0.372 | 0.394 | 0.385 | 0.379 | 0.375 | 0.376 | 0.365 | 0.366 | 0.353 | 0.383 | 0.365 | 0.376 | 0.406 | 0.404 | 0.383 | 0.385 | 0.406 | 0.378 | 0.396 | 0.945 | 1.000 | 0.934 |
| P6_9_3 | 0.373 | 0.373 | 0.356 | 0.386 | 0.383 | 0.370 | 0.373 | 0.374 | 0.363 | 0.364 | 0.359 | 0.381 | 0.370 | 0.374 | 0.376 | 0.381 | 0.359 | 0.362 | 0.382 | 0.355 | 0.373 | 0.961 | 0.934 | 1.000 |

Table S2. Correlation coefficients between seeds and mums. Boxes represent the mums of seeds.

| SITE 2 | Mums | | | | | | |
|---|---|---|---|---|---|---|---|
| **Seeds** | **P2_2_1** | **P2_2_2** | **P2_5_1** | **P2_5_2** | **P2_5_3** | **P2_6_1** | **P2_6_2** |
| X11.2.2.1.1 | 0.7481322 | 0.7184823 | 0.7223035 | 0.7103695 | 0.7019394 | 0.7103695 | 0.7184823 |
| X12.2.2.1.2 | 0.6135579 | 0.6135579 | 0.6215469 | 0.6019163 | 0.6248276 | 0.6019163 | 0.6135579 |
| X13.2.2.1.3 | 0.642118 | 0.6131891 | 0.6446079 | 0.6323612 | 0.6557251 | 0.6323612 | 0.642118 |
| X14.2.2.2.4 | 0.7562177 | 0.7562177 | 0.7272727 | 0.7494916 | 0.7416919 | 0.7494916 | 0.7267683 |
| X15.2.2.2.5 | 0.7965034 | 0.7649364 | 0.800143 | 0.7888531 | 0.810926 | 0.7888531 | 0.7965034 |
| X17.2.2.2.7 | 0.4732913 | 0.4412037 | 0.4510375 | 0.4880893 | 0.4525557 | 0.4880893 | 0.4412037 |
| X18.2.2.2.8 | 0.697319 | 0.697319 | 0.6691739 | 0.6583812 | 0.7120242 | 0.6583812 | 0.6678696 |
| X19.2.2.2.9 | 0.7325526 | 0.7325526 | 0.7347394 | 0.7245655 | 0.7468744 | 0.7245655 | 0.7325526 |
| X28.2.5.1.8 | 0.6510848 | 0.6510848 | 0.6617782 | 0.6714418 | 0.6284778 | 0.6714418 | 0.6510848 |
| X30.2.5.1.10 | 0.643692 | 0.6106064 | 0.648354 | 0.6638178 | 0.6249535 | 0.6638178 | 0.643692 |
| X21.2.5.2.1 | 0.7562177 | 0.7267683 | 0.7563221 | 0.7494916 | 0.7416919 | 0.7494916 | 0.7562177 |
| X23.2.5.2.3 | 0.7704561 | 0.7420896 | 0.7410051 | 0.7360384 | 0.7572864 | 0.7360384 | 0.7420896 |
| X25.2.5.3.5 | 0.7374462 | 0.7092956 | 0.7105715 | 0.7024418 | 0.7231627 | 0.7024418 | 0.7374462 |
| X31.2.6.1.1 | 0.6201917 | 0.6201917 | 0.5955147 | 0.6067686 | 0.601034 | 0.6067686 | 0.5883724 |
| X32.2.6.1.2 | 0.7316702 | 0.7034223 | 0.7580562 | 0.7545468 | 0.718546 | 0.7545468 | 0.7316702 |
| X33.2.6.1.3 | 0.8163699 | 0.7608716 | 0.8145696 | 0.8132779 | 0.8316567 | 0.8132779 | 0.8163699 |
| X35.2.6.1.5 | 0.7029298 | 0.6713628 | 0.6789261 | 0.6597998 | 0.715523 | 0.6597998 | 0.6713628 |
| X36.2.6.2.6 | 0.782482 | 0.7544737 | 0.7822154 | 0.7780633 | 0.7688841 | 0.7780633 | 0.782482 |
| X38.2.6.2.8 | 0.792098 | 0.7629346 | 0.790587 | 0.7867887 | 0.7785583 | 0.7867887 | 0.792098 |

| SITE 6 | Mums | | |
|---|---|---|---|
| **Seeds** | **P6_9_1** | **P6_9_2** | **P6_9_3** |
| X1.6.9.1.1 | 0.8080548 | 0.7843155 | 0.8248229 |
| X2.6.9.1.2 | 0.9159502 | 0.9175526 | 0.9056005 |
| X4.6.9.1.4 | 0.6449718 | 0.6188091 | 0.6648226 |
| X6.6.9.2.6 | 0.6768065 | 0.6701143 | 0.6750206 |
| X7.6.9.2.7 | 0.6082724 | 0.5939991 | 0.6111944 |
| X8.6.9.3.8 | 0.593679 | 0.5879693 | 0.6179963 |
| X9.6.9.3.9 | 0.8587992 | 0.8541273 | 0.8516215 |
| X10.6.9.3.10 | 0.5677475 | 0.5327313 | 0.5927652 |

| SITE 10 | Mums | | | | | | | | | | | | | |
|---|---|---|---|---|---|---|---|---|---|---|---|---|---|---|
| **Seeds** | **P10_1_1** | **P10_1_2** | **P10_2_1** | **P10_2_2** | **P10_2_3** | **P10_3_1** | **P10_3_2** | **P10_3_3** | **P10_5_1** | **P10_5_4** | **P10_6_1** | **P10_6_2** | **P10_7_1** | **P10_7_2** |
| X39.10.1.1.1 | 0.6526973 | 0.6526973 | 0.6389713 | 0.667074 | 0.6258441 | 0.6389713 | 0.6526973 | 0.6547086 | 0.6526973 | 0.6389713 | 0.6825493 | 0.6526973 | 0.6526973 | 0.6821598 |
| X40.10.1.1.2 | 0.6677169 | 0.6677169 | 0.6883991 | 0.6468693 | 0.6359603 | 0.6883991 | 0.6677169 | 0.6433529 | 0.6677169 | 0.6515088 | 0.6588586 | 0.6677169 | 0.6677169 | 0.6640906 |
| X42.10.1.1.4 | 0.7570911 | 0.7570911 | 0.7401078 | 0.7748547 | 0.7238421 | 0.7401078 | 0.7570911 | 0.734684 | 0.7570911 | 0.7401078 | 0.7456732 | 0.7570911 | 0.7570911 | 0.7548304 |
| X45.10.1.2.8 | 0.6816091 | 0.6816091 | 0.6667576 | 0.6163233 | 0.6525426 | 0.6667576 | 0.6816091 | 0.6475865 | 0.6816091 | 0.6667576 | 0.6320048 | 0.6816091 | 0.6816091 | 0.6315789 |
| X46.10.1.2.7 | 0.6440322 | 0.6440322 | 0.6283432 | 0.6222679 | 0.6132917 | 0.6283432 | 0.6440322 | 0.656355 | 0.6440322 | 0.6283432 | 0.635543 | 0.6440322 | 0.6440322 | 0.6389149 |
| X47.10.1.2.9 | 0.7500068 | 0.7500068 | 0.7336052 | 0.7282037 | 0.7179054 | 0.7336052 | 0.7500068 | 0.6840503 | 0.7500068 | 0.7336052 | 0.7007791 | 0.7500068 | 0.7500068 | 0.7062274 |
| X48.10.1.2.10 | 0.7046094 | 0.7046094 | 0.6892363 | 0.680798 | 0.6745217 | 0.6892363 | 0.7046094 | 0.6733595 | 0.7046094 | 0.7282037 | 0.6935563 | 0.7046094 | 0.7046094 | 0.6971526 |
| X50.10.2.1.2 | 0.5863533 | 0.5863533 | 0.572334 | 0.6009975 | 0.5588894 | 0.572334 | 0.5863533 | 0.5914298 | 0.5863533 | 0.572334 | 0.5783635 | 0.5863533 | 0.5863533 | 0.6163233 |
| X51.10.2.1.3 | 0.4038463 | 0.4038463 | 0.3916427 | 0.4165414 | 0.3798903 | 0.3916427 | 0.4038463 | 0.3940095 | 0.4038463 | 0.3916427 | 0.4008542 | 0.4038463 | 0.4038463 | 0.3808559 |
| X52.10.2.1.4 | 0.6570176 | 0.6570176 | 0.6392911 | 0.6410727 | 0.6222523 | 0.6392911 | 0.6570176 | 0.608956 | 0.6570176 | 0.6392911 | 0.6500532 | 0.6570176 | 0.6570176 | 0.6250573 |
| X53.10.2.2.5 | 0.3982242 | 0.3982242 | 0.3867659 | 0.4101534 | 0.3757399 | 0.3867659 | 0.3982242 | 0.3878186 | 0.3982242 | 0.3867659 | 0.3947068 | 0.3982242 | 0.3982242 | 0.3754544 |
| X54.10.2.3.6 | 0.6271301 | 0.6271301 | 0.6115727 | 0.5976187 | 0.5966422 | 0.6115727 | 0.6271301 | 0.5372354 | 0.6271301 | 0.6115727 | 0.575112 | 0.6271301 | 0.6271301 | 0.5676715 |
| X55.10.2.3.7 | 0.5924505 | 0.5924505 | 0.5763664 | 0.609212 | 0.5609047 | 0.5763664 | 0.5924505 | 0.6450229 | 0.5924505 | 0.5763664 | 0.5862686 | 0.5924505 | 0.5924505 | 0.6267128 |
| X56.10.2.3.8 | 0.7006262 | 0.7006262 | 0.6853215 | 0.6786753 | 0.670672 | 0.6853215 | 0.7006262 | 0.7120971 | 0.7006262 | 0.6853215 | 0.6896539 | 0.7006262 | 0.7006262 | 0.6949789 |
| X57.10.2.3.9 | 0.5529027 | 0.5529027 | 0.5400201 | 0.5221624 | 0.5276721 | 0.5400201 | 0.5529027 | 0.5044878 | 0.5529027 | 0.5400201 | 0.5024974 | 0.5529027 | 0.5529027 | 0.4909161 |
| X58.10.2.3.10 | 0.6785009 | 0.6785009 | 0.661596 | 0.6573359 | 0.6453708 | 0.661596 | 0.6785009 | 0.6143954 | 0.6785009 | 0.661596 | 0.6325802 | 0.6785009 | 0.6785009 | 0.6359769 |
| X59.10.3.1.1 | 0.6605001 | 0.6605001 | 0.6437851 | 0.6342032 | 0.6277374 | 0.6437851 | 0.6605001 | 0.6254644 | 0.6605001 | 0.6437851 | 0.6103186 | 0.6605001 | 0.6605001 | 0.6075706 |
| X60.10.3.1.2 | 0.5783399 | 0.5783399 | 0.5648946 | 0.5923926 | 0.5520077 | 0.5648946 | 0.5783399 | 0.6225464 | 0.5783399 | 0.5648946 | 0.5700827 | 0.5783399 | 0.5783399 | 0.6071074 |
| X61.10.3.2.3 | 0.681899 | 0.681899 | 0.6658418 | 0.6636337 | 0.6504476 | 0.6658418 | 0.681899 | 0.698687 | 0.681899 | 0.6658418 | 0.6386408 | 0.681899 | 0.681899 | 0.6807462 |
| X63.10.3.2.5 | 0.6720532 | 0.6720532 | 0.6565907 | 0.6525816 | 0.6417737 | 0.6565907 | 0.6720532 | 0.6863266 | 0.6720532 | 0.6565907 | 0.6965925 | 0.6720532 | 0.6720532 | 0.6690517 |
| X66.10.3.3.8 | 0.6646159 | 0.6646159 | 0.6476213 | 0.6439034 | 0.631302 | 0.6476213 | 0.6646159 | 0.6413497 | 0.6646159 | 0.6476213 | 0.6566478 | 0.6646159 | 0.6646159 | 0.6619391 |
| X67.10.5.1.1 | 0.4218598 | 0.4218598 | 0.4081427 | 0.4361136 | 0.3949178 | 0.4081427 | 0.4218598 | 0.3859415 | 0.4218598 | 0.4081427 | 0.3819644 | 0.4218598 | 0.4218598 | 0.411247 |
| X69.10.5.1.3 | 0.6346698 | 0.6346698 | 0.6185072 | 0.6071729 | 0.6029881 | 0.6185072 | 0.6346698 | 0.596535 | 0.6346698 | 0.6185072 | 0.5843064 | 0.6346698 | 0.6346698 | 0.5792961 |
| X72.10.5.4.6 | 0.4528242 | 0.4528242 | 0.4415909 | 0.4645499 | 0.4308101 | 0.4415909 | 0.4528242 | 0.4387891 | 0.4528242 | 0.4415909 | 0.4470547 | 0.4528242 | 0.4528242 | 0.426622 |
| X73.10.5.4.7 | 0.7485761 | 0.7485761 | 0.7310399 | 0.7669015 | 0.7142295 | 0.7310399 | 0.7485761 | 0.735836 | 0.7485761 | 0.7310399 | 0.7710036 | 0.7485761 | 0.7485761 | 0.7513708 |
| X74.10.5.4.8 | 0.5291841 | 0.5291841 | 0.5167874 | 0.4936794 | 0.5049038 | 0.5167874 | 0.5291841 | 0.5201779 | 0.5291841 | 0.5167874 | 0.5217215 | 0.5291841 | 0.5291841 | 0.5066177 |
| X75.10.6.1.1 | 0.6440322 | 0.6440322 | 0.6283432 | 0.6604146 | 0.6132917 | 0.6283432 | 0.6440322 | 0.656355 | 0.6440322 | 0.6283432 | 0.635543 | 0.6440322 | 0.6440322 | 0.6389149 |
| X76.10.6.1.2 | 0.5197344 | 0.5197344 | 0.5045333 | 0.535556 | 0.4899018 | 0.5045333 | 0.5197344 | 0.4855096 | 0.5197344 | 0.5045333 | 0.5153866 | 0.5197344 | 0.5197344 | 0.5107246 |
| X78.10.6.1.4 | 0.5819315 | 0.5819315 | 0.5668916 | 0.551868 | 0.5524464 | 0.5668916 | 0.5819315 | 0.490264 | 0.5819315 | 0.5668916 | 0.5310843 | 0.5819315 | 0.5819315 | 0.521331 |
| X79.10.6.1.5 | 0.816987 | 0.816987 | 0.7990699 | 0.8357362 | 0.7819184 | 0.7990699 | 0.816987 | 0.7984778 | 0.816987 | 0.7990699 | 0.8042618 | 0.816987 | 0.816987 | 0.8171321 |
| X80.10.6.2.6 | 0.7229966 | 0.7229966 | 0.7071857 | 0.7395429 | 0.6920513 | 0.7071857 | 0.7229966 | 0.7751145 | 0.7229966 | 0.7071857 | 0.7116912 | 0.7229966 | 0.7229966 | 0.7568906 |
| X81.10.6.2.7 | 0.691224 | 0.691224 | 0.6745 | 0.7086896 | 0.6584577 | 0.6745 | 0.691224 | 0.7077017 | 0.691224 | 0.6745 | 0.6819999 | 0.691224 | 0.691224 | 0.7269639 |
| X82.10.6.2.8 | 0.6495132 | 0.6495132 | 0.6345153 | 0.6651907 | 0.6201425 | 0.6345153 | 0.6495132 | 0.6988379 | 0.6495132 | 0.6345153 | 0.6780733 | 0.6495132 | 0.6495132 | 0.6816091 |
| X83.10.7.1.1 | 0.7050345 | 0.7050345 | 0.6883991 | 0.6846427 | 0.6724498 | 0.6883991 | 0.7050345 | 0.7209153 | 0.7050345 | 0.7252894 | 0.6588586 | 0.7050345 | 0.7050345 | 0.702351 |
| X84.10.7.1.2 | 0.579749 | 0.579749 | 0.5648817 | 0.5523568 | 0.5506044 | 0.5648817 | 0.579749 | 0.5843982 | 0.579749 | 0.6067802 | 0.5315547 | 0.579749 | 0.579749 | 0.5680135 |
| X85.10.7.1.3 | 0.4833918 | 0.4833918 | 0.4701503 | 0.4529942 | 0.4574191 | 0.4701503 | 0.4833918 | 0.4359119 | 0.4833918 | 0.4701503 | 0.4359342 | 0.4833918 | 0.4833918 | 0.4220636 |
| X86.10.7.1.4 | 0.8391077 | 0.8391077 | 0.8572382 | 0.8209573 | 0.8030601 | 0.8572382 | 0.8391077 | 0.7845438 | 0.8391077 | 0.8206906 | 0.7900395 | 0.8391077 | 0.8391077 | 0.8027739 |
| X88.10.7.1.6 | 0.8105371 | 0.8105371 | 0.7931913 | 0.8286981 | 0.7765957 | 0.7931913 | 0.8105371 | 0.7886235 | 0.8105371 | 0.7931913 | 0.7974888 | 0.8105371 | 0.8105371 | 0.8087067 |
| X89.10.7.1.7 | 0.6657854 | 0.6657854 | 0.6486603 | 0.643286 | 0.6322139 | 0.6486603 | 0.6657854 | 0.5975497 | 0.6657854 | 0.6486603 | 0.6190594 | 0.6657854 | 0.6657854 | 0.6205517 |
| X90.10.7.1.8 | 0.8072824 | 0.8072824 | 0.7895504 | 0.8258372 | 0.7725756 | 0.7895504 | 0.8072824 | 0.8300216 | 0.8072824 | 0.7895504 | 0.8281982 | 0.8072824 | 0.8072824 | 0.8100687 |
| X91.10.7.2.9 | 0.4800774 | 0.4800774 | 0.4660934 | 0.4946331 | 0.4526343 | 0.4660934 | 0.4800774 | 0.4387749 | 0.4800774 | 0.4660934 | 0.4760049 | 0.4800774 | 0.4800774 | 0.4669458 |
| X92.10.7.2.10 | 0.6266379 | 0.6266379 | 0.610078 | 0.6054614 | 0.631302 | 0.610078 | 0.6266379 | 0.5624144 | 0.6266379 | 0.6476213 | 0.5456651 | 0.6266379 | 0.6266379 | 0.5840639 |

Table S3. Correlation coefficient between all seeds. Black boxes indicate individual sites, identified by the SITE column.

| SITE | IND | X11.2.2.1.1 | X12.2.2.1.2 | X13.2.2.1.3 | X14.2.2.2.4 | X15.2.2.2.5 | X17.2.2.2.7 | X18.2.2.2.8 | X19.2.2.2.9 | X21.2.5.2.1 | X23.2.5.2.3 | X25.2.5.3.5 | X28.2.5.1.8 | X30.2.5.1.10 | X31.2.6.1.1 | X32.2.6.1.2 | X33.2.6.1.3 | X35.2.6.1.5 | X36.2.6.2.6 | X38.2.6.2.8 |
|---|---|---|---|---|---|---|---|---|---|---|---|---|---|---|---|---|---|---|---|---|
| S2 | X11.2.2.1.1 | 1.00 | 0.62 | 0.61 | 0.68 | 0.68 | 0.39 | 0.73 | 0.71 | 0.78 | 0.63 | 0.74 | 0.47 | 0.46 | 0.67 | 0.66 | 0.73 | 0.73 | 0.76 | 0.74 |
| S2 | X12.2.2.1.2 | 0.62 | 1.00 | 0.56 | 0.53 | 0.67 | 0.26 | 0.73 | 0.62 | 0.66 | 0.62 | 0.62 | 0.42 | 0.36 | 0.52 | 0.59 | 0.66 | 0.74 | 0.68 | 0.62 |
| S2 | X13.2.2.1.3 | 0.61 | 0.56 | 1.00 | 0.67 | 0.63 | 0.48 | 0.61 | 0.70 | 0.68 | 0.55 | 0.55 | 0.66 | 0.58 | 0.62 | 0.55 | 0.69 | 0.63 | 0.65 | 0.61 |
| S2 | X14.2.2.2.4 | 0.68 | 0.53 | 0.67 | 1.00 | 0.65 | 0.45 | 0.60 | 0.70 | 0.68 | 0.57 | 0.64 | 0.63 | 0.52 | 0.67 | 0.64 | 0.67 | 0.61 | 0.66 | 0.68 |
| S2 | X15.2.2.2.5 | 0.68 | 0.67 | 0.63 | 0.65 | 1.00 | 0.48 | 0.73 | 0.79 | 0.73 | 0.70 | 0.70 | 0.61 | 0.53 | 0.66 | 0.73 | 0.80 | 0.76 | 0.79 | 0.71 |
| S2 | X17.2.2.2.7 | 0.39 | 0.26 | 0.48 | 0.45 | 0.48 | 1.00 | 0.39 | 0.38 | 0.37 | 0.47 | 0.37 | 0.72 | 0.52 | 0.41 | 0.53 | 0.46 | 0.40 | 0.51 | 0.48 |
| S2 | X18.2.2.2.8 | 0.73 | 0.73 | 0.61 | 0.60 | 0.73 | 0.39 | 1.00 | 0.76 | 0.78 | 0.60 | 0.77 | 0.54 | 0.50 | 0.64 | 0.60 | 0.70 | 0.74 | 0.73 | 0.69 |
| S2 | X19.2.2.2.9 | 0.71 | 0.62 | 0.70 | 0.70 | 0.79 | 0.38 | 0.76 | 1.00 | 0.76 | 0.57 | 0.70 | 0.56 | 0.54 | 0.61 | 0.56 | 0.78 | 0.69 | 0.69 | 0.69 |
| S2 | X21.2.5.2.1 | 0.78 | 0.66 | 0.68 | 0.68 | 0.73 | 0.37 | 0.78 | 0.76 | 1.00 | 0.60 | 0.72 | 0.54 | 0.50 | 0.62 | 0.65 | 0.77 | 0.72 | 0.76 | 0.69 |
| S2 | X23.2.5.2.3 | 0.63 | 0.62 | 0.55 | 0.57 | 0.70 | 0.47 | 0.60 | 0.57 | 0.60 | 1.00 | 0.64 | 0.60 | 0.46 | 0.54 | 0.70 | 0.64 | 0.66 | 0.65 | 0.65 |
| S2 | X25.2.5.3.5 | 0.74 | 0.62 | 0.55 | 0.64 | 0.70 | 0.37 | 0.77 | 0.70 | 0.72 | 0.64 | 1.00 | 0.51 | 0.49 | 0.63 | 0.64 | 0.73 | 0.65 | 0.67 | 0.72 |
| S2 | X28.2.5.1.8 | 0.47 | 0.42 | 0.66 | 0.63 | 0.61 | 0.72 | 0.54 | 0.56 | 0.54 | 0.60 | 0.51 | 1.00 | 0.60 | 0.53 | 0.64 | 0.61 | 0.52 | 0.66 | 0.61 |
| S2 | X30.2.5.1.10 | 0.46 | 0.36 | 0.58 | 0.52 | 0.53 | 0.52 | 0.50 | 0.54 | 0.50 | 0.46 | 0.49 | 0.60 | 1.00 | 0.39 | 0.47 | 0.56 | 0.36 | 0.54 | 0.56 |
| S2 | X31.2.6.1.1 | 0.67 | 0.52 | 0.62 | 0.67 | 0.66 | 0.41 | 0.64 | 0.61 | 0.62 | 0.54 | 0.63 | 0.53 | 0.39 | 1.00 | 0.54 | 0.62 | 0.64 | 0.65 | 0.57 |
| S2 | X32.2.6.1.2 | 0.66 | 0.59 | 0.55 | 0.64 | 0.73 | 0.53 | 0.60 | 0.56 | 0.65 | 0.70 | 0.64 | 0.64 | 0.47 | 0.54 | 1.00 | 0.71 | 0.61 | 0.74 | 0.70 |
| S2 | X33.2.6.1.3 | 0.73 | 0.66 | 0.69 | 0.67 | 0.80 | 0.46 | 0.70 | 0.78 | 0.77 | 0.64 | 0.73 | 0.61 | 0.56 | 0.62 | 0.71 | 1.00 | 0.74 | 0.81 | 0.75 |
| S2 | X35.2.6.1.5 | 0.73 | 0.74 | 0.63 | 0.61 | 0.76 | 0.40 | 0.74 | 0.69 | 0.72 | 0.66 | 0.65 | 0.52 | 0.36 | 0.64 | 0.61 | 0.74 | 1.00 | 0.77 | 0.65 |
| S2 | X36.2.6.2.6 | 0.76 | 0.68 | 0.65 | 0.66 | 0.79 | 0.51 | 0.73 | 0.69 | 0.76 | 0.65 | 0.67 | 0.66 | 0.54 | 0.65 | 0.74 | 0.81 | 0.77 | 1.00 | 0.76 |
| S2 | X38.2.6.2.8 | 0.74 | 0.62 | 0.61 | 0.68 | 0.71 | 0.48 | 0.69 | 0.69 | 0.69 | 0.65 | 0.72 | 0.61 | 0.56 | 0.57 | 0.70 | 0.75 | 0.65 | 0.76 | 1.00 |
| S6 | X1.6.9.1.1 | 0.35 | 0.41 | 0.36 | 0.43 | 0.41 | 0.14 | 0.40 | 0.33 | 0.34 | 0.44 | 0.39 | 0.33 | 0.23 | 0.47 | 0.37 | 0.38 | 0.38 | 0.33 | 0.36 |
| S6 | X2.6.9.1.2 | 0.38 | 0.39 | 0.40 | 0.39 | 0.41 | 0.17 | 0.44 | 0.38 | 0.38 | 0.47 | 0.40 | 0.44 | 0.28 | 0.40 | 0.41 | 0.44 | 0.36 | 0.37 | 0.42 |
| S6 | X4.6.9.1.4 | 0.30 | 0.34 | 0.33 | 0.31 | 0.28 | 0.15 | 0.35 | 0.25 | 0.32 | 0.32 | 0.23 | 0.31 | 0.20 | 0.29 | 0.32 | 0.33 | 0.29 | 0.34 | 0.28 |
| S6 | X6.6.9.2.6 | 0.37 | 0.35 | 0.46 | 0.45 | 0.46 | 0.41 | 0.46 | 0.41 | 0.41 | 0.40 | 0.41 | 0.61 | 0.47 | 0.42 | 0.43 | 0.48 | 0.35 | 0.40 | 0.46 |
| S6 | X7.6.9.2.7 | 0.40 | 0.35 | 0.38 | 0.42 | 0.41 | 0.24 | 0.48 | 0.39 | 0.35 | 0.42 | 0.40 | 0.41 | 0.27 | 0.43 | 0.33 | 0.41 | 0.42 | 0.34 | 0.35 |
| S6 | X8.6.9.3.8 | 0.19 | 0.13 | 0.33 | 0.41 | 0.27 | 0.34 | 0.30 | 0.27 | 0.20 | 0.27 | 0.26 | 0.49 | 0.37 | 0.25 | 0.29 | 0.29 | 0.18 | 0.23 | 0.35 |
| S6 | X9.6.9.3.9 | 0.40 | 0.35 | 0.43 | 0.44 | 0.45 | 0.27 | 0.45 | 0.38 | 0.38 | 0.48 | 0.41 | 0.47 | 0.29 | 0.48 | 0.42 | 0.47 | 0.40 | 0.40 | 0.41 |
| S6 | X10.6.9.3.10 | 0.35 | 0.26 | 0.42 | 0.38 | 0.41 | 0.38 | 0.44 | 0.39 | 0.32 | 0.36 | 0.37 | 0.52 | 0.36 | 0.47 | 0.34 | 0.35 | 0.39 | 0.38 | 0.44 |
| S10 | X39.10.1.1.1 | 0.39 | 0.35 | 0.45 | 0.42 | 0.34 | 0.14 | 0.35 | 0.41 | 0.38 | 0.33 | 0.35 | 0.21 | 0.10 | 0.47 | 0.35 | 0.33 | 0.42 | 0.26 | 0.40 |
| S10 | X40.10.1.1.2 | 0.44 | 0.36 | 0.48 | 0.47 | 0.39 | 0.34 | 0.45 | 0.43 | 0.39 | 0.37 | 0.42 | 0.43 | 0.23 | 0.52 | 0.38 | 0.36 | 0.53 | 0.36 | 0.40 |
| S10 | X42.10.1.1.4 | 0.38 | 0.30 | 0.49 | 0.35 | 0.30 | 0.18 | 0.37 | 0.35 | 0.34 | 0.29 | 0.34 | 0.22 | 0.18 | 0.43 | 0.28 | 0.31 | 0.42 | 0.28 | 0.30 |
| S10 | X45.10.1.2.8 | 0.41 | 0.36 | 0.44 | 0.33 | 0.40 | 0.29 | 0.37 | 0.35 | 0.40 | 0.38 | 0.35 | 0.30 | 0.13 | 0.43 | 0.36 | 0.37 | 0.51 | 0.31 | 0.33 |
| S10 | X46.10.1.2.7 | 0.40 | 0.32 | 0.48 | 0.39 | 0.32 | 0.17 | 0.39 | 0.39 | 0.39 | 0.33 | 0.33 | 0.28 | 0.17 | 0.48 | 0.32 | 0.33 | 0.47 | 0.32 | 0.34 |
| S10 | X47.10.1.2.9 | 0.34 | 0.31 | 0.42 | 0.33 | 0.31 | 0.16 | 0.33 | 0.36 | 0.33 | 0.33 | 0.33 | 0.20 | 0.15 | 0.38 | 0.32 | 0.30 | 0.40 | 0.26 | 0.28 |
| S10 | X48.10.1.2.10 | 0.41 | 0.47 | 0.51 | 0.49 | 0.45 | 0.32 | 0.46 | 0.44 | 0.46 | 0.46 | 0.42 | 0.40 | 0.23 | 0.52 | 0.42 | 0.42 | 0.56 | 0.37 | 0.39 |
| S10 | X50.10.2.1.2 | 0.42 | 0.39 | 0.48 | 0.39 | 0.37 | 0.21 | 0.41 | 0.42 | 0.41 | 0.36 | 0.35 | 0.29 | 0.21 | 0.44 | 0.35 | 0.35 | 0.49 | 0.32 | 0.34 |
| S10 | X51.10.2.1.3 | 0.19 | 0.16 | 0.38 | 0.22 | 0.19 | 0.36 | 0.18 | 0.17 | 0.12 | 0.13 | 0.16 | 0.30 | 0.11 | 0.30 | 0.18 | 0.19 | 0.25 | 0.15 | 0.20 |
| S10 | X52.10.2.1.4 | 0.24 | 0.16 | 0.31 | 0.24 | 0.17 | 0.28 | 0.22 | 0.17 | 0.22 | 0.17 | 0.20 | 0.28 | 0.14 | 0.31 | 0.23 | 0.24 | 0.36 | 0.22 | 0.20 |
| S10 | X53.10.2.2.5 | 0.26 | 0.15 | 0.33 | 0.23 | 0.19 | 0.29 | 0.19 | 0.25 | 0.19 | 0.14 | 0.25 | 0.26 | 0.15 | 0.23 | 0.19 | 0.26 | 0.25 | 0.12 | 0.24 |
| S10 | X54.10.2.3.6 | 0.30 | 0.27 | 0.49 | 0.36 | 0.31 | 0.41 | 0.35 | 0.32 | 0.29 | 0.27 | 0.26 | 0.40 | 0.28 | 0.38 | 0.28 | 0.29 | 0.43 | 0.25 | 0.24 |
| S10 | X55.10.2.3.7 | 0.38 | 0.34 | 0.39 | 0.38 | 0.39 | 0.31 | 0.36 | 0.37 | 0.38 | 0.31 | 0.31 | 0.34 | 0.26 | 0.42 | 0.34 | 0.32 | 0.50 | 0.30 | 0.34 |
| S10 | X56.10.2.3.8 | 0.38 | 0.36 | 0.46 | 0.35 | 0.36 | 0.15 | 0.37 | 0.38 | 0.40 | 0.35 | 0.31 | 0.29 | 0.15 | 0.40 | 0.36 | 0.34 | 0.48 | 0.33 | 0.30 |
| S10 | X57.10.2.3.9 | 0.37 | 0.28 | 0.44 | 0.38 | 0.26 | 0.29 | 0.34 | 0.31 | 0.37 | 0.29 | 0.28 | 0.26 | 0.19 | 0.39 | 0.30 | 0.31 | 0.41 | 0.27 | 0.32 |
| S10 | X58.10.2.3.10 | 0.21 | 0.19 | 0.38 | 0.20 | 0.25 | 0.31 | 0.20 | 0.23 | 0.20 | 0.23 | 0.18 | 0.30 | 0.18 | 0.29 | 0.19 | 0.20 | 0.34 | 0.17 | 0.18 |
| S10 | X59.10.3.1.1 | 0.32 | 0.21 | 0.50 | 0.34 | 0.27 | 0.40 | 0.34 | 0.31 | 0.28 | 0.23 | 0.25 | 0.36 | 0.28 | 0.41 | 0.24 | 0.22 | 0.36 | 0.24 | 0.23 |
| S10 | X60.10.3.1.2 | 0.37 | 0.27 | 0.38 | 0.38 | 0.28 | 0.17 | 0.43 | 0.37 | 0.33 | 0.25 | 0.33 | 0.29 | 0.14 | 0.42 | 0.27 | 0.28 | 0.44 | 0.30 | 0.32 |
| S10 | X61.10.3.2.3 | 0.32 | 0.28 | 0.39 | 0.31 | 0.28 | 0.12 | 0.28 | 0.26 | 0.28 | 0.26 | 0.26 | 0.22 | 0.08 | 0.42 | 0.30 | 0.30 | 0.39 | 0.23 | 0.24 |
| S10 | X63.10.3.2.5 | 0.30 | 0.28 | 0.40 | 0.32 | 0.28 | 0.11 | 0.29 | 0.29 | 0.29 | 0.29 | 0.26 | 0.21 | 0.10 | 0.34 | 0.25 | 0.26 | 0.39 | 0.23 | 0.22 |
| S10 | X66.10.3.3.8 | 0.27 | 0.19 | 0.36 | 0.23 | 0.18 | 0.09 | 0.23 | 0.26 | 0.26 | 0.21 | 0.20 | 0.16 | 0.11 | 0.29 | 0.22 | 0.20 | 0.34 | 0.17 | 0.21 |
| S10 | X67.10.5.1.1 | 0.42 | 0.22 | 0.31 | 0.25 | 0.29 | 0.24 | 0.35 | 0.22 | 0.33 | 0.20 | 0.25 | 0.23 | 0.15 | 0.33 | 0.26 | 0.27 | 0.38 | 0.29 | 0.23 |
| S10 | X69.10.5.1.3 | 0.31 | 0.22 | 0.37 | 0.36 | 0.31 | 0.35 | 0.29 | 0.26 | 0.26 | 0.27 | 0.26 | 0.37 | 0.18 | 0.42 | 0.28 | 0.26 | 0.41 | 0.23 | 0.24 |
| S10 | X72.10.5.4.6 | 0.28 | 0.21 | 0.37 | 0.26 | 0.26 | 0.36 | 0.25 | 0.24 | 0.25 | 0.20 | 0.22 | 0.33 | 0.22 | 0.30 | 0.25 | 0.26 | 0.35 | 0.18 | 0.20 |
| S10 | X73.10.5.4.7 | 0.25 | 0.29 | 0.37 | 0.28 | 0.26 | 0.12 | 0.26 | 0.24 | 0.26 | 0.24 | 0.24 | 0.17 | 0.09 | 0.29 | 0.27 | 0.28 | 0.37 | 0.21 | 0.19 |
| S10 | X74.10.5.4.8 | 0.24 | 0.20 | 0.35 | 0.25 | 0.28 | 0.27 | 0.20 | 0.23 | 0.20 | 0.25 | 0.21 | 0.31 | 0.16 | 0.39 | 0.24 | 0.24 | 0.33 | 0.17 | 0.22 |
| S10 | X75.10.6.1.1 | 0.42 | 0.31 | 0.47 | 0.38 | 0.34 | 0.22 | 0.34 | 0.41 | 0.34 | 0.29 | 0.37 | 0.23 | 0.15 | 0.41 | 0.28 | 0.34 | 0.46 | 0.28 | 0.33 |
| S10 | X76.10.6.1.2 | 0.31 | 0.26 | 0.47 | 0.41 | 0.28 | 0.42 | 0.35 | 0.29 | 0.32 | 0.24 | 0.29 | 0.41 | 0.19 | 0.35 | 0.28 | 0.29 | 0.41 | 0.26 | 0.30 |
| S10 | X78.10.6.1.4 | 0.34 | 0.28 | 0.51 | 0.33 | 0.38 | 0.45 | 0.29 | 0.33 | 0.29 | 0.36 | 0.32 | 0.33 | 0.26 | 0.42 | 0.34 | 0.38 | 0.43 | 0.26 | 0.31 |
| S10 | X79.10.6.1.5 | 0.25 | 0.29 | 0.40 | 0.22 | 0.26 | 0.10 | 0.27 | 0.30 | 0.27 | 0.22 | 0.25 | 0.18 | 0.10 | 0.33 | 0.21 | 0.24 | 0.35 | 0.21 | 0.23 |
| S10 | X80.10.6.2.6 | 0.35 | 0.26 | 0.38 | 0.29 | 0.27 | 0.08 | 0.34 | 0.31 | 0.31 | 0.26 | 0.28 | 0.18 | 0.14 | 0.37 | 0.28 | 0.26 | 0.41 | 0.28 | 0.24 |
| S10 | X81.10.6.2.7 | 0.37 | 0.27 | 0.44 | 0.41 | 0.35 | 0.27 | 0.38 | 0.39 | 0.33 | 0.28 | 0.38 | 0.36 | 0.18 | 0.47 | 0.31 | 0.30 | 0.44 | 0.29 | 0.34 |
| S10 | X82.10.6.2.8 | 0.38 | 0.31 | 0.50 | 0.39 | 0.31 | 0.14 | 0.34 | 0.35 | 0.37 | 0.26 | 0.31 | 0.27 | 0.10 | 0.40 | 0.30 | 0.31 | 0.42 | 0.25 | 0.30 |
| S10 | X83.10.7.1.1 | 0.41 | 0.36 | 0.48 | 0.39 | 0.36 | 0.31 | 0.47 | 0.37 | 0.45 | 0.34 | 0.36 | 0.40 | 0.23 | 0.46 | 0.38 | 0.38 | 0.53 | 0.36 | 0.35 |
| S10 | X84.10.7.1.2 | 0.20 | 0.21 | 0.25 | 0.20 | 0.14 | ### | 0.19 | 0.22 | 0.26 | 0.14 | 0.14 | 0.10 | ### | 0.18 | 0.16 | 0.14 | 0.30 | 0.13 | 0.15 |
| S10 | X85.10.7.1.3 | 0.21 | 0.15 | 0.25 | 0.24 | 0.21 | 0.38 | 0.26 | 0.16 | 0.17 | 0.15 | 0.14 | 0.32 | 0.14 | 0.28 | 0.14 | 0.12 | 0.27 | 0.17 | 0.21 |
| S10 | X86.10.7.1.4 | 0.32 | 0.30 | 0.43 | 0.28 | 0.30 | 0.21 | 0.30 | 0.34 | 0.30 | 0.31 | 0.28 | 0.22 | 0.08 | 0.40 | 0.29 | 0.27 | 0.41 | 0.24 | 0.23 |
| S10 | X88.10.7.1.6 | 0.31 | 0.32 | 0.40 | 0.31 | 0.26 | 0.14 | 0.30 | 0.30 | 0.36 | 0.25 | 0.25 | 0.21 | ### | 0.36 | 0.26 | 0.27 | 0.41 | 0.21 | 0.23 |
| S10 | X89.10.7.1.7 | 0.34 | 0.29 | 0.50 | 0.35 | 0.28 | 0.39 | 0.35 | 0.32 | 0.32 | 0.27 | 0.27 | 0.31 | 0.30 | 0.35 | 0.28 | 0.26 | 0.38 | 0.26 | 0.27 |
| S10 | X90.10.7.1.8 | 0.23 | 0.24 | 0.33 | 0.21 | 0.18 | 0.03 | 0.21 | 0.24 | 0.26 | 0.14 | 0.17 | 0.10 | 0.05 | 0.24 | 0.18 | 0.16 | 0.29 | 0.13 | 0.17 |
| S10 | X91.10.7.2.9 | 0.25 | 0.20 | 0.37 | 0.33 | 0.26 | 0.42 | 0.27 | 0.27 | 0.21 | 0.19 | 0.22 | 0.41 | 0.27 | 0.33 | 0.23 | 0.18 | 0.35 | 0.21 | 0.22 |
| S10 | X92.10.7.2.10 | 0.28 | 0.26 | 0.43 | 0.37 | 0.28 | 0.41 | 0.31 | 0.35 | 0.26 | 0.26 | 0.32 | 0.38 | 0.23 | 0.38 | 0.24 | 0.26 | 0.37 | 0.20 | 0.29 |

| SITE | IND | X1.6.9.1.1 | X2.6.9.1.2 | X4.6.9.1.4 | X6.6.9.2.6 | X7.6.9.2.7 | X8.6.9.3.8 | X9.6.9.3.9 | X10.6.9.3.10 | X39.10.1.1.1 | X40.10.1.1.2 | X42.10.1.1.4 | X45.10.1.2.8 | X46.10.1.2.7 | X47.10.1.2.9 | X48.10.1.2.10 | X50.10.2.1.2 |
|---|---|---|---|---|---|---|---|---|---|---|---|---|---|---|---|---|---|
| S2 | X11.2.2.1.1 | 0.35 | 0.38 | 0.30 | 0.37 | 0.40 | 0.19 | 0.40 | 0.35 | 0.39 | 0.44 | 0.38 | 0.41 | 0.40 | 0.34 | 0.41 | 0.42 |
| S2 | X12.2.2.1.2 | 0.41 | 0.39 | 0.34 | 0.35 | 0.35 | 0.13 | 0.35 | 0.26 | 0.35 | 0.36 | 0.30 | 0.36 | 0.32 | 0.31 | 0.47 | 0.39 |
| S2 | X13.2.2.1.3 | 0.36 | 0.40 | 0.33 | 0.46 | 0.38 | 0.33 | 0.43 | 0.42 | 0.45 | 0.48 | 0.49 | 0.44 | 0.48 | 0.42 | 0.51 | 0.48 |
| S2 | X14.2.2.2.4 | 0.43 | 0.39 | 0.31 | 0.45 | 0.42 | 0.41 | 0.44 | 0.38 | 0.42 | 0.47 | 0.35 | 0.33 | 0.39 | 0.33 | 0.49 | 0.39 |
| S2 | X15.2.2.2.5 | 0.41 | 0.41 | 0.28 | 0.46 | 0.41 | 0.27 | 0.45 | 0.41 | 0.34 | 0.39 | 0.30 | 0.40 | 0.32 | 0.31 | 0.45 | 0.37 |
| S2 | X17.2.2.2.7 | 0.14 | 0.17 | 0.15 | 0.41 | 0.24 | 0.34 | 0.27 | 0.38 | 0.14 | 0.34 | 0.18 | 0.29 | 0.17 | 0.16 | 0.32 | 0.21 |
| S2 | X18.2.2.2.8 | 0.40 | 0.44 | 0.35 | 0.46 | 0.48 | 0.30 | 0.45 | 0.44 | 0.35 | 0.45 | 0.37 | 0.37 | 0.39 | 0.33 | 0.46 | 0.41 |
| S2 | X19.2.2.2.9 | 0.33 | 0.38 | 0.25 | 0.41 | 0.39 | 0.27 | 0.38 | 0.39 | 0.41 | 0.43 | 0.35 | 0.35 | 0.39 | 0.36 | 0.44 | 0.42 |
| S2 | X21.2.5.2.1 | 0.34 | 0.38 | 0.32 | 0.41 | 0.35 | 0.20 | 0.38 | 0.32 | 0.38 | 0.39 | 0.34 | 0.40 | 0.39 | 0.33 | 0.46 | 0.41 |
| S2 | X23.2.5.2.3 | 0.44 | 0.47 | 0.32 | 0.40 | 0.42 | 0.27 | 0.48 | 0.36 | 0.33 | 0.37 | 0.29 | 0.38 | 0.33 | 0.33 | 0.46 | 0.36 |
| S2 | X25.2.5.3.5 | 0.39 | 0.40 | 0.23 | 0.41 | 0.40 | 0.26 | 0.41 | 0.37 | 0.35 | 0.42 | 0.34 | 0.35 | 0.33 | 0.33 | 0.42 | 0.35 |
| S2 | X28.2.5.1.8 | 0.33 | 0.44 | 0.31 | 0.61 | 0.41 | 0.49 | 0.47 | 0.52 | 0.21 | 0.43 | 0.22 | 0.30 | 0.28 | 0.20 | 0.40 | 0.29 |
| S2 | X30.2.5.1.10 | 0.23 | 0.28 | 0.20 | 0.47 | 0.27 | 0.37 | 0.29 | 0.36 | 0.10 | 0.23 | 0.18 | 0.13 | 0.17 | 0.15 | 0.23 | 0.21 |
| S2 | X31.2.6.1.1 | 0.47 | 0.40 | 0.29 | 0.42 | 0.43 | 0.25 | 0.48 | 0.47 | 0.47 | 0.52 | 0.43 | 0.43 | 0.48 | 0.38 | 0.52 | 0.44 |
| S2 | X32.2.6.1.2 | 0.37 | 0.41 | 0.32 | 0.43 | 0.33 | 0.29 | 0.42 | 0.34 | 0.35 | 0.38 | 0.28 | 0.36 | 0.32 | 0.32 | 0.42 | 0.35 |
| S2 | X33.2.6.1.3 | 0.38 | 0.44 | 0.33 | 0.48 | 0.41 | 0.29 | 0.47 | 0.35 | 0.33 | 0.36 | 0.31 | 0.37 | 0.33 | 0.30 | 0.42 | 0.35 |
| S2 | X35.2.6.1.5 | 0.38 | 0.36 | 0.29 | 0.35 | 0.42 | 0.18 | 0.40 | 0.39 | 0.42 | 0.53 | 0.42 | 0.51 | 0.47 | 0.40 | 0.56 | 0.49 |
| S2 | X36.2.6.2.6 | 0.33 | 0.37 | 0.34 | 0.40 | 0.34 | 0.23 | 0.40 | 0.38 | 0.26 | 0.36 | 0.28 | 0.31 | 0.32 | 0.26 | 0.37 | 0.32 |
| S2 | X38.2.6.2.8 | 0.36 | 0.42 | 0.28 | 0.46 | 0.35 | 0.35 | 0.41 | 0.44 | 0.40 | 0.40 | 0.30 | 0.33 | 0.34 | 0.28 | 0.39 | 0.34 |
| S6 | X1.6.9.1.1 | 1.00 | 0.81 | 0.63 | 0.62 | 0.58 | 0.51 | 0.78 | 0.48 | 0.34 | 0.29 | 0.28 | 0.24 | 0.26 | 0.24 | 0.44 | 0.30 |
| S6 | X2.6.9.1.2 | 0.81 | 1.00 | 0.71 | 0.71 | 0.67 | 0.61 | 0.85 | 0.55 | 0.30 | 0.26 | 0.25 | 0.21 | 0.23 | 0.21 | 0.39 | 0.24 |
| S6 | X4.6.9.1.4 | 0.63 | 0.71 | 1.00 | 0.60 | 0.56 | 0.45 | 0.68 | 0.40 | 0.34 | 0.32 | 0.42 | 0.29 | 0.30 | 0.35 | 0.37 | 0.33 |
| S6 | X6.6.9.2.6 | 0.62 | 0.71 | 0.60 | 1.00 | 0.59 | 0.60 | 0.67 | 0.54 | 0.19 | 0.35 | 0.28 | 0.21 | 0.19 | 0.21 | 0.27 | 0.24 |
| S6 | X7.6.9.2.7 | 0.58 | 0.67 | 0.56 | 0.59 | 1.00 | 0.53 | 0.53 | 0.57 | 0.33 | 0.40 | 0.41 | 0.33 | 0.34 | 0.37 | 0.36 | 0.36 |
| S6 | X8.6.9.3.8 | 0.51 | 0.61 | 0.45 | 0.60 | 0.53 | 1.00 | 0.54 | 0.55 | 0.21 | 0.31 | 0.18 | 0.23 | 0.20 | 0.14 | 0.31 | 0.20 |
| S6 | X9.6.9.3.9 | 0.78 | 0.85 | 0.68 | 0.67 | 0.53 | 0.54 | 1.00 | 0.46 | 0.32 | 0.31 | 0.30 | 0.26 | 0.29 | 0.26 | 0.42 | 0.29 |
| S6 | X10.6.9.3.10 | 0.48 | 0.55 | 0.40 | 0.54 | 0.57 | 0.55 | 0.46 | 1.00 | 0.38 | 0.45 | 0.33 | 0.36 | 0.37 | 0.32 | 0.44 | 0.32 |
| S10 | X39.10.1.1.1 | 0.34 | 0.30 | 0.34 | 0.19 | 0.33 | 0.21 | 0.32 | 0.38 | 1.00 | 0.59 | 0.72 | 0.54 | 0.71 | 0.73 | 0.64 | 0.65 |
| S10 | X40.10.1.1.2 | 0.29 | 0.26 | 0.32 | 0.35 | 0.40 | 0.31 | 0.31 | 0.45 | 0.59 | 1.00 | 0.65 | 0.67 | 0.67 | 0.70 | 0.70 | 0.66 |
| S10 | X42.10.1.1.4 | 0.28 | 0.25 | 0.42 | 0.28 | 0.41 | 0.18 | 0.30 | 0.33 | 0.72 | 0.65 | 1.00 | 0.52 | 0.78 | 0.77 | 0.55 | 0.73 |
| S10 | X45.10.1.2.8 | 0.24 | 0.21 | 0.29 | 0.21 | 0.33 | 0.23 | 0.26 | 0.36 | 0.54 | 0.67 | 0.52 | 1.00 | 0.60 | 0.59 | 0.70 | 0.49 |
| S10 | X46.10.1.2.7 | 0.26 | 0.23 | 0.30 | 0.19 | 0.34 | 0.20 | 0.29 | 0.37 | 0.71 | 0.67 | 0.78 | 0.60 | 1.00 | 0.72 | 0.57 | 0.72 |
| S10 | X47.10.1.2.9 | 0.24 | 0.21 | 0.35 | 0.21 | 0.37 | 0.14 | 0.26 | 0.32 | 0.73 | 0.70 | 0.77 | 0.59 | 0.72 | 1.00 | 0.63 | 0.75 |
| S10 | X48.10.1.2.10 | 0.44 | 0.39 | 0.37 | 0.27 | 0.36 | 0.31 | 0.42 | 0.44 | 0.64 | 0.70 | 0.55 | 0.70 | 0.57 | 0.63 | 1.00 | 0.55 |
| S10 | X50.10.2.1.2 | 0.30 | 0.24 | 0.33 | 0.24 | 0.36 | 0.20 | 0.29 | 0.32 | 0.65 | 0.66 | 0.73 | 0.49 | 0.72 | 0.75 | 0.55 | 1.00 |
| S10 | X51.10.2.1.3 | 0.02 | 0.02 | 0.12 | 0.19 | 0.18 | 0.21 | 0.06 | 0.24 | 0.30 | 0.48 | 0.47 | 0.44 | 0.31 | 0.22 | 0.33 | 0.20 |
| S10 | X52.10.2.1.4 | 0.11 | 0.14 | 0.24 | 0.23 | 0.22 | 0.23 | 0.13 | 0.33 | 0.41 | 0.56 | 0.55 | 0.51 | 0.41 | 0.41 | 0.51 | 0.38 |
| S10 | X53.10.2.2.5 | 0.19 | 0.14 | 0.22 | 0.27 | 0.23 | 0.22 | 0.18 | 0.35 | 0.33 | 0.49 | 0.36 | 0.41 | 0.31 | 0.33 | 0.44 | 0.31 |
| S10 | X54.10.2.3.6 | 0.23 | 0.20 | 0.30 | 0.29 | 0.30 | 0.31 | 0.26 | 0.37 | 0.40 | 0.68 | 0.53 | 0.61 | 0.56 | 0.51 | 0.64 | 0.46 |
| S10 | X55.10.2.3.7 | 0.20 | 0.15 | 0.24 | 0.32 | 0.28 | 0.30 | 0.21 | 0.44 | 0.46 | 0.66 | 0.55 | 0.60 | 0.57 | 0.47 | 0.52 | 0.56 |
| S10 | X56.10.2.3.8 | 0.31 | 0.30 | 0.44 | 0.23 | 0.41 | 0.19 | 0.28 | 0.39 | 0.68 | 0.64 | 0.75 | 0.67 | 0.83 | 0.73 | 0.61 | 0.72 |
| S10 | X57.10.2.3.9 | 0.23 | 0.21 | 0.32 | 0.25 | 0.38 | 0.18 | 0.23 | 0.33 | 0.59 | 0.61 | 0.64 | 0.60 | 0.70 | 0.62 | 0.57 | 0.61 |
| S10 | X58.10.2.3.10 | 0.16 | 0.14 | 0.23 | 0.28 | 0.23 | 0.29 | 0.20 | 0.36 | 0.42 | 0.63 | 0.60 | 0.60 | 0.52 | 0.58 | 0.53 | 0.54 |
| S10 | X59.10.3.1.1 | 0.15 | 0.13 | 0.31 | 0.28 | 0.32 | 0.32 | 0.19 | 0.42 | 0.41 | 0.63 | 0.60 | 0.60 | 0.59 | 0.54 | 0.53 | 0.46 |
| S10 | X60.10.3.1.2 | 0.29 | 0.26 | 0.32 | 0.19 | 0.38 | 0.27 | 0.28 | 0.39 | 0.59 | 0.62 | 0.69 | 0.46 | 0.71 | 0.58 | 0.50 | 0.61 |
| S10 | X61.10.3.2.3 | 0.19 | 0.17 | 0.32 | 0.18 | 0.34 | 0.16 | 0.22 | 0.30 | 0.54 | 0.62 | 0.72 | 0.62 | 0.71 | 0.65 | 0.54 | 0.68 |
| S10 | X63.10.3.2.5 | 0.22 | 0.20 | 0.33 | 0.15 | 0.32 | 0.14 | 0.22 | 0.33 | 0.64 | 0.60 | 0.67 | 0.69 | 0.75 | 0.75 | 0.60 | 0.62 |
| S10 | X66.10.3.3.8 | 0.16 | 0.12 | 0.25 | 0.08 | 0.20 | 0.09 | 0.17 | 0.30 | 0.67 | 0.60 | 0.71 | 0.50 | 0.79 | 0.72 | 0.50 | 0.68 |
| S10 | X67.10.5.1.1 | 0.10 | 0.11 | 0.24 | 0.21 | 0.31 | 0.17 | 0.11 | 0.24 | 0.24 | 0.44 | 0.41 | 0.39 | 0.21 | 0.36 | 0.36 | 0.35 |
| S10 | X69.10.5.1.3 | 0.17 | 0.15 | 0.29 | 0.35 | 0.43 | 0.27 | 0.20 | 0.37 | 0.46 | 0.72 | 0.65 | 0.61 | 0.56 | 0.59 | 0.51 | 0.51 |
| S10 | X72.10.5.4.6 | 0.16 | 0.14 | 0.29 | 0.27 | 0.26 | 0.26 | 0.18 | 0.35 | 0.35 | 0.52 | 0.46 | 0.51 | 0.42 | 0.36 | 0.44 | 0.34 |
| S10 | X73.10.5.4.7 | 0.21 | 0.18 | 0.36 | 0.15 | 0.32 | 0.13 | 0.22 | 0.25 | 0.63 | 0.60 | 0.74 | 0.61 | 0.60 | 0.79 | 0.61 | 0.63 |
| S10 | X74.10.5.4.8 | 0.14 | 0.12 | 0.18 | 0.22 | 0.17 | 0.24 | 0.17 | 0.33 | 0.42 | 0.54 | 0.49 | 0.53 | 0.52 | 0.34 | 0.46 | 0.36 |
| S10 | X75.10.6.1.1 | 0.22 | 0.20 | 0.34 | 0.23 | 0.35 | 0.16 | 0.25 | 0.36 | 0.63 | 0.65 | 0.76 | 0.56 | 0.71 | 0.78 | 0.59 | 0.67 |
| S10 | X76.10.6.1.2 | 0.21 | 0.19 | 0.28 | 0.34 | 0.30 | 0.29 | 0.22 | 0.39 | 0.40 | 0.60 | 0.47 | 0.41 | 0.39 | 0.35 | 0.51 | 0.30 |
| S10 | X78.10.6.1.4 | 0.26 | 0.23 | 0.27 | 0.29 | 0.30 | 0.25 | 0.29 | 0.34 | 0.49 | 0.64 | 0.57 | 0.60 | 0.45 | 0.55 | 0.60 | 0.50 |
| S10 | X79.10.6.1.5 | 0.26 | 0.26 | 0.30 | 0.19 | 0.25 | 0.20 | 0.26 | 0.32 | 0.64 | 0.54 | 0.78 | 0.47 | 0.69 | 0.72 | 0.57 | 0.61 |
| S10 | X80.10.6.2.6 | 0.24 | 0.21 | 0.37 | 0.18 | 0.35 | 0.13 | 0.25 | 0.33 | 0.63 | 0.54 | 0.81 | 0.54 | 0.76 | 0.72 | 0.47 | 0.68 |
| S10 | X81.10.6.2.7 | 0.20 | 0.18 | 0.26 | 0.24 | 0.31 | 0.27 | 0.23 | 0.36 | 0.51 | 0.80 | 0.70 | 0.48 | 0.62 | 0.59 | 0.54 | 0.65 |
| S10 | X82.10.6.2.8 | 0.25 | 0.22 | 0.37 | 0.25 | 0.40 | 0.21 | 0.25 | 0.38 | 0.64 | 0.63 | 0.74 | 0.63 | 0.72 | 0.69 | 0.54 | 0.65 |
| S10 | X83.10.7.1.1 | 0.21 | 0.19 | 0.30 | 0.30 | 0.37 | 0.21 | 0.27 | 0.35 | 0.52 | 0.67 | 0.68 | 0.64 | 0.67 | 0.60 | 0.58 | 0.63 |
| S10 | X84.10.7.1.2 | 0.03 | 0.02 | 0.13 | ### | 0.16 | 0.03 | 0.04 | 0.19 | 0.48 | 0.47 | 0.50 | 0.57 | 0.59 | 0.58 | 0.43 | 0.54 |
| S10 | X85.10.7.1.3 | 0.12 | 0.11 | 0.19 | 0.24 | 0.23 | 0.31 | 0.13 | 0.43 | 0.32 | 0.53 | 0.38 | 0.49 | 0.37 | 0.24 | 0.42 | 0.19 |
| S10 | X86.10.7.1.4 | 0.26 | 0.23 | 0.35 | 0.19 | 0.31 | 0.14 | 0.29 | 0.32 | 0.68 | 0.67 | 0.82 | 0.67 | 0.77 | 0.80 | 0.64 | 0.73 |
| S10 | X88.10.7.1.6 | 0.29 | 0.26 | 0.40 | 0.19 | 0.34 | 0.14 | 0.29 | 0.29 | 0.74 | 0.63 | 0.78 | 0.60 | 0.73 | 0.72 | 0.63 | 0.72 |
| S10 | X89.10.7.1.7 | 0.24 | 0.19 | 0.36 | 0.28 | 0.30 | 0.32 | 0.22 | 0.42 | 0.51 | 0.63 | 0.61 | 0.54 | 0.56 | 0.56 | 0.60 | 0.59 |
| S10 | X90.10.7.1.8 | 0.12 | 0.13 | 0.32 | 0.13 | 0.27 | 0.09 | 0.11 | 0.25 | 0.66 | 0.55 | 0.71 | 0.55 | 0.64 | 0.76 | 0.52 | 0.57 |
| S10 | X91.10.7.2.9 | 0.18 | 0.13 | 0.27 | 0.29 | 0.27 | 0.39 | 0.16 | 0.40 | 0.36 | 0.61 | 0.44 | 0.44 | 0.43 | 0.38 | 0.48 | 0.41 |
| S10 | X92.10.7.2.10 | 0.21 | 0.18 | 0.20 | 0.25 | 0.32 | 0.28 | 0.19 | 0.40 | 0.47 | 0.65 | 0.53 | 0.53 | 0.55 | 0.45 | 0.56 | 0.51 |

| SITE | IND | X51.10.2.1.3 | X52.10.2.1.4 | X53.10.2.2.5 | X54.10.2.3.6 | X55.10.2.3.7 | X56.10.2.3.8 | X57.10.2.3.9 | X58.10.2.3.10 | X59.10.3.1.1 | X60.10.3.1.2 | X61.10.3.2.3 | X63.10.3.2.5 | X66.10.3.3.8 | X67.10.5.1.1 | X69.10.5.1.3 | X72.10.5.4.6 | X73.10.5.4.7 | X74.10.5.4.8 | X75.10.6.1.1 | X76.10.6.1.2 |
|---|---|---|---|---|---|---|---|---|---|---|---|---|---|---|---|---|---|---|---|---|---|
| S2 | X11.2.2.1.1 | 0.19 | 0.24 | 0.26 | 0.30 | 0.38 | 0.38 | 0.37 | 0.21 | 0.32 | 0.37 | 0.32 | 0.30 | 0.27 | 0.42 | 0.31 | 0.28 | 0.25 | 0.24 | 0.42 | 0.31 |
| S2 | X12.2.2.1.2 | 0.16 | 0.16 | 0.15 | 0.27 | 0.34 | 0.36 | 0.28 | 0.19 | 0.21 | 0.27 | 0.28 | 0.28 | 0.19 | 0.22 | 0.22 | 0.21 | 0.29 | 0.20 | 0.31 | 0.26 |
| S2 | X13.2.2.1.3 | 0.38 | 0.31 | 0.33 | 0.49 | 0.39 | 0.46 | 0.44 | 0.38 | 0.50 | 0.38 | 0.39 | 0.40 | 0.36 | 0.31 | 0.37 | 0.37 | 0.37 | 0.35 | 0.47 | 0.47 |
| S2 | X14.2.2.2.4 | 0.22 | 0.24 | 0.23 | 0.36 | 0.38 | 0.35 | 0.38 | 0.20 | 0.34 | 0.38 | 0.31 | 0.32 | 0.23 | 0.25 | 0.36 | 0.26 | 0.28 | 0.25 | 0.38 | 0.41 |
| S2 | X15.2.2.2.5 | 0.19 | 0.17 | 0.19 | 0.31 | 0.39 | 0.36 | 0.26 | 0.25 | 0.27 | 0.28 | 0.28 | 0.28 | 0.18 | 0.29 | 0.31 | 0.26 | 0.26 | 0.28 | 0.34 | 0.28 |
| S2 | X17.2.2.2.7 | 0.36 | 0.28 | 0.29 | 0.41 | 0.31 | 0.15 | 0.29 | 0.31 | 0.40 | 0.17 | 0.12 | 0.11 | 0.09 | 0.24 | 0.35 | 0.36 | 0.12 | 0.27 | 0.22 | 0.42 |
| S2 | X18.2.2.2.8 | 0.18 | 0.22 | 0.19 | 0.35 | 0.36 | 0.37 | 0.34 | 0.20 | 0.34 | 0.43 | 0.28 | 0.29 | 0.23 | 0.35 | 0.29 | 0.25 | 0.26 | 0.20 | 0.34 | 0.35 |
| S2 | X19.2.2.2.9 | 0.17 | 0.17 | 0.25 | 0.32 | 0.37 | 0.38 | 0.31 | 0.23 | 0.31 | 0.37 | 0.26 | 0.29 | 0.26 | 0.22 | 0.26 | 0.24 | 0.24 | 0.23 | 0.41 | 0.29 |
| S2 | X21.2.5.2.1 | 0.12 | 0.22 | 0.19 | 0.29 | 0.38 | 0.40 | 0.37 | 0.20 | 0.28 | 0.33 | 0.28 | 0.29 | 0.26 | 0.33 | 0.26 | 0.25 | 0.26 | 0.20 | 0.34 | 0.32 |
| S2 | X23.2.5.2.3 | 0.13 | 0.17 | 0.14 | 0.27 | 0.31 | 0.35 | 0.29 | 0.23 | 0.23 | 0.25 | 0.26 | 0.29 | 0.21 | 0.20 | 0.27 | 0.20 | 0.24 | 0.25 | 0.29 | 0.24 |
| S2 | X25.2.5.3.5 | 0.16 | 0.20 | 0.25 | 0.26 | 0.31 | 0.31 | 0.28 | 0.18 | 0.25 | 0.33 | 0.26 | 0.26 | 0.20 | 0.25 | 0.26 | 0.22 | 0.24 | 0.21 | 0.37 | 0.29 |
| S2 | X28.2.5.1.8 | 0.30 | 0.28 | 0.26 | 0.40 | 0.34 | 0.29 | 0.26 | 0.30 | 0.36 | 0.29 | 0.22 | 0.21 | 0.16 | 0.23 | 0.37 | 0.33 | 0.17 | 0.31 | 0.23 | 0.41 |
| S2 | X30.2.5.1.10 | 0.11 | 0.14 | 0.15 | 0.28 | 0.26 | 0.15 | 0.19 | 0.18 | 0.28 | 0.14 | 0.08 | 0.10 | 0.11 | 0.15 | 0.18 | 0.22 | 0.09 | 0.16 | 0.15 | 0.19 |
| S2 | X31.2.6.1.1 | 0.30 | 0.31 | 0.23 | 0.38 | 0.42 | 0.40 | 0.39 | 0.29 | 0.41 | 0.42 | 0.42 | 0.34 | 0.29 | 0.33 | 0.42 | 0.30 | 0.29 | 0.39 | 0.41 | 0.35 |
| S2 | X32.2.6.1.2 | 0.18 | 0.23 | 0.19 | 0.28 | 0.34 | 0.36 | 0.30 | 0.19 | 0.24 | 0.27 | 0.30 | 0.25 | 0.22 | 0.26 | 0.28 | 0.25 | 0.27 | 0.24 | 0.28 | 0.28 |
| S2 | X33.2.6.1.3 | 0.19 | 0.24 | 0.26 | 0.29 | 0.32 | 0.34 | 0.31 | 0.20 | 0.22 | 0.28 | 0.30 | 0.26 | 0.20 | 0.27 | 0.26 | 0.26 | 0.28 | 0.24 | 0.34 | 0.29 |
| S2 | X35.2.6.1.5 | 0.25 | 0.36 | 0.25 | 0.43 | 0.50 | 0.48 | 0.41 | 0.34 | 0.36 | 0.44 | 0.39 | 0.39 | 0.34 | 0.38 | 0.41 | 0.35 | 0.37 | 0.33 | 0.46 | 0.41 |
| S2 | X36.2.6.2.6 | 0.15 | 0.22 | 0.12 | 0.25 | 0.30 | 0.33 | 0.27 | 0.17 | 0.24 | 0.30 | 0.23 | 0.23 | 0.17 | 0.29 | 0.23 | 0.18 | 0.21 | 0.17 | 0.28 | 0.26 |
| S2 | X38.2.6.2.8 | 0.20 | 0.20 | 0.24 | 0.24 | 0.34 | 0.30 | 0.32 | 0.18 | 0.23 | 0.32 | 0.24 | 0.22 | 0.21 | 0.23 | 0.24 | 0.20 | 0.19 | 0.22 | 0.33 | 0.30 |
| S6 | X1.6.9.1.1 | 0.02 | 0.11 | 0.19 | 0.23 | 0.20 | 0.31 | 0.23 | 0.16 | 0.15 | 0.29 | 0.19 | 0.22 | 0.16 | 0.10 | 0.17 | 0.16 | 0.21 | 0.14 | 0.22 | 0.21 |
| S6 | X2.6.9.1.2 | 0.02 | 0.14 | 0.14 | 0.20 | 0.15 | 0.30 | 0.21 | 0.14 | 0.13 | 0.26 | 0.17 | 0.20 | 0.12 | 0.11 | 0.15 | 0.14 | 0.18 | 0.12 | 0.20 | 0.19 |
| S6 | X4.6.9.1.4 | 0.12 | 0.24 | 0.22 | 0.30 | 0.24 | 0.44 | 0.32 | 0.23 | 0.31 | 0.32 | 0.32 | 0.33 | 0.25 | 0.24 | 0.29 | 0.29 | 0.36 | 0.18 | 0.34 | 0.28 |
| S6 | X6.6.9.2.6 | 0.19 | 0.23 | 0.27 | 0.29 | 0.32 | 0.23 | 0.25 | 0.28 | 0.28 | 0.19 | 0.18 | 0.15 | 0.08 | 0.21 | 0.35 | 0.27 | 0.15 | 0.22 | 0.23 | 0.34 |
| S6 | X7.6.9.2.7 | 0.18 | 0.22 | 0.23 | 0.30 | 0.28 | 0.41 | 0.38 | 0.23 | 0.32 | 0.38 | 0.34 | 0.32 | 0.20 | 0.31 | 0.43 | 0.26 | 0.32 | 0.17 | 0.35 | 0.30 |
| S6 | X8.6.9.3.8 | 0.21 | 0.23 | 0.22 | 0.31 | 0.30 | 0.19 | 0.18 | 0.29 | 0.32 | 0.27 | 0.16 | 0.14 | 0.09 | 0.17 | 0.27 | 0.26 | 0.13 | 0.24 | 0.16 | 0.29 |
| S6 | X9.6.9.3.9 | 0.06 | 0.13 | 0.18 | 0.26 | 0.21 | 0.28 | 0.23 | 0.20 | 0.19 | 0.28 | 0.22 | 0.22 | 0.17 | 0.11 | 0.20 | 0.18 | 0.22 | 0.17 | 0.25 | 0.22 |
| S6 | X10.6.9.3.10 | 0.24 | 0.33 | 0.35 | 0.37 | 0.44 | 0.39 | 0.33 | 0.36 | 0.42 | 0.39 | 0.30 | 0.33 | 0.30 | 0.24 | 0.37 | 0.35 | 0.25 | 0.33 | 0.36 | 0.39 |
| S10 | X39.10.1.1.1 | 0.30 | 0.41 | 0.33 | 0.40 | 0.46 | 0.68 | 0.59 | 0.42 | 0.41 | 0.59 | 0.54 | 0.64 | 0.67 | 0.24 | 0.46 | 0.35 | 0.63 | 0.42 | 0.63 | 0.40 |
| S10 | X40.10.1.1.2 | 0.48 | 0.56 | 0.49 | 0.68 | 0.66 | 0.64 | 0.61 | 0.63 | 0.63 | 0.62 | 0.62 | 0.60 | 0.60 | 0.44 | 0.72 | 0.52 | 0.60 | 0.54 | 0.65 | 0.60 |
| S10 | X42.10.1.1.4 | 0.47 | 0.55 | 0.36 | 0.53 | 0.55 | 0.75 | 0.64 | 0.60 | 0.60 | 0.69 | 0.72 | 0.67 | 0.71 | 0.41 | 0.65 | 0.46 | 0.74 | 0.49 | 0.76 | 0.47 |
| S10 | X45.10.1.2.8 | 0.44 | 0.51 | 0.41 | 0.61 | 0.60 | 0.67 | 0.60 | 0.60 | 0.60 | 0.46 | 0.62 | 0.69 | 0.50 | 0.39 | 0.61 | 0.51 | 0.61 | 0.53 | 0.56 | 0.41 |
| S10 | X46.10.1.2.7 | 0.31 | 0.41 | 0.31 | 0.56 | 0.57 | 0.83 | 0.70 | 0.52 | 0.59 | 0.71 | 0.71 | 0.75 | 0.79 | 0.21 | 0.56 | 0.42 | 0.60 | 0.52 | 0.71 | 0.39 |
| S10 | X47.10.1.2.9 | 0.22 | 0.41 | 0.33 | 0.51 | 0.47 | 0.73 | 0.62 | 0.58 | 0.54 | 0.58 | 0.65 | 0.75 | 0.72 | 0.36 | 0.59 | 0.36 | 0.79 | 0.34 | 0.78 | 0.35 |
| S10 | X48.10.1.2.10 | 0.33 | 0.51 | 0.44 | 0.64 | 0.52 | 0.61 | 0.57 | 0.53 | 0.53 | 0.50 | 0.54 | 0.60 | 0.50 | 0.36 | 0.51 | 0.44 | 0.61 | 0.46 | 0.59 | 0.51 |
| S10 | X50.10.2.1.2 | 0.20 | 0.38 | 0.31 | 0.46 | 0.56 | 0.72 | 0.61 | 0.54 | 0.46 | 0.61 | 0.68 | 0.62 | 0.68 | 0.35 | 0.51 | 0.34 | 0.63 | 0.36 | 0.67 | 0.30 |
| S10 | X51.10.2.1.3 | 1.00 | 0.52 | 0.49 | 0.48 | 0.45 | 0.25 | 0.40 | 0.45 | 0.59 | 0.35 | 0.41 | 0.27 | 0.26 | 0.40 | 0.57 | 0.60 | 0.33 | 0.59 | 0.33 | 0.54 |
| S10 | X52.10.2.1.4 | 0.52 | 1.00 | 0.54 | 0.64 | 0.60 | 0.49 | 0.52 | 0.67 | 0.62 | 0.42 | 0.49 | 0.50 | 0.49 | 0.44 | 0.67 | 0.58 | 0.60 | 0.63 | 0.37 | 0.49 |
| S10 | X53.10.2.2.5 | 0.49 | 0.54 | 1.00 | 0.65 | 0.43 | 0.32 | 0.40 | 0.49 | 0.48 | 0.34 | 0.42 | 0.31 | 0.38 | 0.37 | 0.46 | 0.69 | 0.37 | 0.59 | 0.38 | 0.51 |
| S10 | X54.10.2.3.6 | 0.48 | 0.64 | 0.65 | 1.00 | 0.54 | 0.56 | 0.59 | 0.66 | 0.70 | 0.51 | 0.53 | 0.47 | 0.51 | 0.39 | 0.60 | 0.77 | 0.51 | 0.62 | 0.47 | 0.64 |
| S10 | X55.10.2.3.7 | 0.45 | 0.60 | 0.43 | 0.54 | 1.00 | 0.55 | 0.45 | 0.66 | 0.66 | 0.51 | 0.61 | 0.51 | 0.53 | 0.47 | 0.71 | 0.57 | 0.51 | 0.66 | 0.55 | 0.47 |
| S10 | X56.10.2.3.8 | 0.25 | 0.49 | 0.32 | 0.56 | 0.55 | 1.00 | 0.67 | 0.59 | 0.56 | 0.68 | 0.71 | 0.85 | 0.76 | 0.28 | 0.57 | 0.46 | 0.73 | 0.44 | 0.72 | 0.40 |
| S10 | X57.10.2.3.9 | 0.40 | 0.52 | 0.40 | 0.59 | 0.45 | 0.67 | 1.00 | 0.52 | 0.55 | 0.51 | 0.53 | 0.64 | 0.52 | 0.33 | 0.60 | 0.51 | 0.62 | 0.45 | 0.65 | 0.42 |
| S10 | X58.10.2.3.10 | 0.45 | 0.67 | 0.49 | 0.66 | 0.66 | 0.59 | 0.52 | 1.00 | 0.69 | 0.33 | 0.55 | 0.59 | 0.61 | 0.49 | 0.71 | 0.65 | 0.63 | 0.59 | 0.57 | 0.48 |
| S10 | X59.10.3.1.1 | 0.59 | 0.62 | 0.48 | 0.70 | 0.66 | 0.56 | 0.55 | 0.69 | 1.00 | 0.51 | 0.59 | 0.59 | 0.51 | 0.52 | 0.79 | 0.69 | 0.53 | 0.71 | 0.61 | 0.59 |
| S10 | X60.10.3.1.2 | 0.35 | 0.42 | 0.34 | 0.51 | 0.51 | 0.68 | 0.51 | 0.33 | 0.51 | 1.00 | 0.61 | 0.54 | 0.63 | 0.43 | 0.52 | 0.46 | 0.56 | 0.44 | 0.58 | 0.49 |
| S10 | X61.10.3.2.3 | 0.41 | 0.49 | 0.42 | 0.53 | 0.61 | 0.71 | 0.53 | 0.55 | 0.59 | 0.61 | 1.00 | 0.64 | 0.61 | 0.44 | 0.63 | 0.49 | 0.64 | 0.61 | 0.63 | 0.38 |
| S10 | X63.10.3.2.5 | 0.27 | 0.50 | 0.31 | 0.47 | 0.51 | 0.85 | 0.64 | 0.59 | 0.59 | 0.54 | 0.64 | 1.00 | 0.72 | 0.26 | 0.61 | 0.41 | 0.82 | 0.43 | 0.77 | 0.38 |
| S10 | X66.10.3.3.8 | 0.26 | 0.49 | 0.38 | 0.51 | 0.53 | 0.76 | 0.52 | 0.61 | 0.51 | 0.63 | 0.61 | 0.72 | 1.00 | 0.29 | 0.52 | 0.45 | 0.66 | 0.43 | 0.64 | 0.38 |
| S10 | X67.10.5.1.1 | 0.40 | 0.44 | 0.37 | 0.39 | 0.47 | 0.28 | 0.33 | 0.49 | 0.52 | 0.43 | 0.44 | 0.26 | 0.29 | 1.00 | 0.60 | 0.51 | 0.39 | 0.42 | 0.39 | 0.40 |
| S10 | X69.10.5.1.3 | 0.57 | 0.67 | 0.46 | 0.60 | 0.71 | 0.57 | 0.60 | 0.71 | 0.79 | 0.52 | 0.63 | 0.61 | 0.52 | 0.60 | 1.00 | 0.66 | 0.62 | 0.72 | 0.62 | 0.49 |
| S10 | X72.10.5.4.6 | 0.60 | 0.58 | 0.69 | 0.77 | 0.57 | 0.46 | 0.51 | 0.65 | 0.69 | 0.46 | 0.49 | 0.41 | 0.45 | 0.51 | 0.66 | 1.00 | 0.44 | 0.75 | 0.48 | 0.55 |
| S10 | X73.10.5.4.7 | 0.33 | 0.60 | 0.37 | 0.51 | 0.51 | 0.73 | 0.62 | 0.63 | 0.53 | 0.56 | 0.64 | 0.82 | 0.66 | 0.39 | 0.62 | 0.44 | 1.00 | 0.39 | 0.71 | 0.48 |
| S10 | X74.10.5.4.8 | 0.59 | 0.63 | 0.59 | 0.62 | 0.66 | 0.44 | 0.45 | 0.59 | 0.71 | 0.44 | 0.61 | 0.43 | 0.43 | 0.42 | 0.72 | 0.75 | 0.39 | 1.00 | 0.43 | 0.49 |
| S10 | X75.10.6.1.1 | 0.33 | 0.37 | 0.38 | 0.47 | 0.55 | 0.72 | 0.65 | 0.57 | 0.61 | 0.58 | 0.63 | 0.77 | 0.64 | 0.39 | 0.62 | 0.48 | 0.71 | 0.43 | 1.00 | 0.45 |
| S10 | X76.10.6.1.2 | 0.54 | 0.49 | 0.51 | 0.64 | 0.47 | 0.40 | 0.42 | 0.48 | 0.59 | 0.49 | 0.38 | 0.38 | 0.38 | 0.40 | 0.49 | 0.55 | 0.48 | 0.49 | 0.45 | 1.00 |
| S10 | X78.10.6.1.4 | 0.60 | 0.63 | 0.65 | 0.71 | 0.54 | 0.46 | 0.52 | 0.66 | 0.66 | 0.40 | 0.53 | 0.47 | 0.48 | 0.46 | 0.64 | 0.65 | 0.51 | 0.63 | 0.47 | 0.56 |
| S10 | X79.10.6.1.5 | 0.31 | 0.44 | 0.35 | 0.48 | 0.47 | 0.67 | 0.49 | 0.59 | 0.48 | 0.60 | 0.59 | 0.63 | 0.66 | 0.27 | 0.46 | 0.34 | 0.69 | 0.40 | 0.71 | 0.46 |
| S10 | X80.10.6.2.6 | 0.23 | 0.39 | 0.22 | 0.40 | 0.57 | 0.83 | 0.48 | 0.55 | 0.55 | 0.67 | 0.74 | 0.76 | 0.73 | 0.37 | 0.56 | 0.36 | 0.70 | 0.39 | 0.75 | 0.33 |
| S10 | X81.10.6.2.7 | 0.54 | 0.55 | 0.44 | 0.56 | 0.68 | 0.60 | 0.46 | 0.58 | 0.69 | 0.68 | 0.66 | 0.53 | 0.62 | 0.49 | 0.70 | 0.51 | 0.56 | 0.61 | 0.61 | 0.56 |
| S10 | X82.10.6.2.8 | 0.47 | 0.42 | 0.41 | 0.47 | 0.54 | 0.73 | 0.70 | 0.49 | 0.59 | 0.64 | 0.70 | 0.77 | 0.62 | 0.38 | 0.57 | 0.52 | 0.68 | 0.50 | 0.77 | 0.50 |
| S10 | X83.10.7.1.1 | 0.37 | 0.56 | 0.31 | 0.57 | 0.69 | 0.70 | 0.64 | 0.56 | 0.63 | 0.65 | 0.68 | 0.60 | 0.56 | 0.47 | 0.75 | 0.49 | 0.60 | 0.54 | 0.62 | 0.44 |
| S10 | X84.10.7.1.2 | 0.15 | 0.35 | 0.10 | 0.26 | 0.57 | 0.64 | 0.43 | 0.47 | 0.43 | 0.55 | 0.60 | 0.64 | 0.63 | 0.35 | 0.48 | 0.25 | 0.58 | 0.33 | 0.58 | 0.23 |
| S10 | X85.10.7.1.3 | 0.59 | 0.57 | 0.51 | 0.66 | 0.43 | 0.34 | 0.43 | 0.51 | 0.62 | 0.46 | 0.33 | 0.29 | 0.29 | 0.39 | 0.55 | 0.67 | 0.28 | 0.65 | 0.25 | 0.63 |
| S10 | X86.10.7.1.4 | 0.32 | 0.56 | 0.40 | 0.61 | 0.57 | 0.81 | 0.61 | 0.70 | 0.63 | 0.61 | 0.74 | 0.76 | 0.76 | 0.38 | 0.65 | 0.47 | 0.78 | 0.53 | 0.72 | 0.40 |
| S10 | X88.10.7.1.6 | 0.34 | 0.56 | 0.42 | 0.52 | 0.54 | 0.77 | 0.63 | 0.59 | 0.51 | 0.67 | 0.68 | 0.72 | 0.69 | 0.37 | 0.56 | 0.45 | 0.81 | 0.48 | 0.68 | 0.49 |
| S10 | X89.10.7.1.7 | 0.46 | 0.70 | 0.55 | 0.75 | 0.66 | 0.60 | 0.64 | 0.76 | 0.80 | 0.46 | 0.53 | 0.56 | 0.59 | 0.46 | 0.64 | 0.63 | 0.60 | 0.57 | 0.55 | 0.59 |
| S10 | X90.10.7.1.8 | 0.29 | 0.50 | 0.33 | 0.44 | 0.49 | 0.76 | 0.55 | 0.57 | 0.57 | 0.59 | 0.59 | 0.80 | 0.69 | 0.33 | 0.55 | 0.39 | 0.82 | 0.34 | 0.71 | 0.45 |
| S10 | X91.10.7.2.9 | 0.50 | 0.63 | 0.58 | 0.75 | 0.57 | 0.47 | 0.46 | 0.66 | 0.70 | 0.53 | 0.38 | 0.41 | 0.52 | 0.53 | 0.65 | 0.74 | 0.45 | 0.60 | 0.41 | 0.60 |
| S10 | X92.10.7.2.10 | 0.43 | 0.59 | 0.44 | 0.66 | 0.70 | 0.53 | 0.49 | 0.61 | 0.72 | 0.53 | 0.54 | 0.46 | 0.51 | 0.36 | 0.63 | 0.51 | 0.47 | 0.60 | 0.51 | 0.61 |

| SITE | IND | X78.10.6.1.4 | X79.10.6.1.5 | X80.10.6.2.6 | X81.10.6.2.7 | X82.10.6.2.8 | X83.10.7.1.1 | X84.10.7.1.2 | X85.10.7.1.3 | X86.10.7.1.4 | X88.10.7.1.6 | X89.10.7.1.7 | X90.10.7.1.8 | X91.10.7.2.9 | X92.10.7.2.10 |
|---|---|---|---|---|---|---|---|---|---|---|---|---|---|---|---|
| S2 | X11.2.2.1.1 | 0.34 | 0.25 | 0.35 | 0.37 | 0.38 | 0.41 | 0.20 | 0.21 | 0.32 | 0.31 | 0.34 | 0.23 | 0.25 | 0.28 |
| S2 | X12.2.2.1.2 | 0.28 | 0.29 | 0.26 | 0.27 | 0.31 | 0.36 | 0.21 | 0.15 | 0.30 | 0.32 | 0.29 | 0.24 | 0.20 | 0.26 |
| S2 | X13.2.2.1.3 | 0.51 | 0.40 | 0.38 | 0.44 | 0.50 | 0.48 | 0.25 | 0.25 | 0.43 | 0.40 | 0.50 | 0.33 | 0.37 | 0.43 |
| S2 | X14.2.2.2.4 | 0.33 | 0.22 | 0.29 | 0.41 | 0.39 | 0.39 | 0.20 | 0.24 | 0.28 | 0.31 | 0.35 | 0.21 | 0.33 | 0.37 |
| S2 | X15.2.2.2.5 | 0.38 | 0.26 | 0.27 | 0.35 | 0.31 | 0.36 | 0.14 | 0.21 | 0.30 | 0.26 | 0.28 | 0.18 | 0.26 | 0.28 |
| S2 | X17.2.2.2.7 | 0.45 | 0.10 | 0.08 | 0.27 | 0.14 | 0.31 | ### | 0.38 | 0.21 | 0.14 | 0.39 | 0.03 | 0.42 | 0.41 |
| S2 | X18.2.2.2.8 | 0.29 | 0.27 | 0.34 | 0.38 | 0.34 | 0.47 | 0.19 | 0.26 | 0.30 | 0.30 | 0.35 | 0.21 | 0.27 | 0.31 |
| S2 | X19.2.2.2.9 | 0.33 | 0.30 | 0.31 | 0.39 | 0.35 | 0.37 | 0.22 | 0.16 | 0.34 | 0.30 | 0.32 | 0.24 | 0.27 | 0.35 |
| S2 | X21.2.5.2.1 | 0.29 | 0.27 | 0.31 | 0.33 | 0.37 | 0.45 | 0.26 | 0.17 | 0.30 | 0.36 | 0.32 | 0.26 | 0.21 | 0.26 |
| S2 | X23.2.5.2.3 | 0.36 | 0.22 | 0.26 | 0.28 | 0.26 | 0.34 | 0.14 | 0.15 | 0.31 | 0.25 | 0.27 | 0.14 | 0.19 | 0.26 |
| S2 | X25.2.5.3.5 | 0.32 | 0.25 | 0.28 | 0.38 | 0.31 | 0.36 | 0.14 | 0.14 | 0.28 | 0.25 | 0.27 | 0.17 | 0.22 | 0.32 |
| S2 | X28.2.5.1.8 | 0.33 | 0.18 | 0.18 | 0.36 | 0.27 | 0.40 | 0.10 | 0.32 | 0.22 | 0.21 | 0.31 | 0.10 | 0.41 | 0.38 |
| S2 | X30.2.5.1.10 | 0.26 | 0.10 | 0.14 | 0.18 | 0.10 | 0.23 | ### | 0.14 | 0.08 | ### | 0.30 | 0.05 | 0.27 | 0.23 |
| S2 | X31.2.6.1.1 | 0.42 | 0.33 | 0.37 | 0.47 | 0.40 | 0.46 | 0.18 | 0.28 | 0.40 | 0.36 | 0.35 | 0.24 | 0.33 | 0.38 |
| S2 | X32.2.6.1.2 | 0.34 | 0.21 | 0.28 | 0.31 | 0.30 | 0.38 | 0.16 | 0.14 | 0.29 | 0.26 | 0.28 | 0.18 | 0.23 | 0.24 |
| S2 | X33.2.6.1.3 | 0.38 | 0.24 | 0.26 | 0.30 | 0.31 | 0.38 | 0.14 | 0.12 | 0.27 | 0.27 | 0.26 | 0.16 | 0.18 | 0.26 |
| S2 | X35.2.6.1.5 | 0.43 | 0.35 | 0.41 | 0.44 | 0.42 | 0.53 | 0.30 | 0.27 | 0.41 | 0.41 | 0.38 | 0.29 | 0.35 | 0.37 |
| S2 | X36.2.6.2.6 | 0.26 | 0.21 | 0.28 | 0.29 | 0.25 | 0.36 | 0.13 | 0.17 | 0.24 | 0.21 | 0.26 | 0.13 | 0.21 | 0.20 |
| S2 | X38.2.6.2.8 | 0.31 | 0.23 | 0.24 | 0.34 | 0.30 | 0.35 | 0.15 | 0.21 | 0.23 | 0.23 | 0.27 | 0.17 | 0.22 | 0.29 |
| S6 | X1.6.9.1.1 | 0.26 | 0.26 | 0.24 | 0.20 | 0.25 | 0.21 | 0.03 | 0.12 | 0.26 | 0.29 | 0.24 | 0.12 | 0.18 | 0.21 |
| S6 | X2.6.9.1.2 | 0.23 | 0.26 | 0.21 | 0.18 | 0.22 | 0.19 | 0.02 | 0.11 | 0.23 | 0.26 | 0.19 | 0.13 | 0.13 | 0.18 |
| S6 | X4.6.9.1.4 | 0.27 | 0.30 | 0.37 | 0.26 | 0.37 | 0.30 | 0.13 | 0.19 | 0.35 | 0.40 | 0.36 | 0.32 | 0.27 | 0.20 |
| S6 | X6.6.9.2.6 | 0.29 | 0.19 | 0.18 | 0.24 | 0.25 | 0.30 | ### | 0.24 | 0.19 | 0.19 | 0.28 | 0.13 | 0.29 | 0.25 |
| S6 | X7.6.9.2.7 | 0.30 | 0.25 | 0.35 | 0.31 | 0.40 | 0.37 | 0.16 | 0.23 | 0.31 | 0.34 | 0.30 | 0.27 | 0.27 | 0.32 |
| S6 | X8.6.9.3.8 | 0.25 | 0.20 | 0.13 | 0.27 | 0.21 | 0.21 | 0.03 | 0.31 | 0.14 | 0.14 | 0.32 | 0.09 | 0.39 | 0.28 |
| S6 | X9.6.9.3.9 | 0.29 | 0.26 | 0.25 | 0.23 | 0.25 | 0.27 | 0.04 | 0.13 | 0.29 | 0.29 | 0.22 | 0.11 | 0.16 | 0.19 |
| S6 | X10.6.9.3.10 | 0.34 | 0.32 | 0.33 | 0.36 | 0.38 | 0.35 | 0.19 | 0.43 | 0.32 | 0.29 | 0.42 | 0.25 | 0.40 | 0.40 |
| S10 | X39.10.1.1.1 | 0.49 | 0.64 | 0.63 | 0.51 | 0.64 | 0.52 | 0.48 | 0.32 | 0.68 | 0.74 | 0.51 | 0.66 | 0.36 | 0.47 |
| S10 | X40.10.1.1.2 | 0.64 | 0.54 | 0.54 | 0.80 | 0.63 | 0.67 | 0.47 | 0.53 | 0.67 | 0.63 | 0.63 | 0.55 | 0.61 | 0.65 |
| S10 | X42.10.1.1.4 | 0.57 | 0.78 | 0.81 | 0.70 | 0.74 | 0.68 | 0.50 | 0.38 | 0.82 | 0.78 | 0.61 | 0.71 | 0.44 | 0.53 |
| S10 | X45.10.1.2.8 | 0.60 | 0.47 | 0.54 | 0.48 | 0.63 | 0.64 | 0.57 | 0.49 | 0.67 | 0.60 | 0.54 | 0.55 | 0.44 | 0.53 |
| S10 | X46.10.1.2.7 | 0.45 | 0.69 | 0.76 | 0.62 | 0.72 | 0.67 | 0.59 | 0.37 | 0.77 | 0.73 | 0.56 | 0.64 | 0.43 | 0.55 |
| S10 | X47.10.1.2.9 | 0.55 | 0.72 | 0.72 | 0.59 | 0.69 | 0.60 | 0.58 | 0.24 | 0.80 | 0.72 | 0.56 | 0.76 | 0.38 | 0.45 |
| S10 | X48.10.1.2.10 | 0.60 | 0.57 | 0.47 | 0.54 | 0.54 | 0.58 | 0.43 | 0.42 | 0.64 | 0.63 | 0.60 | 0.52 | 0.48 | 0.56 |
| S10 | X50.10.2.1.2 | 0.50 | 0.61 | 0.68 | 0.65 | 0.65 | 0.63 | 0.54 | 0.19 | 0.73 | 0.72 | 0.59 | 0.57 | 0.41 | 0.51 |
| S10 | X51.10.2.1.3 | 0.60 | 0.31 | 0.23 | 0.54 | 0.47 | 0.37 | 0.15 | 0.59 | 0.32 | 0.34 | 0.46 | 0.29 | 0.50 | 0.43 |
| S10 | X52.10.2.1.4 | 0.63 | 0.44 | 0.39 | 0.55 | 0.42 | 0.56 | 0.35 | 0.57 | 0.56 | 0.56 | 0.70 | 0.50 | 0.63 | 0.59 |
| S10 | X53.10.2.2.5 | 0.65 | 0.35 | 0.22 | 0.44 | 0.41 | 0.31 | 0.10 | 0.51 | 0.40 | 0.42 | 0.55 | 0.33 | 0.58 | 0.44 |
| S10 | X54.10.2.3.6 | 0.71 | 0.48 | 0.40 | 0.56 | 0.47 | 0.57 | 0.26 | 0.66 | 0.61 | 0.52 | 0.75 | 0.44 | 0.75 | 0.66 |
| S10 | X55.10.2.3.7 | 0.54 | 0.47 | 0.57 | 0.68 | 0.54 | 0.69 | 0.57 | 0.43 | 0.57 | 0.54 | 0.66 | 0.49 | 0.57 | 0.70 |
| S10 | X56.10.2.3.8 | 0.46 | 0.67 | 0.83 | 0.60 | 0.73 | 0.70 | 0.64 | 0.34 | 0.81 | 0.77 | 0.60 | 0.76 | 0.47 | 0.53 |
| S10 | X57.10.2.3.9 | 0.52 | 0.49 | 0.48 | 0.46 | 0.70 | 0.64 | 0.43 | 0.43 | 0.61 | 0.63 | 0.64 | 0.55 | 0.46 | 0.49 |
| S10 | X58.10.2.3.10 | 0.66 | 0.59 | 0.55 | 0.58 | 0.49 | 0.56 | 0.47 | 0.51 | 0.70 | 0.59 | 0.76 | 0.57 | 0.66 | 0.61 |
| S10 | X59.10.3.1.1 | 0.66 | 0.48 | 0.55 | 0.69 | 0.59 | 0.63 | 0.43 | 0.62 | 0.63 | 0.51 | 0.80 | 0.57 | 0.70 | 0.72 |
| S10 | X60.10.3.1.2 | 0.40 | 0.60 | 0.67 | 0.68 | 0.64 | 0.65 | 0.55 | 0.46 | 0.61 | 0.67 | 0.46 | 0.59 | 0.53 | 0.53 |
| S10 | X61.10.3.2.3 | 0.53 | 0.59 | 0.74 | 0.66 | 0.70 | 0.68 | 0.60 | 0.33 | 0.74 | 0.68 | 0.53 | 0.59 | 0.38 | 0.54 |
| S10 | X63.10.3.2.5 | 0.47 | 0.63 | 0.76 | 0.53 | 0.77 | 0.60 | 0.64 | 0.29 | 0.76 | 0.72 | 0.56 | 0.80 | 0.41 | 0.46 |
| S10 | X66.10.3.3.8 | 0.48 | 0.66 | 0.73 | 0.62 | 0.62 | 0.56 | 0.63 | 0.29 | 0.76 | 0.69 | 0.59 | 0.69 | 0.52 | 0.51 |
| S10 | X67.10.5.1.1 | 0.46 | 0.27 | 0.37 | 0.49 | 0.38 | 0.47 | 0.35 | 0.39 | 0.38 | 0.37 | 0.46 | 0.33 | 0.53 | 0.36 |
| S10 | X69.10.5.1.3 | 0.64 | 0.46 | 0.56 | 0.70 | 0.57 | 0.75 | 0.48 | 0.55 | 0.65 | 0.56 | 0.64 | 0.55 | 0.65 | 0.63 |
| S10 | X72.10.5.4.6 | 0.65 | 0.34 | 0.36 | 0.51 | 0.52 | 0.49 | 0.25 | 0.67 | 0.47 | 0.45 | 0.63 | 0.39 | 0.74 | 0.51 |
| S10 | X73.10.5.4.7 | 0.51 | 0.69 | 0.70 | 0.56 | 0.68 | 0.60 | 0.58 | 0.28 | 0.78 | 0.81 | 0.60 | 0.82 | 0.45 | 0.47 |
| S10 | X74.10.5.4.8 | 0.63 | 0.40 | 0.39 | 0.61 | 0.50 | 0.54 | 0.33 | 0.65 | 0.53 | 0.48 | 0.57 | 0.34 | 0.60 | 0.60 |
| S10 | X75.10.6.1.1 | 0.47 | 0.71 | 0.75 | 0.61 | 0.77 | 0.62 | 0.58 | 0.25 | 0.72 | 0.68 | 0.55 | 0.71 | 0.41 | 0.51 |
| S10 | X76.10.6.1.2 | 0.56 | 0.46 | 0.33 | 0.56 | 0.50 | 0.44 | 0.23 | 0.63 | 0.40 | 0.49 | 0.59 | 0.45 | 0.60 | 0.61 |
| S10 | X78.10.6.1.4 | 1.00 | 0.45 | 0.34 | 0.59 | 0.50 | 0.47 | 0.24 | 0.55 | 0.57 | 0.49 | 0.70 | 0.41 | 0.56 | 0.59 |
| S10 | X79.10.6.1.5 | 0.45 | 1.00 | 0.73 | 0.56 | 0.63 | 0.54 | 0.52 | 0.33 | 0.77 | 0.76 | 0.49 | 0.76 | 0.36 | 0.49 |
| S10 | X80.10.6.2.6 | 0.34 | 0.73 | 1.00 | 0.63 | 0.66 | 0.70 | 0.63 | 0.22 | 0.80 | 0.73 | 0.46 | 0.73 | 0.36 | 0.46 |
| S10 | X81.10.6.2.7 | 0.59 | 0.56 | 0.63 | 1.00 | 0.62 | 0.65 | 0.46 | 0.49 | 0.66 | 0.62 | 0.62 | 0.57 | 0.63 | 0.67 |
| S10 | X82.10.6.2.8 | 0.50 | 0.63 | 0.66 | 0.62 | 1.00 | 0.60 | 0.60 | 0.36 | 0.63 | 0.72 | 0.56 | 0.71 | 0.44 | 0.49 |
| S10 | X83.10.7.1.1 | 0.47 | 0.54 | 0.70 | 0.65 | 0.60 | 1.00 | 0.57 | 0.40 | 0.70 | 0.63 | 0.57 | 0.58 | 0.48 | 0.65 |
| S10 | X84.10.7.1.2 | 0.24 | 0.52 | 0.63 | 0.46 | 0.60 | 0.57 | 1.00 | 0.18 | 0.57 | 0.59 | 0.41 | 0.65 | 0.30 | 0.52 |
| S10 | X85.10.7.1.3 | 0.55 | 0.33 | 0.22 | 0.49 | 0.36 | 0.40 | 0.18 | 1.00 | 0.38 | 0.37 | 0.52 | 0.31 | 0.67 | 0.55 |
| S10 | X86.10.7.1.4 | 0.57 | 0.77 | 0.80 | 0.66 | 0.63 | 0.70 | 0.57 | 0.38 | 1.00 | 0.83 | 0.63 | 0.73 | 0.50 | 0.59 |
| S10 | X88.10.7.1.6 | 0.49 | 0.76 | 0.73 | 0.62 | 0.72 | 0.63 | 0.59 | 0.37 | 0.83 | 1.00 | 0.60 | 0.76 | 0.46 | 0.55 |
| S10 | X89.10.7.1.7 | 0.70 | 0.49 | 0.46 | 0.62 | 0.56 | 0.57 | 0.41 | 0.52 | 0.63 | 0.60 | 1.00 | 0.57 | 0.74 | 0.74 |
| S10 | X90.10.7.1.8 | 0.41 | 0.76 | 0.73 | 0.57 | 0.71 | 0.58 | 0.65 | 0.31 | 0.73 | 0.76 | 0.57 | 1.00 | 0.45 | 0.47 |
| S10 | X91.10.7.2.9 | 0.56 | 0.36 | 0.36 | 0.63 | 0.44 | 0.48 | 0.30 | 0.67 | 0.50 | 0.46 | 0.74 | 0.45 | 1.00 | 0.66 |
| S10 | X92.10.7.2.10 | 0.59 | 0.49 | 0.46 | 0.67 | 0.49 | 0.65 | 0.52 | 0.55 | 0.59 | 0.55 | 0.74 | 0.47 | 0.66 | 1.00 |

Table S4. Site pairwise FST matrix calculated using Cockerham and Wei (1987).

| SITE | S1 | S2 | S3 | S4 | S5 | S6 | S7 | S8 | S9 | S10 | S11 | S12 | S13 | S14 | S15 |
|---|---|---|---|---|---|---|---|---|---|---|---|---|---|---|---|
| S1 | 0.000 | 0.040 | 0.284 | 0.233 | 0.133 | 0.216 | 0.190 | 0.153 | 0.125 | 0.125 | 0.309 | 0.136 | 0.176 | 0.166 | 0.129 |
| S2 | 0.040 | 0.000 | 0.324 | 0.238 | 0.176 | 0.179 | 0.195 | 0.148 | 0.148 | 0.192 | 0.321 | 0.109 | 0.187 | 0.177 | 0.177 |
| S3 | 0.284 | 0.324 | 0.000 | 0.290 | 0.212 | 0.245 | 0.248 | 0.219 | 0.201 | 0.232 | 0.354 | 0.171 | 0.258 | 0.301 | 0.289 |
| S4 | 0.233 | 0.238 | 0.290 | 0.000 | 0.074 | 0.085 | 0.214 | 0.144 | 0.053 | 0.168 | 0.297 | 0.137 | 0.186 | 0.222 | 0.220 |
| S5 | 0.133 | 0.176 | 0.212 | 0.074 | 0.000 | 0.150 | 0.139 | 0.057 | 0.122 | 0.022 | 0.261 | 0.238 | 0.124 | 0.113 | 0.142 |
| S6 | 0.216 | 0.179 | 0.245 | 0.085 | 0.150 | 0.000 | 0.158 | 0.107 | 0.012 | 0.016 | 0.236 | 0.082 | 0.123 | 0.164 | 0.205 |
| S7 | 0.190 | 0.195 | 0.248 | 0.214 | 0.139 | 0.158 | 0.000 | 0.138 | 0.117 | 0.180 | 0.269 | 0.125 | 0.184 | 0.189 | 0.207 |
| S8 | 0.153 | 0.148 | 0.219 | 0.144 | 0.057 | 0.107 | 0.138 | 0.000 | 0.112 | 0.147 | 0.253 | 0.036 | 0.165 | 0.166 | 0.151 |
| S9 | 0.125 | 0.148 | 0.201 | 0.053 | 0.122 | 0.012 | 0.117 | 0.112 | 0.000 | 0.011 | 0.227 | 0.082 | 0.116 | 0.126 | 0.138 |
| S10 | 0.125 | 0.192 | 0.232 | 0.168 | 0.022 | 0.016 | 0.180 | 0.147 | 0.011 | 0.000 | 0.160 | 0.019 | 0.170 | 0.182 | 0.160 |
| S11 | 0.309 | 0.321 | 0.354 | 0.297 | 0.261 | 0.236 | 0.269 | 0.253 | 0.227 | 0.160 | 0.000 | 0.157 | 0.255 | 0.300 | 0.301 |
| S12 | 0.136 | 0.109 | 0.171 | 0.137 | 0.238 | 0.082 | 0.125 | 0.036 | 0.082 | 0.019 | 0.157 | 0.000 | 0.085 | 0.127 | 0.146 |
| S13 | 0.176 | 0.187 | 0.258 | 0.186 | 0.124 | 0.123 | 0.184 | 0.165 | 0.116 | 0.170 | 0.255 | 0.085 | 0.000 | 0.189 | 0.192 |
| S14 | 0.166 | 0.177 | 0.301 | 0.222 | 0.113 | 0.164 | 0.189 | 0.166 | 0.126 | 0.182 | 0.300 | 0.127 | 0.189 | 0.000 | 0.178 |
| S15 | 0.129 | 0.177 | 0.289 | 0.220 | 0.142 | 0.205 | 0.207 | 0.151 | 0.138 | 0.160 | 0.301 | 0.146 | 0.192 | 0.178 | 0.000 |

Table S5. Correlation coefficent between all of the sampled ramets within each grid.

### S3 - GRID

| | S3_G_0_0 | S3_G_0_0.5 | S3_G_0_1.5 | S3_G_0_2 | S3_G_0_2.5 | S3_G_0_3 | S3_G_0_3.5 | S3_G_0.5_0 | S3_G_0.5_0.5 | S3_G_0.5_1 | S3_G_0.5_1.5 | S3_G_0.5_2.5 | S3_G_0.5_3 | S3_G_0.5_3.5 | S3_G_1_0 | S3_G_1_1 | S3_G_1_1.5 | S3_G_1_3 | S3_G_1_4 | S3_G_1.5_0 | S3_G_1.5_1.5 | S3_G_1.5_2 | S3_G_1.5_2.5 |
|---|---|---|---|---|---|---|---|---|---|---|---|---|---|---|---|---|---|---|---|---|---|---|---|
| S3_G_0_0 | 1.00 | 0.83 | 0.78 | 0.74 | 0.70 | 0.67 | 0.82 | 0.78 | 0.80 | 0.79 | 0.83 | 0.80 | 0.78 | 0.78 | 0.82 | 0.64 | 0.88 | 0.74 | 0.78 | 0.78 | 0.78 | 0.78 | 0.82 |
| S3_G_0_0.5 | 0.83 | 1.00 | 0.92 | 0.82 | 0.77 | 0.74 | 0.97 | 0.92 | 0.89 | 0.74 | 0.85 | 0.89 | 0.92 | 0.83 | 0.97 | 0.62 | 0.93 | 0.70 | 0.92 | 0.92 | 0.92 | 0.92 | 0.87 |
| S3_G_0_1.5 | 0.78 | 0.92 | 1.00 | 0.88 | 0.94 | 0.89 | 0.94 | 1.00 | 0.97 | 0.79 | 0.92 | 0.97 | 1.00 | 0.89 | 0.94 | 0.81 | 0.88 | 0.88 | 1.00 | 1.00 | 1.00 | 1.00 | 0.94 |
| S3_G_0_2 | 0.74 | 0.82 | 0.88 | 1.00 | 0.78 | 0.74 | 0.78 | 0.88 | 0.90 | 0.62 | 0.93 | 0.90 | 0.88 | 0.88 | 0.78 | 0.90 | 0.83 | 0.83 | 0.88 | 0.88 | 0.88 | 0.88 | 0.93 |
| S3_G_0_2.5 | 0.70 | 0.77 | 0.94 | 0.78 | 1.00 | 0.94 | 0.87 | 0.94 | 0.94 | 0.83 | 0.87 | 0.94 | 0.94 | 0.82 | 0.87 | 0.85 | 0.78 | 0.93 | 0.94 | 0.94 | 0.94 | 0.94 | 0.87 |
| S3_G_0_3 | 0.67 | 0.74 | 0.89 | 0.74 | 0.94 | 1.00 | 0.82 | 0.89 | 0.89 | 0.79 | 0.83 | 0.89 | 0.89 | 0.78 | 0.82 | 0.81 | 0.74 | 0.88 | 0.89 | 0.89 | 0.89 | 0.89 | 0.82 |
| S3_G_0_3.5 | 0.82 | 0.97 | 0.94 | 0.78 | 0.87 | 0.82 | 1.00 | 0.94 | 0.94 | 0.83 | 0.87 | 0.94 | 0.94 | 0.82 | 1.00 | 0.66 | 0.93 | 0.78 | 0.94 | 0.94 | 0.94 | 0.94 | 0.87 |
| S3_G_0.5_0 | 0.78 | 0.92 | 1.00 | 0.88 | 0.94 | 0.89 | 0.94 | 1.00 | 0.97 | 0.79 | 0.92 | 0.97 | 1.00 | 0.89 | 0.94 | 0.81 | 0.88 | 0.88 | 1.00 | 1.00 | 1.00 | 1.00 | 0.94 |
| S3_G_0.5_0.5 | 0.80 | 0.89 | 0.97 | 0.90 | 0.94 | 0.89 | 0.94 | 0.97 | 1.00 | 0.81 | 0.97 | 1.00 | 0.97 | 0.89 | 0.94 | 0.87 | 0.90 | 0.90 | 0.97 | 0.97 | 0.97 | 0.97 | 0.94 |
| S3_G_0.5_1 | 0.79 | 0.74 | 0.79 | 0.62 | 0.83 | 0.79 | 0.83 | 0.79 | 0.81 | 1.00 | 0.83 | 0.81 | 0.79 | 0.79 | 0.83 | 0.65 | 0.88 | 0.88 | 0.79 | 0.79 | 0.79 | 0.79 | 0.83 |
| S3_G_0.5_1.5 | 0.83 | 0.85 | 0.92 | 0.93 | 0.87 | 0.83 | 0.87 | 0.92 | 0.97 | 0.83 | 1.00 | 0.97 | 0.92 | 0.92 | 0.87 | 0.90 | 0.93 | 0.93 | 0.92 | 0.92 | 0.92 | 0.92 | 0.97 |
| S3_G_0.5_2.5 | 0.80 | 0.89 | 0.97 | 0.90 | 0.94 | 0.89 | 0.94 | 0.97 | 1.00 | 0.81 | 0.97 | 1.00 | 0.97 | 0.89 | 0.94 | 0.87 | 0.90 | 0.90 | 0.97 | 0.97 | 0.97 | 0.97 | 0.94 |
| S3_G_0.5_3 | 0.78 | NA | 1.00 | 0.88 | 0.94 | 0.89 | 0.94 | 1.00 | 0.97 | 0.79 | 0.92 | 0.97 | 1.00 | 0.89 | 0.94 | 0.81 | 0.88 | 0.88 | 1.00 | 1.00 | 1.00 | 1.00 | 0.94 |
| S3_G_0.5_3.5 | 0.78 | 0.83 | 0.89 | 0.88 | 0.82 | 0.78 | 0.82 | 0.89 | 0.89 | 0.79 | 0.92 | 0.89 | 0.89 | 1.00 | 0.82 | 0.81 | 0.88 | 0.88 | 0.89 | 0.89 | 0.89 | 0.89 | 0.94 |
| S3_G_1_0 | 0.82 | 0.97 | 0.94 | 0.78 | 0.87 | 0.82 | 1.00 | 0.94 | 0.94 | 0.83 | 0.87 | 0.94 | 0.94 | 0.82 | 1.00 | 0.66 | 0.93 | 0.78 | 0.94 | 0.94 | 0.94 | 0.94 | 0.87 |
| S3_G_1_1 | 0.64 | 0.62 | 0.81 | 0.90 | 0.85 | 0.81 | 0.66 | 0.81 | 0.87 | 0.65 | 0.90 | 0.87 | 0.81 | 0.81 | 0.66 | 1.00 | 0.69 | 0.90 | 0.81 | 0.81 | 0.81 | 0.81 | 0.85 |
| S3_G_1_1.5 | 0.88 | 0.93 | 0.88 | 0.83 | 0.78 | 0.74 | 0.93 | 0.88 | 0.90 | 0.88 | 0.93 | 0.90 | 0.88 | 0.88 | 0.93 | 0.69 | 1.00 | 0.83 | 0.88 | 0.88 | 0.88 | 0.88 | 0.93 |
| S3_G_1_3 | 0.74 | 0.70 | 0.88 | 0.83 | 0.93 | 0.88 | 0.78 | 0.88 | 0.90 | 0.88 | 0.93 | 0.90 | 0.88 | 0.88 | 0.78 | 0.90 | 0.83 | 1.00 | 0.88 | 0.88 | 0.88 | 0.88 | 0.93 |
| S3_G_1_4 | 0.78 | 0.92 | 1.00 | 0.88 | 0.94 | 0.89 | 0.94 | 1.00 | 0.97 | 0.79 | 0.92 | 0.97 | 1.00 | 0.89 | 0.94 | 0.81 | 0.88 | 0.88 | 1.00 | 1.00 | 1.00 | 1.00 | 0.94 |
| S3_G_1.5_0 | 0.78 | 0.92 | 1.00 | 0.88 | 0.94 | 0.89 | 0.94 | 1.00 | 0.97 | 0.79 | 0.92 | 0.97 | 1.00 | 0.89 | 0.94 | 0.81 | 0.88 | 0.88 | 1.00 | 1.00 | 1.00 | 1.00 | 0.94 |
| S3_G_1.5_1.5 | 0.78 | 0.92 | 1.00 | 0.88 | 0.94 | 0.89 | 0.94 | 1.00 | 0.97 | 0.79 | 0.92 | 0.97 | 1.00 | 0.89 | 0.94 | 0.81 | 0.88 | 0.88 | 1.00 | 1.00 | 1.00 | 1.00 | 0.94 |
| S3_G_1.5_2 | 0.78 | 0.92 | 1.00 | 0.88 | 0.94 | 0.89 | 0.94 | 1.00 | 0.97 | 0.79 | 0.92 | 0.97 | 1.00 | 0.89 | 0.94 | 0.81 | 0.88 | 0.88 | 1.00 | 1.00 | 1.00 | 1.00 | 0.94 |
| S3_G_1.5_2.5 | 0.82 | 0.87 | 0.94 | 0.93 | 0.87 | 0.82 | 0.87 | 0.94 | 0.94 | 0.83 | 0.97 | 0.94 | 0.94 | 0.94 | 0.87 | 0.85 | 0.93 | 0.93 | 0.94 | 0.94 | 0.94 | 0.94 | 1.00 |
| S3_G_1.5_3 | 0.72 | 0.75 | 0.89 | 0.90 | 0.84 | 0.80 | 0.75 | 0.89 | 0.87 | 0.73 | 0.89 | 0.87 | 0.89 | 0.89 | 0.75 | 0.87 | 0.79 | 0.90 | 0.89 | 0.89 | 0.89 | 0.89 | 0.94 |
| S3_G_1.5_4 | 0.82 | 0.87 | 0.94 | 0.93 | 0.87 | 0.82 | 0.87 | 0.94 | 0.94 | 0.83 | 0.97 | 0.94 | 0.94 | 0.94 | 0.87 | 0.85 | 0.93 | 0.93 | 0.94 | 0.94 | 0.94 | 0.94 | 1.00 |
| S3_G_1.5_4.5 | 0.87 | 0.94 | 0.87 | 0.78 | 0.74 | 0.71 | 0.91 | 0.87 | 0.85 | 0.87 | 0.87 | 0.85 | 0.87 | 0.87 | 0.91 | 0.60 | 0.98 | 0.78 | 0.87 | 0.87 | 0.87 | 0.87 | 0.91 |
| S3_G_2_0.5 | 0.78 | 0.92 | 1.00 | 0.88 | 0.94 | 0.89 | 0.94 | 1.00 | 0.97 | 0.79 | 0.92 | 0.97 | 1.00 | 0.89 | 0.94 | 0.81 | 0.88 | 0.88 | 1.00 | 1.00 | 1.00 | 1.00 | 0.94 |
| S3_G_2_1.5 | 0.82 | 0.87 | 0.94 | 0.93 | 0.87 | 0.82 | 0.87 | 0.94 | 0.94 | 0.83 | 0.97 | 0.94 | 0.94 | 0.94 | 0.87 | 0.85 | 0.93 | 0.93 | 0.94 | 0.94 | 0.94 | 0.94 | 1.00 |
| S3_G_2_2.5 | 0.75 | 0.88 | 0.95 | 0.84 | 0.90 | 0.85 | 0.90 | 0.95 | 0.93 | 0.76 | 0.88 | 0.93 | 0.95 | 0.85 | 0.90 | 0.78 | 0.84 | 0.84 | 0.95 | 0.95 | 0.95 | 0.95 | 0.90 |
| S3_G_2_3 | 0.75 | 0.88 | 0.95 | 0.84 | 0.90 | 0.85 | 0.90 | 0.95 | 0.93 | 0.76 | 0.88 | 0.93 | 0.95 | 0.95 | 0.90 | 0.78 | 0.84 | 0.84 | 0.95 | 0.95 | 0.95 | 0.95 | 0.90 |
| S3_G_2_4 | 0.80 | 0.97 | 0.97 | 0.90 | 0.84 | 0.80 | 0.94 | 0.97 | 0.93 | 0.73 | 0.89 | 0.93 | 0.97 | 0.89 | 0.94 | 0.74 | 0.90 | 0.79 | 0.97 | 0.97 | 0.97 | 0.97 | 0.94 |
| S3_G_2.5_0 | 0.43 | 0.44 | 0.43 | 0.40 | 0.45 | 0.43 | 0.45 | 0.43 | 0.42 | 0.43 | 0.40 | 0.42 | 0.43 | 0.38 | 0.45 | 0.42 | 0.40 | 0.40 | 0.43 | 0.43 | 0.43 | 0.43 | 0.39 |
| S3_G_2.5_1 | 0.70 | 0.77 | 0.94 | 0.78 | 1.00 | 0.94 | 0.87 | 0.94 | 0.94 | 0.83 | 0.87 | 0.94 | 0.94 | 0.82 | 0.87 | 0.85 | 0.78 | 0.93 | 0.94 | 0.94 | 0.94 | 0.94 | 0.87 |
| S3_G_2.5_1.5 | 0.39 | 0.41 | 0.39 | 0.42 | 0.41 | 0.45 | 0.41 | 0.39 | 0.40 | 0.44 | 0.41 | 0.40 | 0.39 | 0.45 | 0.41 | 0.43 | 0.42 | 0.42 | 0.39 | 0.39 | 0.39 | 0.39 | 0.41 |
| S3_G_2.5_3.5 | 0.82 | 0.87 | 0.94 | 0.93 | 0.87 | 0.82 | 0.87 | 0.94 | 0.94 | 0.83 | 0.97 | 0.94 | 0.94 | 0.94 | 0.87 | 0.85 | 0.93 | 0.93 | 0.94 | 0.94 | 0.94 | 0.94 | 1.00 |
| S3_G_2.5_4 | 0.49 | 0.40 | 0.38 | 0.41 | 0.39 | 0.38 | 0.39 | 0.38 | 0.39 | 0.43 | 0.40 | 0.39 | 0.38 | 0.38 | 0.39 | 0.43 | 0.41 | 0.41 | 0.38 | 0.38 | 0.38 | 0.38 | 0.39 |
| S3_G_2.5_4.5 | 0.47 | 0.43 | 0.42 | 0.44 | 0.43 | 0.42 | 0.43 | 0.42 | 0.42 | 0.45 | 0.43 | 0.42 | 0.42 | 0.42 | 0.43 | 0.45 | 0.44 | 0.44 | 0.42 | 0.42 | 0.42 | 0.42 | 0.43 |
| S3_G_3_0 | 0.50 | 0.42 | 0.40 | 0.42 | 0.41 | 0.40 | 0.41 | 0.40 | 0.40 | 0.44 | 0.42 | 0.40 | 0.40 | 0.40 | 0.41 | 0.44 | 0.42 | 0.42 | 0.40 | 0.40 | 0.40 | 0.40 | 0.41 |
| S3_G_3_1.5 | 0.52 | 0.45 | 0.44 | 0.45 | 0.45 | 0.44 | 0.45 | 0.44 | 0.44 | 0.46 | 0.45 | 0.44 | 0.44 | 0.44 | 0.45 | 0.46 | 0.45 | 0.45 | 0.44 | 0.44 | 0.44 | 0.44 | 0.45 |
| S3_G_3_1.5.1 | 0.51 | 0.47 | 0.41 | 0.46 | 0.37 | 0.41 | 0.43 | 0.41 | 0.40 | 0.41 | 0.42 | 0.40 | 0.41 | 0.41 | 0.43 | 0.41 | 0.46 | 0.39 | 0.41 | 0.41 | 0.41 | 0.41 | 0.43 |
| S3_G_3_3 | 0.46 | 0.39 | 0.37 | 0.40 | 0.38 | 0.41 | 0.38 | 0.37 | 0.38 | 0.42 | 0.39 | 0.38 | 0.37 | 0.37 | 0.38 | 0.42 | 0.40 | 0.40 | 0.37 | 0.37 | 0.37 | 0.37 | 0.38 |
| S3_G_3_3.5 | 0.47 | 0.39 | 0.37 | 0.40 | 0.39 | 0.42 | 0.39 | 0.37 | 0.38 | 0.43 | 0.39 | 0.38 | 0.37 | 0.37 | 0.39 | 0.42 | 0.40 | 0.40 | 0.37 | 0.37 | 0.37 | 0.37 | 0.39 |
| S3_G_3_4.5 | 0.47 | 0.39 | 0.37 | 0.40 | 0.39 | 0.47 | 0.39 | 0.37 | 0.38 | 0.43 | 0.39 | 0.38 | 0.37 | 0.37 | 0.39 | 0.42 | 0.40 | 0.40 | 0.37 | 0.37 | 0.37 | 0.37 | 0.39 |
| S3_G_3.5_0.5 | 0.63 | 0.49 | 0.45 | 0.48 | 0.41 | 0.45 | 0.46 | 0.45 | 0.44 | 0.44 | 0.46 | 0.44 | 0.45 | 0.45 | 0.46 | 0.44 | 0.48 | 0.42 | 0.45 | 0.45 | 0.45 | 0.45 | 0.46 |
| S3_G_3.5_1 | 0.55 | 0.41 | 0.39 | 0.41 | 0.40 | 0.39 | 0.40 | 0.39 | 0.39 | 0.43 | 0.41 | 0.39 | 0.39 | 0.39 | 0.40 | 0.43 | 0.41 | 0.41 | 0.39 | 0.39 | 0.39 | 0.39 | 0.40 |
| S3_G_3.5_1.5 | 0.44 | 0.49 | 0.49 | 0.42 | 0.52 | 0.49 | 0.52 | 0.49 | 0.48 | 0.44 | 0.41 | 0.48 | 0.49 | 0.44 | 0.52 | 0.43 | 0.42 | 0.42 | 0.49 | 0.49 | 0.49 | 0.49 | 0.40 |
| S3_G_3.5_2 | 0.48 | 0.49 | 0.43 | 0.47 | 0.39 | 0.43 | 0.45 | 0.43 | 0.43 | 0.43 | 0.44 | 0.43 | 0.43 | 0.48 | 0.45 | 0.43 | 0.47 | 0.41 | 0.43 | 0.43 | 0.43 | 0.43 | 0.45 |
| S3_G_4_3 | 0.44 | 0.45 | 0.44 | 0.41 | 0.45 | 0.44 | 0.45 | 0.44 | 0.43 | 0.43 | 0.40 | 0.43 | 0.44 | 0.38 | 0.45 | 0.43 | 0.41 | 0.41 | 0.44 | 0.44 | 0.44 | 0.44 | 0.40 |
| S3_G_4.5_3.5 | 0.53 | 0.40 | 0.38 | 0.41 | 0.39 | 0.38 | 0.39 | 0.38 | 0.39 | 0.43 | 0.40 | 0.39 | 0.38 | 0.38 | 0.39 | 0.43 | 0.41 | 0.41 | 0.38 | 0.38 | 0.38 | 0.38 | 0.39 |
| S3_G4_4_1 | 0.48 | 0.41 | 0.39 | 0.42 | 0.41 | 0.39 | 0.41 | 0.39 | 0.40 | 0.44 | 0.41 | 0.40 | 0.39 | 0.39 | 0.41 | 0.44 | 0.42 | 0.42 | 0.39 | 0.39 | 0.39 | 0.39 | 0.41 |

| | S3_G_1.5_3 | S3_G_1.5_4 | S3_G_1.5_4.5 | S3_G_2_0.5 | S3_G_2_1.5 | S3_G_2_2.5 | S3_G_2_3 | S3_G_2_4 | S3_G_2.5_0 | S3_G_2.5_1 | S3_G_2.5_1.5 | S3_G_2.5_3.5 | S3_G_2.5_4 | S3_G_2.5_4.5 | S3_G_3_0 | S3_G_3_1.5 | S3_G_3_1.5.1 | S3_G_3_3 | S3_G_3_3.5 | S3_G_3_4.5 | S3_G_3.5_0.5 | S3_G_3.5_1 | S3_G_3.5_1.5 | S3_G_3.5_2 | S3_G_4_3 | S3_G_4.5_3.5 | S3_G4_4_1 |
|---|---|---|---|---|---|---|---|---|---|---|---|---|---|---|---|---|---|---|---|---|---|---|---|---|---|---|---|
| S3_G_0_0 | 0.72 | 0.82 | 0.87 | 0.78 | 0.82 | 0.75 | 0.75 | 0.80 | 0.43 | 0.70 | 0.39 | 0.82 | 0.49 | 0.47 | 0.50 | 0.52 | 0.51 | 0.46 | 0.47 | 0.47 | 0.63 | 0.55 | 0.44 | 0.48 | 0.44 | 0.53 | 0.48 |
| S3_G_0_0.5 | 0.75 | 0.87 | 0.94 | 0.92 | 0.87 | 0.88 | 0.88 | 0.97 | 0.44 | 0.77 | 0.41 | 0.87 | 0.40 | 0.43 | 0.42 | 0.45 | 0.47 | 0.39 | 0.39 | 0.39 | 0.49 | 0.41 | 0.49 | 0.49 | 0.45 | 0.40 | 0.41 |
| S3_G_0_1.5 | 0.89 | 0.94 | 0.87 | 1.00 | 0.94 | 0.95 | 0.95 | 0.97 | 0.43 | 0.94 | 0.39 | 0.94 | 0.38 | 0.42 | 0.40 | 0.44 | 0.41 | 0.37 | 0.37 | 0.37 | 0.45 | 0.39 | 0.49 | 0.43 | 0.44 | 0.38 | 0.39 |
| S3_G_0_2 | 0.90 | 0.93 | 0.78 | 0.88 | 0.93 | 0.84 | 0.84 | 0.90 | 0.40 | 0.78 | 0.42 | 0.93 | 0.41 | 0.44 | 0.42 | 0.45 | 0.46 | 0.40 | 0.40 | 0.40 | 0.48 | 0.41 | 0.42 | 0.47 | 0.41 | 0.41 | 0.42 |
| S3_G_0_2.5 | 0.84 | 0.87 | 0.74 | 0.94 | 0.87 | 0.90 | 0.90 | 0.84 | 0.45 | 1.00 | 0.41 | 0.87 | 0.39 | 0.43 | 0.41 | 0.45 | 0.37 | 0.38 | 0.39 | 0.39 | 0.41 | 0.40 | 0.52 | 0.39 | 0.45 | 0.39 | 0.41 |
| S3_G_0_3 | 0.80 | 0.82 | 0.71 | 0.89 | 0.82 | 0.85 | 0.85 | 0.80 | 0.43 | 0.94 | 0.45 | 0.82 | 0.38 | 0.42 | 0.40 | 0.44 | 0.41 | 0.41 | 0.42 | 0.47 | 0.45 | 0.39 | 0.49 | 0.43 | 0.44 | 0.38 | 0.39 |
| S3_G_0_3.5 | 0.75 | 0.87 | 0.91 | 0.94 | 0.87 | 0.90 | 0.90 | 0.94 | 0.45 | 0.87 | 0.41 | 0.87 | 0.39 | 0.43 | 0.41 | 0.45 | 0.43 | 0.38 | 0.39 | 0.39 | 0.46 | 0.40 | 0.52 | 0.45 | 0.45 | 0.39 | 0.41 |
| S3_G_0.5_0 | 0.89 | 0.94 | 0.87 | 1.00 | 0.94 | 0.95 | 0.95 | 0.97 | 0.43 | 0.94 | 0.39 | 0.94 | 0.38 | 0.42 | 0.40 | 0.44 | 0.41 | 0.37 | 0.37 | 0.37 | 0.45 | 0.39 | 0.49 | 0.43 | 0.44 | 0.38 | 0.39 |
| S3_G_0.5_0.5 | 0.87 | 0.94 | 0.85 | 0.97 | 0.94 | 0.93 | 0.93 | 0.93 | 0.42 | 0.94 | 0.40 | 0.94 | 0.39 | 0.42 | 0.40 | 0.44 | 0.40 | 0.38 | 0.38 | 0.38 | 0.44 | 0.39 | 0.48 | 0.43 | 0.43 | 0.39 | 0.40 |
| S3_G_0.5_1 | 0.73 | 0.83 | 0.87 | 0.79 | 0.83 | 0.76 | 0.76 | 0.73 | 0.43 | 0.83 | 0.44 | 0.83 | 0.43 | 0.45 | 0.44 | 0.46 | 0.41 | 0.42 | 0.43 | 0.43 | 0.44 | 0.43 | 0.44 | 0.43 | 0.43 | 0.43 | 0.44 |
| S3_G_0.5_1.5 | 0.89 | 0.97 | 0.87 | 0.92 | 0.97 | 0.88 | 0.88 | 0.89 | 0.40 | 0.87 | 0.41 | 0.97 | 0.40 | 0.43 | 0.42 | 0.45 | 0.42 | 0.39 | 0.39 | 0.39 | 0.46 | 0.41 | 0.41 | 0.44 | 0.40 | 0.40 | 0.41 |
| S3_G_0.5_2.5 | 0.87 | 0.94 | 0.85 | 0.97 | 0.94 | 0.93 | 0.93 | 0.93 | 0.42 | 0.94 | 0.40 | 0.94 | 0.39 | 0.42 | 0.40 | 0.44 | 0.40 | 0.38 | 0.38 | 0.38 | 0.44 | 0.39 | 0.48 | 0.43 | 0.43 | 0.39 | 0.40 |
| S3_G_0.5_3 | 0.89 | 0.94 | 0.87 | 1.00 | 0.94 | 0.95 | 0.95 | 0.97 | 0.43 | 0.94 | 0.39 | 0.94 | 0.38 | 0.42 | 0.40 | 0.44 | 0.41 | 0.37 | 0.37 | 0.37 | 0.45 | 0.39 | 0.49 | 0.43 | 0.44 | 0.38 | 0.39 |
| S3_G_0.5_3.5 | 0.89 | 0.94 | 0.87 | 0.89 | 0.94 | 0.85 | 0.95 | 0.89 | 0.38 | 0.82 | 0.45 | 0.94 | 0.38 | 0.42 | 0.40 | 0.44 | 0.41 | 0.37 | 0.37 | 0.37 | 0.45 | 0.39 | 0.44 | 0.48 | 0.38 | 0.38 | 0.39 |
| S3_G_1_0 | 0.75 | 0.87 | 0.91 | 0.94 | 0.87 | 0.90 | 0.90 | 0.94 | 0.45 | 0.87 | 0.41 | 0.87 | 0.39 | 0.43 | 0.41 | 0.45 | 0.43 | 0.38 | 0.39 | 0.39 | 0.46 | 0.40 | 0.52 | 0.45 | 0.45 | 0.39 | 0.41 |
| S3_G_1_1 | 0.87 | 0.85 | 0.60 | 0.81 | 0.85 | 0.78 | 0.78 | 0.74 | 0.42 | 0.85 | 0.43 | 0.85 | 0.43 | 0.45 | 0.44 | 0.46 | 0.41 | 0.42 | 0.42 | 0.42 | 0.44 | 0.43 | 0.43 | 0.43 | 0.43 | 0.43 | 0.44 |
| S3_G_1_1.5 | 0.79 | 0.93 | 0.98 | 0.88 | 0.93 | 0.84 | 0.84 | 0.90 | 0.40 | 0.78 | 0.42 | 0.93 | 0.41 | 0.44 | 0.42 | 0.45 | 0.46 | 0.40 | 0.40 | 0.40 | 0.48 | 0.41 | 0.42 | 0.47 | 0.41 | 0.41 | 0.42 |
| S3_G_1_3 | 0.90 | 0.93 | 0.78 | 0.88 | 0.93 | 0.84 | 0.84 | 0.79 | 0.40 | 0.93 | 0.42 | 0.93 | 0.41 | 0.44 | 0.42 | 0.45 | 0.39 | 0.40 | 0.40 | 0.40 | 0.42 | 0.41 | 0.42 | 0.41 | 0.41 | 0.41 | 0.42 |
| S3_G_1_4 | 0.89 | 0.94 | 0.87 | 1.00 | 0.94 | 0.95 | 0.95 | 0.97 | 0.43 | 0.94 | 0.39 | 0.94 | 0.38 | 0.42 | 0.40 | 0.44 | 0.41 | 0.37 | 0.37 | 0.37 | 0.45 | 0.39 | 0.49 | 0.43 | 0.44 | 0.38 | 0.39 |
| S3_G_1.5_0 | 0.89 | 0.94 | 0.87 | 1.00 | 0.94 | 0.95 | 0.95 | 0.97 | 0.43 | 0.94 | 0.39 | 0.94 | 0.38 | 0.42 | 0.40 | 0.44 | 0.41 | 0.37 | 0.37 | 0.37 | 0.45 | 0.39 | 0.49 | 0.43 | 0.44 | 0.38 | 0.39 |
| S3_G_1.5_1.5 | 0.89 | 0.94 | 0.87 | 1.00 | 0.94 | 0.95 | 0.95 | 0.97 | 0.43 | 0.94 | 0.39 | 0.94 | 0.38 | 0.42 | 0.40 | 0.44 | 0.41 | 0.37 | 0.37 | 0.37 | 0.45 | 0.39 | 0.49 | 0.43 | 0.44 | 0.38 | 0.39 |
| S3_G_1.5_2 | 0.89 | 0.94 | 0.87 | 1.00 | 0.94 | 0.95 | 0.95 | 0.97 | 0.43 | 0.94 | 0.39 | 0.94 | 0.38 | 0.42 | 0.40 | 0.44 | 0.41 | 0.37 | 0.37 | 0.37 | 0.45 | 0.39 | 0.49 | 0.43 | 0.44 | 0.38 | 0.39 |
| S3_G_1.5_2.5 | 0.94 | 1.00 | 0.91 | 0.94 | 1.00 | 0.90 | 0.90 | 0.94 | 0.39 | 0.87 | 0.41 | 1.00 | 0.39 | 0.43 | 0.41 | 0.45 | 0.43 | 0.38 | 0.39 | 0.39 | 0.46 | 0.40 | 0.40 | 0.45 | 0.40 | 0.39 | 0.41 |
| S3_G_1.5_3 | 1.00 | 0.94 | 0.78 | 0.89 | 0.94 | 0.85 | 0.85 | 0.87 | 0.38 | 0.84 | 0.40 | 0.94 | 0.43 | 0.42 | 0.44 | 0.51 | 0.44 | 0.41 | 0.42 | 0.42 | 0.44 | 0.39 | 0.40 | 0.47 | 0.39 | 0.43 | 0.43 |
| S3_G_1.5_4 | 0.94 | 1.00 | 0.91 | 0.94 | 1.00 | 0.90 | 0.90 | 0.94 | 0.39 | 0.87 | 0.41 | 1.00 | 0.39 | 0.43 | 0.41 | 0.45 | 0.43 | 0.38 | 0.39 | 0.39 | 0.46 | 0.40 | 0.40 | 0.45 | 0.40 | 0.39 | 0.41 |
| S3_G_1.5_4.5 | 0.78 | 0.91 | 1.00 | 0.87 | 0.91 | 0.83 | 0.83 | 0.91 | 0.41 | 0.74 | 0.42 | 0.91 | 0.41 | 0.44 | 0.43 | 0.46 | 0.47 | 0.40 | 0.41 | 0.41 | 0.50 | 0.42 | 0.42 | 0.49 | 0.41 | 0.41 | 0.42 |
| S3_G_2_0.5 | 0.89 | 0.94 | 0.87 | 1.00 | 0.94 | 0.95 | 0.95 | 0.97 | 0.43 | 0.94 | 0.39 | 0.94 | 0.38 | 0.42 | 0.40 | 0.44 | 0.41 | 0.37 | 0.37 | 0.37 | 0.45 | 0.39 | 0.49 | 0.43 | 0.44 | 0.38 | 0.39 |
| S3_G_2_1.5 | 0.94 | 1.00 | 0.91 | 0.94 | 1.00 | 0.90 | 0.90 | 0.94 | 0.39 | 0.87 | 0.41 | 1.00 | 0.39 | 0.43 | 0.41 | 0.45 | 0.43 | 0.38 | 0.39 | 0.39 | 0.46 | 0.40 | 0.40 | 0.45 | 0.40 | 0.39 | 0.41 |
| S3_G_2_2.5 | 0.85 | 0.90 | 0.83 | 0.95 | 0.90 | 1.00 | 0.91 | 0.93 | 0.41 | 0.90 | 0.43 | 0.90 | 0.37 | 0.41 | 0.39 | 0.43 | 0.44 | 0.36 | 0.41 | 0.41 | 0.43 | 0.42 | 0.47 | 0.46 | 0.52 | 0.37 | 0.38 |
| S3_G_2_3 | 0.85 | 0.90 | 0.83 | 0.95 | 0.90 | 0.91 | 1.00 | 0.93 | 0.41 | 0.90 | 0.43 | 0.90 | 0.37 | 0.41 | 0.39 | 0.43 | 0.39 | 0.36 | 0.36 | 0.36 | 0.43 | 0.37 | 0.52 | 0.46 | 0.42 | 0.37 | 0.38 |
| S3_G_2_4 | 0.87 | 0.94 | 0.91 | 0.97 | 0.94 | 0.93 | 0.93 | 1.00 | 0.42 | 0.84 | 0.40 | 0.94 | 0.39 | 0.42 | 0.40 | 0.44 | 0.44 | 0.38 | 0.38 | 0.38 | 0.48 | 0.39 | 0.48 | 0.47 | 0.43 | 0.39 | 0.40 |
| S3_G_2.5_0 | 0.38 | 0.39 | 0.41 | 0.43 | 0.39 | 0.41 | 0.41 | 0.42 | 1.00 | 0.45 | 0.66 | 0.39 | 0.61 | 0.51 | 0.65 | 0.61 | 0.53 | 0.57 | 0.52 | 0.52 | 0.57 | 0.63 | 0.70 | 0.61 | 0.59 | 0.69 | 0.69 |
| S3_G_2.5_1 | 0.84 | 0.87 | 0.74 | 0.94 | 0.87 | 0.90 | 0.90 | 0.84 | 0.45 | 1.00 | 0.41 | 0.87 | 0.39 | 0.43 | 0.41 | 0.45 | 0.37 | 0.38 | 0.39 | 0.39 | 0.41 | 0.40 | 0.52 | 0.39 | 0.45 | 0.39 | 0.41 |
| S3_G_2.5_1.5 | 0.40 | 0.41 | 0.42 | 0.39 | 0.41 | 0.43 | 0.43 | 0.40 | 0.66 | 0.41 | 1.00 | 0.41 | 0.59 | 0.58 | 0.65 | 0.52 | 0.55 | 0.55 | 0.61 | 0.58 | 0.48 | 0.64 | 0.61 | 0.59 | 0.60 | 0.53 | 0.54 |
| S3_G_2.5_3.5 | 0.94 | 1.00 | 0.91 | 0.94 | 1.00 | 0.90 | 0.90 | 0.94 | 0.39 | 0.87 | 0.41 | 1.00 | 0.39 | 0.43 | 0.41 | 0.45 | 0.43 | 0.38 | 0.39 | 0.39 | 0.46 | 0.40 | 0.40 | 0.45 | 0.40 | 0.39 | 0.41 |
| S3_G_2.5_4 | 0.43 | 0.39 | 0.41 | 0.38 | 0.39 | 0.37 | 0.37 | 0.39 | 0.61 | 0.39 | 0.59 | 0.39 | 1.00 | 0.70 | 0.82 | 0.64 | 0.61 | 0.60 | 0.61 | 0.61 | 0.65 | 0.70 | 0.59 | 0.46 | 0.58 | 0.73 | 0.67 |
| S3_G_2.5_4.5 | 0.42 | 0.43 | 0.44 | 0.42 | 0.43 | 0.41 | 0.41 | 0.42 | 0.51 | 0.43 | 0.58 | 0.43 | 0.70 | 1.00 | 0.72 | 0.63 | 0.73 | 0.72 | 0.76 | 0.74 | 0.58 | 0.59 | 0.56 | 0.42 | 0.55 | 0.71 | 0.69 |
| S3_G_3_0 | 0.44 | 0.41 | 0.43 | 0.40 | 0.41 | 0.39 | 0.39 | 0.40 | 0.65 | 0.41 | 0.65 | 0.41 | 0.82 | 0.72 | 1.00 | 0.63 | 0.58 | 0.57 | 0.60 | 0.58 | 0.59 | 0.61 | 0.62 | 0.51 | 0.65 | 0.71 | 0.67 |
| S3_G_3_1.5 | 0.51 | 0.45 | 0.46 | 0.44 | 0.45 | 0.43 | 0.43 | 0.44 | 0.61 | 0.45 | 0.52 | 0.45 | 0.64 | 0.63 | 0.63 | 1.00 | 0.62 | 0.59 | 0.63 | 0.63 | 0.61 | 0.59 | 0.62 | 0.49 | 0.58 | 0.68 | 0.65 |
| S3_G_3_1.5.1 | 0.44 | 0.43 | 0.47 | 0.41 | 0.43 | 0.44 | 0.39 | 0.44 | 0.53 | 0.37 | 0.55 | 0.43 | 0.61 | 0.73 | 0.58 | 0.62 | 1.00 | 0.94 | 0.94 | 0.94 | 0.57 | 0.60 | 0.57 | 0.57 | 0.56 | 0.61 | 0.60 |
| S3_G_3_3 | 0.41 | 0.38 | 0.40 | 0.37 | 0.38 | 0.36 | 0.36 | 0.38 | 0.57 | 0.38 | 0.55 | 0.38 | 0.60 | 0.72 | 0.57 | 0.59 | 0.94 | 1.00 | 0.91 | 0.89 | 0.56 | 0.57 | 0.59 | 0.54 | 0.53 | 0.59 | 0.58 |
| S3_G_3_3.5 | 0.42 | 0.39 | 0.41 | 0.37 | 0.39 | 0.41 | 0.36 | 0.38 | 0.52 | 0.39 | 0.61 | 0.39 | 0.61 | 0.76 | 0.60 | 0.63 | 0.94 | 0.91 | 1.00 | 0.91 | 0.53 | 0.58 | 0.53 | 0.51 | 0.59 | 0.59 | 0.56 |
| S3_G_3_4.5 | 0.42 | 0.39 | 0.41 | 0.37 | 0.39 | 0.41 | 0.36 | 0.38 | 0.52 | 0.39 | 0.58 | 0.39 | 0.61 | 0.74 | 0.58 | 0.63 | 0.94 | 0.89 | 0.91 | 1.00 | 0.57 | 0.60 | 0.55 | 0.53 | 0.52 | 0.61 | 0.60 |
| S3_G_3.5_0.5 | 0.44 | 0.46 | 0.50 | 0.45 | 0.46 | 0.43 | 0.43 | 0.48 | 0.57 | 0.41 | 0.48 | 0.46 | 0.65 | 0.58 | 0.59 | 0.61 | 0.57 | 0.56 | 0.53 | 0.57 | 1.00 | 0.60 | 0.62 | 0.62 | 0.50 | 0.69 | 0.64 |
| S3_G_3.5_1 | 0.39 | 0.40 | 0.42 | 0.39 | 0.40 | 0.42 | 0.37 | 0.39 | 0.63 | 0.40 | 0.64 | 0.40 | 0.70 | 0.59 | 0.61 | 0.59 | 0.60 | 0.57 | 0.58 | 0.60 | 0.60 | 1.00 | 0.63 | 0.57 | 0.63 | 0.72 | 0.72 |
| S3_G_3.5_1.5 | 0.40 | 0.40 | 0.42 | 0.49 | 0.40 | 0.47 | 0.52 | 0.48 | 0.70 | 0.52 | 0.61 | 0.40 | 0.59 | 0.56 | 0.62 | 0.62 | 0.57 | 0.59 | 0.53 | 0.55 | 0.62 | 0.63 | 1.00 | 0.61 | 0.62 | 0.62 | 0.63 |
| S3_G_3.5_2 | 0.47 | 0.45 | 0.49 | 0.43 | 0.45 | 0.46 | 0.46 | 0.47 | 0.61 | 0.39 | 0.59 | 0.45 | 0.46 | 0.42 | 0.51 | 0.49 | 0.57 | 0.54 | 0.51 | 0.53 | 0.62 | 0.57 | 0.61 | 1.00 | 0.68 | 0.55 | 0.56 |
| S3_G_4_3 | 0.39 | 0.40 | 0.41 | 0.44 | 0.40 | 0.52 | 0.42 | 0.43 | 0.59 | 0.45 | 0.60 | 0.40 | 0.58 | 0.55 | 0.65 | 0.58 | 0.56 | 0.53 | 0.59 | 0.52 | 0.50 | 0.63 | 0.62 | 0.68 | 1.00 | 0.61 | 0.53 |
| S3_G_4.5_3.5 | 0.43 | 0.39 | 0.41 | 0.38 | 0.39 | 0.37 | 0.37 | 0.39 | 0.69 | 0.39 | 0.53 | 0.39 | 0.73 | 0.71 | 0.71 | 0.68 | 0.61 | 0.59 | 0.59 | 0.61 | 0.69 | 0.72 | 0.62 | 0.55 | 0.61 | 1.00 | 0.94 |
| S3_G4_4_1 | 0.43 | 0.41 | 0.42 | 0.39 | 0.41 | 0.38 | 0.38 | 0.40 | 0.69 | 0.41 | 0.54 | 0.41 | 0.67 | 0.69 | 0.67 | 0.65 | 0.60 | 0.58 | 0.56 | 0.60 | 0.64 | 0.72 | 0.63 | 0.56 | 0.53 | 0.94 | 1.00 |

### S5 - GRID

| | S5_G_.0.5_0.5 | S5_G_.0.5_1 | S5_G_.1_0 | S5_G_.1_3 | S5_G_.1_3.5 | S5_G_.1_3.5.1 | S5_G_.1_4.5 | S5_G_0_0.5 | S5_G_0_1 | S5_G_0_1.5 | S5_G_0_3 | S5_G_0_4.5 | S5_G_0.5_0.5 | S5_G_0.5_1 | S5_G_0.5_1.5 | S5_G_0.5_2 | S5_G_0.5_2.5 | S5_G_0.5_2.5.1 | S5_G_0.5_3 | S5_G_0.5_3.5 | S5_G_0.5_4 | S5_G_1_0 | S5_G_1_0.5 | S5_G_1_1 |
|---|---|---|---|---|---|---|---|---|---|---|---|---|---|---|---|---|---|---|---|---|---|---|---|---|
| S5_G_.0.5_0.5 | 1.00 | 0.87 | 0.98 | 0.97 | 0.95 | 0.94 | 0.91 | 0.92 | 0.98 | 0.97 | 0.97 | 0.96 | 0.90 | 0.86 | 0.95 | 0.96 | 0.97 | 0.97 | 0.95 | 0.93 | 0.95 | 0.95 | 0.94 | 0.98 |
| S5_G_.0.5_1 | 0.87 | 1.00 | 0.88 | 0.89 | 0.78 | 0.85 | 0.82 | 0.82 | 0.88 | 0.90 | 0.90 | 0.90 | 0.90 | 0.90 | 0.83 | 0.90 | 0.85 | 0.85 | 0.88 | 0.93 | 0.91 | 0.86 | 0.88 | 0.86 |
| S5_G_.1_0 | 0.98 | 0.88 | 1.00 | 0.94 | 0.93 | 0.94 | 0.91 | 0.92 | 0.96 | 0.98 | 0.98 | 0.95 | 0.90 | 0.86 | 0.96 | 0.94 | 0.94 | 0.94 | 0.92 | 0.94 | 0.96 | 0.93 | 0.92 | 0.97 |
| S5_G_.1_3 | 0.97 | 0.89 | 0.94 | 1.00 | 0.91 | 0.90 | 0.88 | 0.88 | 0.98 | 0.93 | 0.93 | 0.97 | 0.84 | 0.84 | 0.89 | 0.96 | 0.95 | 0.95 | 0.94 | 0.91 | 0.89 | 0.92 | 0.94 | 0.95 |
| S5_G_.1_3.5 | 0.95 | 0.78 | 0.93 | 0.91 | 1.00 | 0.91 | 0.88 | 0.89 | 0.93 | 0.88 | 0.88 | 0.86 | 0.87 | 0.76 | 0.92 | 0.88 | 0.90 | 0.90 | 0.85 | 0.87 | 0.88 | 0.90 | 0.92 | 0.97 |
| S5_G_.1_3.5.1 | 0.94 | 0.85 | 0.94 | 0.90 | 0.91 | 1.00 | 0.97 | 0.97 | 0.94 | 0.94 | 0.94 | 0.93 | 0.92 | 0.87 | 0.98 | 0.90 | 0.94 | 0.94 | 0.94 | 0.90 | 0.92 | 0.97 | 0.91 | 0.97 |
| S5_G_.1_4.5 | 0.91 | 0.82 | 0.91 | 0.88 | 0.88 | 0.97 | 1.00 | 0.94 | 0.91 | 0.91 | 0.91 | 0.90 | 0.89 | 0.85 | 0.95 | 0.88 | 0.92 | 0.92 | 0.91 | 0.88 | 0.90 | 0.94 | 0.89 | 0.94 |
| S5_G_0_0.5 | 0.92 | 0.82 | 0.92 | 0.88 | 0.89 | 0.97 | 0.94 | 1.00 | 0.92 | 0.92 | 0.92 | 0.88 | 0.90 | 0.84 | 0.95 | 0.88 | 0.91 | 0.91 | 0.92 | 0.88 | 0.89 | 0.96 | 0.90 | 0.96 |
| S5_G_0_1 | 0.98 | 0.88 | 0.96 | 0.98 | 0.93 | 0.94 | 0.91 | 0.92 | 1.00 | 0.95 | 0.95 | 0.95 | 0.86 | 0.86 | 0.91 | 0.94 | 0.98 | 0.98 | 0.96 | 0.94 | 0.91 | 0.93 | 0.92 | 0.97 |
| S5_G_0_1.5 | 0.97 | 0.90 | 0.98 | 0.93 | 0.88 | 0.94 | 0.91 | 0.92 | 0.95 | 1.00 | 1.00 | 0.96 | 0.93 | 0.90 | 0.95 | 0.96 | 0.93 | 0.93 | 0.95 | 0.93 | 0.95 | 0.95 | 0.94 | 0.95 |
| S5_G_0_3 | 0.97 | 0.90 | 0.98 | 0.93 | 0.88 | 0.94 | 0.91 | 0.92 | 0.95 | 1.00 | 1.00 | 0.96 | 0.93 | 0.90 | 0.95 | 0.96 | 0.93 | 0.93 | 0.95 | 0.93 | 0.95 | 0.95 | 0.94 | 0.95 |
| S5_G_0_4.5 | 0.96 | 0.90 | 0.95 | 0.97 | 0.86 | 0.93 | 0.90 | 0.88 | 0.95 | 0.96 | 0.96 | 1.00 | 0.88 | 0.88 | 0.93 | 0.97 | 0.95 | 0.95 | 0.95 | 0.91 | 0.93 | 0.94 | 0.93 | 0.94 |
| S5_G_0.5_0.5 | 0.90 | 0.90 | 0.90 | 0.84 | 0.87 | 0.92 | 0.89 | 0.90 | 0.86 | 0.93 | 0.93 | 0.88 | 1.00 | 0.89 | 0.94 | 0.89 | 0.87 | 0.87 | 0.90 | 0.88 | 0.94 | 0.95 | 0.94 | 0.92 |
| S5_G_0.5_1 | 0.86 | 0.90 | 0.86 | 0.84 | 0.76 | 0.87 | 0.85 | 0.84 | 0.86 | 0.90 | 0.90 | 0.88 | 0.89 | 1.00 | 0.85 | 0.86 | 0.87 | 0.87 | 0.90 | 0.88 | 0.90 | 0.88 | 0.84 | 0.85 |
| S5_G_0.5_1.5 | 0.95 | 0.83 | 0.96 | 0.89 | 0.92 | 0.98 | 0.95 | 0.95 | 0.91 | 0.95 | 0.95 | 0.93 | 0.94 | 0.85 | 1.00 | 0.91 | 0.93 | 0.93 | 0.91 | 0.89 | 0.95 | 0.97 | 0.92 | 0.97 |
| S5_G_0.5_2 | 0.96 | 0.90 | 0.94 | 0.96 | 0.88 | 0.90 | 0.88 | 0.88 | 0.94 | 0.96 | 0.96 | 0.97 | 0.89 | 0.86 | 0.91 | 1.00 | 0.93 | 0.93 | 0.94 | 0.89 | 0.91 | 0.94 | 0.96 | 0.94 |
| S5_G_0.5_2.5 | 0.97 | 0.85 | 0.94 | 0.95 | 0.90 | 0.94 | 0.92 | 0.91 | 0.98 | 0.93 | 0.93 | 0.95 | 0.87 | 0.87 | 0.93 | 0.93 | 1.00 | 1.00 | 0.98 | 0.91 | 0.93 | 0.96 | 0.91 | 0.96 |
| S5_G_0.5_2.5.1 | 0.97 | 0.85 | 0.94 | 0.95 | 0.90 | 0.94 | 0.92 | 0.91 | 0.98 | 0.93 | 0.93 | 0.95 | 0.87 | 0.87 | 0.93 | 0.93 | 1.00 | 1.00 | 0.98 | 0.91 | 0.93 | 0.96 | 0.91 | 0.96 |
| S5_G_0.5_3 | 0.95 | 0.88 | 0.92 | 0.94 | 0.85 | 0.94 | 0.91 | 0.92 | 0.96 | 0.95 | 0.95 | 0.95 | 0.90 | 0.90 | 0.91 | 0.94 | 0.98 | 0.98 | 1.00 | 0.90 | 0.91 | 0.97 | 0.92 | 0.93 |
| S5_G_0.5_3.5 | 0.93 | 0.93 | 0.94 | 0.91 | 0.87 | 0.90 | 0.88 | 0.88 | 0.94 | 0.93 | 0.93 | 0.91 | 0.88 | 0.88 | 0.89 | 0.89 | 0.91 | 0.91 | 0.90 | 1.00 | 0.93 | 0.88 | 0.87 | 0.92 |
| S5_G_0.5_4 | 0.95 | 0.91 | 0.96 | 0.89 | 0.88 | 0.92 | 0.90 | 0.89 | 0.91 | 0.95 | 0.95 | 0.93 | 0.94 | 0.90 | 0.95 | 0.91 | 0.93 | 0.93 | 0.91 | 0.93 | 1.00 | 0.94 | 0.89 | 0.94 |
| S5_G_1_0 | 0.95 | 0.86 | 0.93 | 0.92 | 0.90 | 0.97 | 0.94 | 0.96 | 0.93 | 0.95 | 0.95 | 0.94 | 0.95 | 0.88 | 0.97 | 0.94 | 0.96 | 0.96 | 0.97 | 0.88 | 0.94 | 1.00 | 0.96 | 0.97 |
| S5_G_1_0.5 | 0.94 | 0.88 | 0.92 | 0.94 | 0.92 | 0.91 | 0.89 | 0.90 | 0.92 | 0.94 | 0.94 | 0.93 | 0.94 | 0.84 | 0.92 | 0.96 | 0.91 | 0.91 | 0.92 | 0.87 | 0.89 | 0.96 | 1.00 | 0.96 |
| S5_G_1_1 | 0.98 | 0.86 | 0.97 | 0.95 | 0.97 | 0.97 | 0.94 | 0.96 | 0.97 | 0.95 | 0.95 | 0.94 | 0.92 | 0.85 | 0.97 | 0.94 | 0.96 | 0.96 | 0.93 | 0.92 | 0.94 | 0.97 | 0.96 | 1.00 |
| S5_G_1_1.5 | 0.98 | 0.84 | 0.96 | 0.94 | 0.93 | 0.94 | 0.91 | 0.92 | 0.96 | 0.95 | 0.95 | 0.95 | 0.90 | 0.86 | 0.96 | 0.94 | 0.98 | 0.98 | 0.96 | 0.90 | 0.96 | 0.97 | 0.92 | 0.97 |
| S5_G_1_2 | 0.96 | 0.87 | 0.91 | 0.98 | 0.88 | 0.91 | 0.88 | 0.87 | 0.96 | 0.91 | 0.91 | 0.98 | 0.85 | 0.85 | 0.90 | 0.96 | 0.97 | 0.97 | 0.96 | 0.88 | 0.90 | 0.94 | 0.93 | 0.94 |
| S5_G_1_2.5 | 0.96 | 0.87 | 0.91 | 0.98 | 0.88 | 0.91 | 0.88 | 0.87 | 0.96 | 0.91 | 0.91 | 0.98 | 0.85 | 0.85 | 0.90 | 0.96 | 0.97 | 0.97 | 0.96 | 0.88 | 0.90 | 0.94 | 0.93 | 0.94 |
| S5_G_1_3 | 0.92 | 0.85 | 0.89 | 0.91 | 0.86 | 0.91 | 0.88 | 0.94 | 0.89 | 0.92 | 0.92 | 0.92 | 0.91 | 0.84 | 0.92 | 0.94 | 0.90 | 0.90 | 0.93 | 0.83 | 0.88 | 0.97 | 0.95 | 0.94 |
| S5_G_1_4.5 | 0.93 | 0.85 | 0.94 | 0.91 | 0.83 | 0.90 | 0.88 | 0.88 | 0.90 | 0.93 | 0.93 | 0.97 | 0.84 | 0.84 | 0.93 | 0.93 | 0.91 | 0.91 | 0.90 | 0.87 | 0.93 | 0.92 | 0.87 | 0.92 |
| S5_G_1.5_0 | 0.92 | 0.86 | 0.90 | 0.92 | 0.87 | 0.92 | 0.90 | 0.96 | 0.93 | 0.92 | 0.92 | 0.89 | 0.88 | 0.85 | 0.90 | 0.91 | 0.92 | 0.92 | 0.93 | 0.88 | 0.86 | 0.94 | 0.93 | 0.94 |
| S5_G_1.5_0.5 | 0.97 | 0.87 | 0.95 | 0.93 | 0.88 | 0.94 | 0.91 | 0.92 | 0.95 | 0.97 | 0.97 | 0.96 | 0.93 | 0.90 | 0.95 | 0.96 | 0.97 | 0.97 | 0.98 | 0.89 | 0.95 | 0.98 | 0.94 | 0.95 |
| S5_G_1.5_1.5 | 0.95 | 0.87 | 0.96 | 0.93 | 0.92 | 0.98 | 0.95 | 0.95 | 0.96 | 0.95 | 0.95 | 0.93 | 0.90 | 0.85 | 0.95 | 0.91 | 0.93 | 0.93 | 0.91 | 0.93 | 0.90 | 0.94 | 0.92 | 0.97 |
| S5_G_1.5_2 | 0.98 | 0.88 | 1.00 | 0.94 | 0.93 | 0.94 | 0.91 | 0.92 | 0.96 | 0.98 | 0.98 | 0.95 | 0.90 | 0.86 | 0.96 | 0.94 | 0.94 | 0.94 | 0.92 | 0.94 | 0.96 | 0.93 | 0.92 | 0.97 |
| S5_G_1.5_2.5 | 0.94 | 0.94 | 0.94 | 0.90 | 0.86 | 0.93 | 0.91 | 0.90 | 0.94 | 0.94 | 0.94 | 0.93 | 0.92 | 0.92 | 0.92 | 0.90 | 0.94 | 0.94 | 0.94 | 0.96 | 0.98 | 0.92 | 0.87 | 0.92 |
| S5_G_1.5_3 | 0.94 | 0.85 | 0.94 | 0.90 | 0.86 | 0.93 | 0.91 | 0.90 | 0.94 | 0.94 | 0.94 | 0.93 | 0.87 | 0.87 | 0.92 | 0.90 | 0.94 | 0.94 | 0.94 | 0.90 | 0.92 | 0.92 | 0.87 | 0.92 |
| S5_G_1.5_4 | 0.94 | 0.85 | 0.94 | 0.90 | 0.86 | 0.93 | 0.91 | 0.90 | 0.94 | 0.94 | 0.94 | 0.93 | 0.87 | 0.87 | 0.92 | 0.90 | 0.94 | 0.94 | 0.94 | 0.90 | 0.92 | 0.92 | 0.87 | 0.92 |
| S5_G_1.5_4.5 | 0.95 | 0.87 | 0.96 | 0.93 | 0.92 | 0.98 | 0.95 | 0.95 | 0.96 | 0.95 | 0.95 | 0.93 | 0.90 | 0.85 | 0.95 | 0.91 | 0.93 | 0.93 | 0.91 | 0.93 | 0.90 | 0.94 | 0.92 | 0.97 |
| S5_G_2_1 | 0.93 | 0.78 | 0.90 | 0.87 | 0.95 | 0.90 | 0.88 | 0.93 | 0.90 | 0.89 | 0.89 | 0.85 | 0.92 | 0.80 | 0.93 | 0.89 | 0.91 | 0.91 | 0.90 | 0.83 | 0.89 | 0.95 | 0.94 | 0.95 |
| S5_G_2_3 | 0.42 | 0.45 | 0.42 | 0.42 | 0.42 | 0.40 | 0.42 | 0.41 | 0.42 | 0.42 | 0.42 | 0.42 | 0.43 | 0.46 | 0.41 | 0.42 | 0.42 | 0.42 | 0.42 | 0.42 | 0.43 | 0.41 | 0.42 | 0.41 |
| S5_G_2_3.5 | 0.42 | 0.45 | 0.42 | 0.43 | 0.43 | 0.41 | 0.40 | 0.42 | 0.42 | 0.42 | 0.42 | 0.42 | 0.43 | 0.47 | 0.41 | 0.42 | 0.42 | 0.42 | 0.42 | 0.43 | 0.43 | 0.42 | 0.43 | 0.42 |
| S5_G_2_4 | 0.42 | 0.46 | 0.42 | 0.43 | 0.43 | 0.41 | 0.41 | 0.42 | 0.42 | 0.42 | 0.42 | 0.43 | 0.44 | 0.47 | 0.42 | 0.42 | 0.42 | 0.42 | 0.42 | 0.43 | 0.44 | 0.42 | 0.43 | 0.42 |
| S5_G_2_4.5 | 0.42 | 0.46 | 0.42 | 0.43 | 0.43 | 0.41 | 0.41 | 0.42 | 0.42 | 0.42 | 0.42 | 0.43 | 0.44 | 0.48 | 0.42 | 0.42 | 0.42 | 0.42 | 0.42 | 0.43 | 0.44 | 0.42 | 0.43 | 0.42 |
| S5_G_2.5_0 | 0.43 | 0.46 | 0.43 | 0.43 | 0.45 | 0.44 | 0.45 | 0.45 | 0.43 | 0.43 | 0.43 | 0.43 | 0.45 | 0.47 | 0.44 | 0.43 | 0.43 | 0.43 | 0.43 | 0.43 | 0.44 | 0.44 | 0.45 | 0.44 |
| S5_G_2.5_4 | 0.42 | 0.48 | 0.42 | 0.42 | 0.44 | 0.43 | 0.44 | 0.44 | 0.42 | 0.42 | 0.42 | 0.42 | 0.46 | 0.50 | 0.43 | 0.42 | 0.42 | 0.42 | 0.42 | 0.44 | 0.45 | 0.43 | 0.44 | 0.43 |
| S5_G_3_1 | 0.42 | 0.42 | 0.42 | 0.42 | 0.42 | 0.40 | 0.40 | 0.41 | 0.42 | 0.42 | 0.42 | 0.42 | 0.41 | 0.45 | 0.41 | 0.43 | 0.41 | 0.41 | 0.42 | 0.42 | 0.41 | 0.41 | 0.42 | 0.41 |
| S5_G_3_1.5 | 0.41 | 0.41 | 0.41 | 0.41 | 0.44 | 0.44 | 0.47 | 0.45 | 0.41 | 0.41 | 0.41 | 0.41 | 0.43 | 0.43 | 0.43 | 0.41 | 0.41 | 0.41 | 0.41 | 0.41 | 0.40 | 0.43 | 0.44 | 0.43 |
| S5_G_3_2 | 0.45 | 0.44 | 0.46 | 0.45 | 0.48 | 0.46 | 0.47 | 0.47 | 0.44 | 0.45 | 0.45 | 0.45 | 0.47 | 0.45 | 0.48 | 0.45 | 0.44 | 0.44 | 0.44 | 0.43 | 0.45 | 0.46 | 0.47 | 0.46 |
| S5_G_3_2.5 | 0.46 | 0.44 | 0.46 | 0.45 | 0.46 | 0.44 | 0.43 | 0.44 | 0.44 | 0.46 | 0.46 | 0.45 | 0.45 | 0.47 | 0.46 | 0.45 | 0.44 | 0.44 | 0.44 | 0.43 | 0.46 | 0.45 | 0.46 | 0.45 |
| S5_G_3.5_3.5 | 0.48 | 0.48 | 0.48 | 0.47 | 0.50 | 0.51 | 0.50 | 0.52 | 0.47 | 0.49 | 0.49 | 0.48 | 0.52 | 0.49 | 0.52 | 0.49 | 0.47 | 0.47 | 0.48 | 0.46 | 0.48 | 0.52 | 0.52 | 0.50 |

| | S5_G_1_1.5 | S5_G_1_2 | S5_G_1_2.5 | S5_G_1_3 | S5_G_1_4.5 | S5_G_1.5_0 | S5_G_1.5_0.5 | S5_G_1.5_1.5 | S5_G_1.5_2 | S5_G_1.5_2.5 | S5_G_1.5_3 | S5_G_1.5_4 | S5_G_1.5_4.5 | S5_G_2_1 | S5_G_2_3 | S5_G_2_3.5 | S5_G_2_4 | S5_G_2_4.5 | S5_G_2.5_0 | S5_G_2.5_4 | S5_G_3_1 | S5_G_3_1.5 | S5_G_3_2 | S5_G_3_2.5 | S5_G_3.5_3.5 |
|---|---|---|---|---|---|---|---|---|---|---|---|---|---|---|---|---|---|---|---|---|---|---|---|---|---|
| S5_G_.0.5_0.5 | 0.98 | 0.96 | 0.96 | 0.92 | 0.93 | 0.92 | 0.97 | 0.95 | 0.98 | 0.94 | 0.94 | 0.94 | 0.95 | 0.93 | 0.42 | 0.42 | 0.42 | 0.42 | 0.43 | 0.42 | 0.42 | 0.41 | 0.45 | 0.46 | 0.48 |
| S5_G_.0.5_1 | 0.84 | 0.87 | 0.87 | 0.85 | 0.85 | 0.86 | 0.87 | 0.87 | 0.88 | 0.94 | 0.85 | 0.85 | 0.87 | 0.78 | 0.45 | 0.45 | 0.46 | 0.46 | 0.46 | 0.48 | 0.42 | 0.41 | 0.44 | 0.44 | 0.48 |
| S5_G_.1_0 | 0.96 | 0.91 | 0.91 | 0.89 | 0.94 | 0.90 | 0.95 | 0.96 | 1.00 | 0.94 | 0.94 | 0.94 | 0.96 | 0.90 | 0.42 | 0.42 | 0.42 | 0.42 | 0.43 | 0.42 | 0.42 | 0.41 | 0.46 | 0.46 | 0.48 |
| S5_G_.1_3 | 0.94 | 0.98 | 0.98 | 0.91 | 0.91 | 0.92 | 0.93 | 0.93 | 0.94 | 0.90 | 0.90 | 0.90 | 0.93 | 0.87 | 0.42 | 0.43 | 0.43 | 0.43 | 0.43 | 0.42 | 0.42 | 0.41 | 0.45 | 0.45 | 0.47 |
| S5_G_.1_3.5 | 0.93 | 0.88 | 0.88 | 0.86 | 0.83 | 0.87 | 0.88 | 0.92 | 0.93 | 0.86 | 0.86 | 0.86 | 0.92 | 0.95 | 0.42 | 0.43 | 0.43 | 0.43 | 0.45 | 0.44 | 0.42 | 0.44 | 0.48 | 0.46 | 0.50 |
| S5_G_.1_3.5.1 | 0.94 | 0.91 | 0.91 | 0.91 | 0.90 | 0.92 | 0.94 | 0.98 | 0.94 | 0.93 | 0.93 | 0.93 | 0.98 | 0.90 | 0.40 | 0.41 | 0.41 | 0.41 | 0.44 | 0.43 | 0.40 | 0.44 | 0.46 | 0.44 | 0.51 |
| S5_G_.1_4.5 | 0.91 | 0.88 | 0.88 | 0.88 | 0.88 | 0.90 | 0.91 | 0.95 | 0.91 | 0.91 | 0.91 | 0.91 | 0.95 | 0.88 | 0.42 | 0.40 | 0.41 | 0.41 | 0.45 | 0.44 | 0.40 | 0.47 | 0.47 | 0.43 | 0.50 |
| S5_G_0_0.5 | 0.92 | 0.87 | 0.87 | 0.94 | 0.88 | 0.96 | 0.92 | 0.95 | 0.92 | 0.90 | 0.90 | 0.90 | 0.95 | 0.93 | 0.41 | 0.42 | 0.42 | 0.42 | 0.45 | 0.44 | 0.41 | 0.45 | 0.47 | 0.44 | 0.52 |
| S5_G_0_1 | 0.96 | 0.96 | 0.96 | 0.89 | 0.90 | 0.93 | 0.95 | 0.96 | 0.96 | 0.94 | 0.94 | 0.94 | 0.96 | 0.90 | 0.42 | 0.42 | 0.42 | 0.42 | 0.43 | 0.42 | 0.42 | 0.41 | 0.44 | 0.44 | 0.47 |
| S5_G_0_1.5 | 0.95 | 0.91 | 0.91 | 0.92 | 0.93 | 0.92 | 0.97 | 0.95 | 0.98 | 0.94 | 0.94 | 0.94 | 0.95 | 0.89 | 0.42 | 0.42 | 0.42 | 0.42 | 0.43 | 0.42 | 0.42 | 0.41 | 0.45 | 0.46 | 0.49 |
| S5_G_0_3 | 0.95 | 0.91 | 0.91 | 0.92 | 0.93 | 0.92 | 0.97 | 0.95 | 0.98 | 0.94 | 0.94 | 0.94 | 0.95 | 0.89 | 0.42 | 0.42 | 0.42 | 0.42 | 0.43 | 0.42 | 0.42 | 0.41 | 0.45 | 0.46 | 0.49 |
| S5_G_0_4.5 | 0.95 | 0.98 | 0.98 | 0.92 | 0.97 | 0.89 | 0.96 | 0.93 | 0.95 | 0.93 | 0.93 | 0.93 | 0.93 | 0.85 | 0.42 | 0.42 | 0.43 | 0.43 | 0.43 | 0.42 | 0.42 | 0.41 | 0.45 | 0.45 | 0.48 |
| S5_G_0.5_0.5 | 0.90 | 0.85 | 0.85 | 0.91 | 0.84 | 0.88 | 0.93 | 0.90 | 0.90 | 0.92 | 0.87 | 0.87 | 0.90 | 0.92 | 0.43 | 0.43 | 0.44 | 0.44 | 0.45 | 0.46 | 0.41 | 0.43 | 0.47 | 0.45 | 0.52 |
| S5_G_0.5_1 | 0.86 | 0.85 | 0.85 | 0.84 | 0.84 | 0.85 | 0.90 | 0.85 | 0.86 | 0.92 | 0.87 | 0.87 | 0.85 | 0.80 | 0.46 | 0.47 | 0.47 | 0.48 | 0.47 | 0.50 | 0.45 | 0.43 | 0.45 | 0.47 | 0.49 |
| S5_G_0.5_1.5 | 0.96 | 0.90 | 0.90 | 0.92 | 0.93 | 0.90 | 0.95 | 0.95 | 0.96 | 0.92 | 0.92 | 0.92 | 0.95 | 0.93 | 0.41 | 0.41 | 0.42 | 0.42 | 0.44 | 0.43 | 0.41 | 0.43 | 0.48 | 0.46 | 0.52 |
| S5_G_0.5_2 | 0.94 | 0.96 | 0.96 | 0.94 | 0.93 | 0.91 | 0.96 | 0.91 | 0.94 | 0.90 | 0.90 | 0.90 | 0.91 | 0.89 | 0.42 | 0.42 | 0.42 | 0.42 | 0.43 | 0.42 | 0.43 | 0.41 | 0.45 | 0.45 | 0.49 |
| S5_G_0.5_2.5 | 0.98 | 0.97 | 0.97 | 0.90 | 0.91 | 0.92 | 0.97 | 0.93 | 0.94 | 0.94 | 0.94 | 0.94 | 0.93 | 0.91 | 0.42 | 0.42 | 0.42 | 0.42 | 0.43 | 0.42 | 0.41 | 0.41 | 0.44 | 0.44 | 0.47 |
| S5_G_0.5_2.5.1 | 0.98 | 0.97 | 0.97 | 0.90 | 0.91 | 0.92 | 0.97 | 0.93 | 0.94 | 0.94 | 0.94 | 0.94 | 0.93 | 0.91 | 0.42 | 0.42 | 0.42 | 0.42 | 0.43 | 0.42 | 0.41 | 0.41 | 0.44 | 0.44 | 0.47 |
| S5_G_0.5_3 | 0.96 | 0.96 | 0.96 | 0.93 | 0.90 | 0.93 | 0.98 | 0.91 | 0.92 | 0.94 | 0.94 | 0.94 | 0.91 | 0.90 | 0.42 | 0.42 | 0.42 | 0.42 | 0.43 | 0.42 | 0.42 | 0.41 | 0.44 | 0.44 | 0.48 |
| S5_G_0.5_3.5 | 0.90 | 0.88 | 0.88 | 0.83 | 0.87 | 0.88 | 0.89 | 0.93 | 0.94 | 0.96 | 0.90 | 0.90 | 0.93 | 0.83 | 0.42 | 0.43 | 0.43 | 0.43 | 0.43 | 0.44 | 0.42 | 0.41 | 0.43 | 0.43 | 0.46 |
| S5_G_0.5_4 | 0.96 | 0.90 | 0.90 | 0.88 | 0.93 | 0.86 | 0.95 | 0.90 | 0.96 | 0.98 | 0.92 | 0.92 | 0.90 | 0.89 | 0.43 | 0.43 | 0.44 | 0.44 | 0.44 | 0.45 | 0.41 | 0.40 | 0.45 | 0.46 | 0.48 |
| S5_G_1_0 | 0.97 | 0.94 | 0.94 | 0.97 | 0.92 | 0.94 | 0.98 | 0.94 | 0.93 | 0.92 | 0.92 | 0.92 | 0.94 | 0.95 | 0.41 | 0.42 | 0.42 | 0.42 | 0.44 | 0.43 | 0.41 | 0.43 | 0.46 | 0.45 | 0.52 |
| S5_G_1_0.5 | 0.92 | 0.93 | 0.93 | 0.95 | 0.87 | 0.93 | 0.94 | 0.92 | 0.92 | 0.87 | 0.87 | 0.87 | 0.92 | 0.94 | 0.42 | 0.43 | 0.43 | 0.43 | 0.45 | 0.44 | 0.42 | 0.44 | 0.47 | 0.46 | 0.52 |
| S5_G_1_1 | 0.97 | 0.94 | 0.94 | 0.94 | 0.92 | 0.94 | 0.95 | 0.97 | 0.97 | 0.92 | 0.92 | 0.92 | 0.97 | 0.95 | 0.41 | 0.42 | 0.42 | 0.42 | 0.44 | 0.43 | 0.41 | 0.43 | 0.46 | 0.45 | 0.50 |
| S5_G_1_1.5 | 1.00 | 0.96 | 0.96 | 0.93 | 0.94 | 0.90 | 0.98 | 0.91 | 0.96 | 0.94 | 0.94 | 0.94 | 0.91 | 0.94 | 0.42 | 0.42 | 0.42 | 0.42 | 0.43 | 0.42 | 0.42 | 0.41 | 0.46 | 0.46 | 0.48 |
| S5_G_1_2 | 0.96 | 1.00 | 1.00 | 0.93 | 0.93 | 0.90 | 0.96 | 0.90 | 0.91 | 0.91 | 0.91 | 0.91 | 0.90 | 0.88 | 0.42 | 0.43 | 0.43 | 0.43 | 0.43 | 0.42 | 0.42 | 0.41 | 0.45 | 0.45 | 0.48 |
| S5_G_1_2.5 | 0.96 | 1.00 | 1.00 | 0.93 | 0.93 | 0.90 | 0.96 | 0.90 | 0.91 | 0.91 | 0.91 | 0.91 | 0.90 | 0.88 | 0.42 | 0.43 | 0.43 | 0.43 | 0.43 | 0.42 | 0.42 | 0.41 | 0.45 | 0.45 | 0.48 |
| S5_G_1_3 | 0.93 | 0.93 | 0.93 | 1.00 | 0.91 | 0.94 | 0.95 | 0.88 | 0.89 | 0.86 | 0.86 | 0.86 | 0.88 | 0.95 | 0.42 | 0.43 | 0.43 | 0.43 | 0.45 | 0.44 | 0.42 | 0.44 | 0.48 | 0.46 | 0.53 |
| S5_G_1_4.5 | 0.94 | 0.93 | 0.93 | 0.91 | 1.00 | 0.84 | 0.93 | 0.89 | 0.94 | 0.90 | 0.90 | 0.90 | 0.89 | 0.83 | 0.42 | 0.43 | 0.43 | 0.43 | 0.43 | 0.42 | 0.42 | 0.41 | 0.47 | 0.47 | 0.49 |
| S5_G_1.5_0 | 0.90 | 0.90 | 0.90 | 0.94 | 0.84 | 1.00 | 0.92 | 0.94 | 0.90 | 0.88 | 0.88 | 0.88 | 0.94 | 0.92 | 0.43 | 0.43 | 0.44 | 0.44 | 0.44 | 0.43 | 0.43 | 0.46 | 0.48 | 0.43 | 0.50 |
| S5_G_1.5_0.5 | 0.98 | 0.96 | 0.96 | 0.95 | 0.93 | 0.92 | 1.00 | 0.91 | 0.95 | 0.94 | 0.94 | 0.94 | 0.91 | 0.93 | 0.42 | 0.42 | 0.42 | 0.42 | 0.43 | 0.42 | 0.42 | 0.41 | 0.45 | 0.46 | 0.49 |
| S5_G_1.5_1.5 | 0.91 | 0.90 | 0.90 | 0.88 | 0.89 | 0.94 | 0.91 | 1.00 | 0.96 | 0.92 | 0.92 | 0.92 | 1.00 | 0.89 | 0.41 | 0.41 | 0.42 | 0.42 | 0.44 | 0.43 | 0.41 | 0.43 | 0.45 | 0.44 | 0.50 |
| S5_G_1.5_2 | 0.96 | 0.91 | 0.91 | 0.89 | 0.94 | 0.90 | 0.95 | 0.96 | 1.00 | 0.94 | 0.94 | 0.94 | 0.96 | 0.90 | 0.42 | 0.42 | 0.42 | 0.42 | 0.43 | 0.42 | 0.42 | 0.41 | 0.46 | 0.46 | 0.48 |
| S5_G_1.5_2.5 | 0.94 | 0.91 | 0.91 | 0.86 | 0.90 | 0.88 | 0.94 | 0.92 | 0.94 | 1.00 | 0.93 | 0.93 | 0.92 | 0.85 | 0.43 | 0.43 | 0.44 | 0.44 | 0.44 | 0.45 | 0.40 | 0.39 | 0.43 | 0.44 | 0.47 |
| S5_G_1.5_3 | 0.94 | 0.91 | 0.91 | 0.86 | 0.90 | 0.88 | 0.94 | 0.92 | 0.94 | 0.93 | 1.00 | 0.93 | 0.92 | 0.85 | 0.40 | 0.41 | 0.41 | 0.41 | 0.44 | 0.41 | 0.43 | 0.44 | 0.46 | 0.44 | 0.47 |
| S5_G_1.5_4 | 0.94 | 0.91 | 0.91 | 0.86 | 0.90 | 0.88 | 0.94 | 0.92 | 0.94 | 0.93 | 0.93 | 1.00 | 0.92 | 0.85 | 0.40 | 0.41 | 0.41 | 0.41 | 0.42 | 0.41 | 0.40 | 0.39 | 0.43 | 0.44 | 0.47 |
| S5_G_1.5_4.5 | 0.91 | 0.90 | 0.90 | 0.88 | 0.89 | 0.94 | 0.91 | 1.00 | 0.96 | 0.92 | 0.92 | 0.92 | 1.00 | 0.89 | 0.41 | 0.41 | 0.42 | 0.42 | 0.44 | 0.43 | 0.41 | 0.43 | 0.45 | 0.44 | 0.50 |
| S5_G_2_1 | 0.94 | 0.88 | 0.88 | 0.95 | 0.83 | 0.92 | 0.93 | 0.89 | 0.90 | 0.85 | 0.85 | 0.85 | 0.89 | 1.00 | 0.42 | 0.43 | 0.43 | 0.43 | 0.45 | 0.44 | 0.42 | 0.45 | 0.49 | 0.47 | 0.52 |
| S5_G_2_3 | 0.42 | 0.42 | 0.42 | 0.42 | 0.42 | 0.43 | 0.42 | 0.41 | 0.42 | 0.43 | 0.40 | 0.40 | 0.41 | 0.42 | 1.00 | 0.96 | 0.96 | 0.96 | 0.64 | 0.68 | 0.57 | 0.60 | 0.62 | 0.68 | 0.65 |
| S5_G_2_3.5 | 0.42 | 0.43 | 0.43 | 0.43 | 0.43 | 0.43 | 0.42 | 0.41 | 0.42 | 0.43 | 0.41 | 0.41 | 0.41 | 0.43 | 0.96 | 1.00 | 0.93 | 0.94 | 0.64 | 0.64 | 0.58 | 0.58 | 0.60 | 0.67 | 0.64 |
| S5_G_2_4 | 0.42 | 0.43 | 0.43 | 0.43 | 0.43 | 0.44 | 0.42 | 0.42 | 0.42 | 0.44 | 0.41 | 0.41 | 0.42 | 0.43 | 0.96 | 0.93 | 1.00 | 0.94 | 0.63 | 0.67 | 0.55 | 0.58 | 0.61 | 0.67 | 0.64 |
| S5_G_2_4.5 | 0.42 | 0.43 | 0.43 | 0.43 | 0.43 | 0.44 | 0.42 | 0.42 | 0.42 | 0.44 | 0.41 | 0.41 | 0.42 | 0.43 | 0.96 | 0.94 | 0.94 | 1.00 | 0.65 | 0.68 | 0.55 | 0.58 | 0.61 | 0.68 | 0.65 |
| S5_G_2.5_0 | 0.43 | 0.43 | 0.43 | 0.45 | 0.43 | 0.44 | 0.43 | 0.44 | 0.43 | 0.44 | 0.44 | 0.42 | 0.44 | 0.45 | 0.64 | 0.64 | 0.63 | 0.65 | 1.00 | 0.70 | 0.69 | 0.70 | 0.71 | 0.69 | 0.65 |
| S5_G_2.5_4 | 0.42 | 0.42 | 0.42 | 0.44 | 0.42 | 0.43 | 0.42 | 0.43 | 0.42 | 0.45 | 0.41 | 0.41 | 0.43 | 0.44 | 0.68 | 0.64 | 0.67 | 0.68 | 0.70 | 1.00 | 0.60 | 0.68 | 0.71 | 0.66 | 0.66 |
| S5_G_3_1 | 0.42 | 0.42 | 0.42 | 0.42 | 0.42 | 0.43 | 0.42 | 0.41 | 0.42 | 0.40 | 0.43 | 0.40 | 0.41 | 0.42 | 0.57 | 0.58 | 0.55 | 0.55 | 0.69 | 0.60 | 1.00 | 0.64 | 0.66 | 0.63 | 0.60 |
| S5_G_3_1.5 | 0.41 | 0.41 | 0.41 | 0.44 | 0.41 | 0.46 | 0.41 | 0.43 | 0.41 | 0.39 | 0.44 | 0.39 | 0.43 | 0.45 | 0.60 | 0.58 | 0.58 | 0.58 | 0.70 | 0.68 | 0.64 | 1.00 | 0.79 | 0.65 | 0.65 |
| S5_G_3_2 | 0.46 | 0.45 | 0.45 | 0.48 | 0.47 | 0.48 | 0.45 | 0.45 | 0.46 | 0.43 | 0.46 | 0.43 | 0.45 | 0.49 | 0.62 | 0.60 | 0.61 | 0.61 | 0.71 | 0.71 | 0.66 | 0.79 | 1.00 | 0.68 | 0.66 |
| S5_G_3_2.5 | 0.46 | 0.45 | 0.45 | 0.46 | 0.47 | 0.43 | 0.46 | 0.44 | 0.46 | 0.44 | 0.44 | 0.44 | 0.44 | 0.47 | 0.68 | 0.67 | 0.67 | 0.68 | 0.69 | 0.66 | 0.63 | 0.65 | 0.68 | 1.00 | 0.91 |
| S5_G_3.5_3.5 | 0.48 | 0.48 | 0.48 | 0.53 | 0.49 | 0.50 | 0.49 | 0.50 | 0.48 | 0.47 | 0.47 | 0.47 | 0.50 | 0.52 | 0.65 | 0.64 | 0.64 | 0.65 | 0.65 | 0.66 | 0.60 | 0.65 | 0.66 | 0.91 | 1.00 |

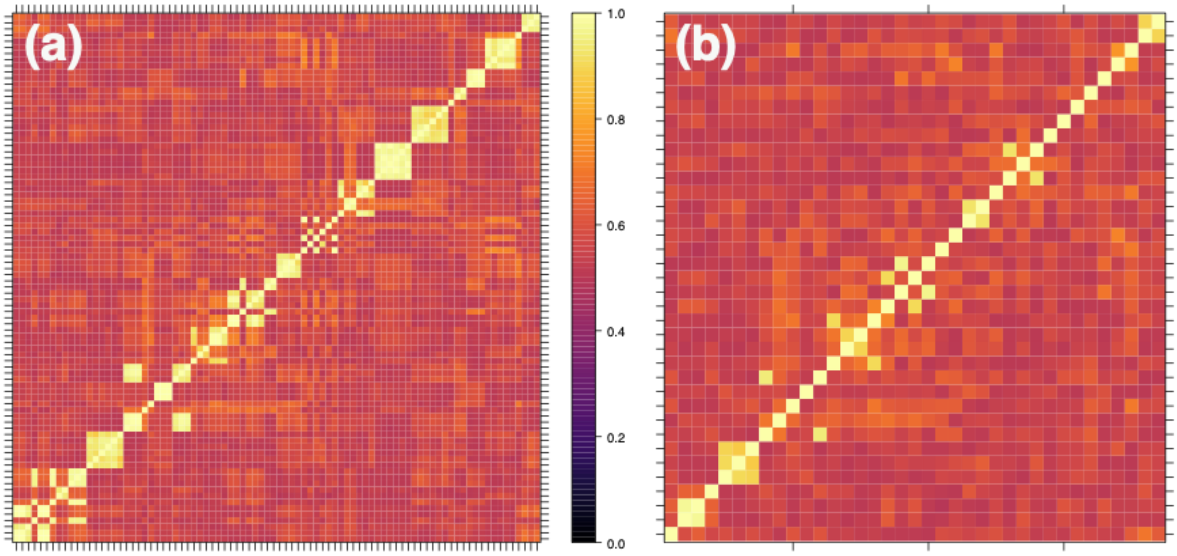


Figure S1. Heat maps displaying the correlation ($r^2$) between all (a) 88 individuals collected and (b) the 30 MLGs kept for species-wide site level analysis.
